# Condensation of pericentrin proteins in human cells illuminates phase separation in centrosome assembly

**DOI:** 10.1101/2020.05.08.084749

**Authors:** Xueer Jiang, Dac Bang Tam Ho, Karan Mahe, Jennielee Mia, Guadalupe Sepulveda, Mark Antkowiak, Linhao Jiang, Soichiro Yamada, Li-En Jao

## Abstract

At the onset of mitosis, centrosomes expand the pericentriolar material (PCM) to maximize their microtubule-organizing activity. This step, termed centrosome maturation, ensures proper spindle organization and faithful chromosome segregation. However, as the centrosome expands, how PCM proteins are recruited and held together without membrane enclosure remains elusive. We found that endogenously expressed pericentrin (PCNT), a conserved PCM scaffold protein, condenses into dynamic granules during late G2/early mitosis before incorporating into mitotic centrosomes. Furthermore, the N-terminal portion of PCNT—enriched with conserved coiled-coils (CCs) and low-complexity regions (LCRs)—phase separates into dynamic condensates that selectively recruit PCM proteins and nucleate microtubules in cells. We propose that CCs and LCRs, two prevalent sequence features in the centrosomal proteome, are preserved under evolutionary pressure in part to mediate liquid-liquid phase separation, a process that bestows upon the centrosome distinct properties critical for its assembly and functions.

## Introduction

The centrosome acts as a major microtubule organizing center (MTOC) in many animal cells and consists of a pair of centrioles embedded in a proteinaceous network of pericentriolar material (PCM) (***Conduit et al., 2015***; ***Rieder and Borisy, 1982***; ***Vorobjev and Chentsov Yu, 1982***; ***Wang et al., 2011***; ***Woodruff et al., 2014***). The MTOC activity of the centrosome is determined by the PCM, which acts as a scaffold to recruit MT regulators and nucleators, such as *γ*-tubulin ring complexes (*γ*-TuRCs) (***Jeng and Stearns, 1999***; ***Moritz et al., 1995b***, ***1998***; ***Oegema et al., 1999***; ***Zheng et al., 1995***). The PCM is not a static structure. In interphase cells, relatively small amounts of PCM are assembled around the centriole and organized as a layered, nanometer-sized toroid (***Fu and Glover, 2012***; ***Lawo et al., 2012***; ***Mennella et al., 2012***; ***Sonnen et al., 2012***). As the cell enters mitosis, the PCM expands dramatically into a micron-sized ensemble, with a concomitant increase in its MTOC activity as the mitotic spindle forms in a process termed centrosome maturation (***Khodjakov and Rieder, 1999***; ***Mahen and Venkitaraman, 2012***; ***Mennella et al., 2014***; ***Palazzo et al., 2000***; ***Piehl et al., 2004***).

Over the past decades, proteins important for centrosome maturation have been identified, and the molecular framework of centrosome maturation has been revealed (***Andersen et al., 2003***; ***Dobbelaere et al., 2008***; ***Goshima et al., 2007***; ***Hamill et al., 2002***; ***Hutchins et al., 2010***; ***Neumann et al., 2010***; ***Sonnichsen et al., 2005***; ***Woodruff et al., 2015***). At the molecular level, centrosome maturation is initiated upon phosphorylation of core PCM components—e.g., spindle-defective protein 5 (SPD-5), centrosomin (Cnn)/CDK5RAP2 (CEP215), and pericentrin (PCNT)—by mitotic kinases such as Polo/polo-like kinase 1 (PLK1) and aurora kinase A (***Barr and Gergely, 2007***; ***Berdnik and Knoblich, 2002***; ***Conduit et al., 2014a***; ***Hannak et al., 2001***; ***Joukov et al., 2014***; ***Kinoshita et al., 2005***; ***Lee and Rhee, 2011***; ***Woodruff et al., 2017***, ***2015***; ***Wueseke et al., 2016***). These events trigger the cooperative assembly of additional PCM proteins (***Alvarez-Rodrigo et al., 2019***; ***Chinen et al., 2021***; ***Conduit et al., 2014b***; ***Hamill et al., 2002***; ***Kemp et al., 2004***; ***Meng et al., 2015***) and *γ*-TuRCs, leading to two mitotic centrosomes with maximized MTOC activities that facilitate bipolar spindle assembly and subsequent chromosome segregation (***Chinen et al., 2021***; ***Conduit et al., 2015***; ***Watanabe et al., 2020***; ***Woodruff et al., 2014***). While the mechanism of centrosome maturation has been elucidated at the molecular level, the biophysical principle of PCM assembly remains elusive at the organellar level—without an enclosing membrane, what keeps the crowded PCM proteins from dispersing?

Liquid-liquid phase separation (LLPS)—a process through which macromolecules de-mix and partition from a single phase into two or more distinct phases in a concentration-dependent manner—has emerged as a mechanism that underlies a variety of cellular processes involving non-membrane-bound compartments or organelles (reviews in ***Banani et al., 2017***; ***Holehouse and Pappu, 2018***; ***Hyman et al., 2014***; ***Shin and Brangwynne, 2017***). Recently, Woodruff and colleagues proposed that the centrosome is formed through LLPS (***Woodruff et al., 2017***). They showed that *in vitro* purified SPD-5, a core PCM protein with extensive coiled-coils in *C. elegans* (***Hamill et al., 2002***), forms spherical liquid “condensates” *in vitro* in the presence of crowding reagents, which mimic the dense cytoplasm (***Woodruff et al., 2017***). SPD-5 condensates possess a centrosome-like activity *in vitro*, capable of nucleating MTs after selectively recruiting tubulin dimers and cognate proteins (ZYG-9 and TPXL-1) (***Woodruff et al., 2017***). These data are also consistent with the mathematical modeling of centrosomes as autocatalytic droplets formed by LLPS (***Zwicker et al., 2014***). However, it is unclear how closely this *in vitro* system reflects centrosomal MT nucleation *in vivo*. As Woodruff and colleagues did not include *γ*-tubulin in their study (***Woodruff et al., 2017***), it also remains to be determined whether SPD-5 condensates can recruit *γ*-tubulin, a critical *in vivo* MT nucleation factor for many species (***Felix et al., 1994***; ***Hannak et al., 2002***; ***Joshi et al., 1992***; ***Oakley et al., 1990***; ***Stearns et al., 1991***; ***Stearns and Kirschner, 1994***; ***Zheng et al., 1991***).

Contrary data suggest that LLPS may not play a role in centrosome assembly. For example, Cnn—a major mitotic PCM component and functional homolog of SPD-5 in *D. melanogaster*—does not undergo dynamic internal rearrangements as it incorporates into the centrosome *in vivo* (***Conduit et al., 2010***, ***2014a***). Two short conserved domains of Cnn can self-assemble into solid-like scaffolds *in vitro*, but no liquid-to-solid phase transition has been observed (***Feng et al., 2017***). However, the action of these Cnn segments in the context of full-length Cnn *in vivo* remains unknown. Together, with the available evidence, it remains elusive whether LLPS underlies centrosome assembly.

In vertebrates, PCNT plays a particularly important role in PCM assembly as it is required for the initiation (***Lee and Rhee, 2011***; ***Zimmerman et al., 2004***) and recruitment of key PCM components during centrosome maturation (***Haren et al., 2009***; ***Lawo et al., 2012***; ***Zimmerman et al., 2004***). We recently showed that PCNT enrichment during centrosome maturation is controlled by a co-translational targeting mechanism that ensures timely production and spatial deposition of PCNT at mitotic centrosomes (***Sepulveda et al., 2018***). Indeed, PCNT expression is tightly regulated. For example, human loss-of-function mutations of PCNT cause microcephalic osteodysplastic primordial dwarfism type II (***Anitha et al., 2009***; ***Delaval and Doxsey, 2010***; ***Griffith et al., 2008***; ***Numata et al., 2009***; ***Rauch et al., 2008***), whereas elevated PCNT levels disrupt ciliary protein trafficking and sonic hedgehog signaling, and may contribute to clinical features of Down syndrome (***Galati et al., 2018***). Despite its importance at the cellular and organismal levels, the precise function of PCNT in centrosome assembly remains enigmatic.

Here we demonstrate that endogenously GFP-tagged human PCNT forms droplet-like granules around centrosomes during late G2/early M phases. These GFP-PCNT granules appear to fuse and split in seconds and are dissolved by several aliphatic alcohols, which disrupt weak hydrophobic interactions between sequences that can promote LLPS (***Lin et al., 2016***). These data suggest that full-length PCNT may undergo LLPS at physiologically relevant conditions during centrosome maturation. We further show that the N-terminal and middle segments of PCNT—enriched with conserved coiled-coils (CCs) and low-complexity regions (LCRs)—undergo LLPS in a concentration-dependent manner with defined phase transitioning boundaries. Similar to dynamic pericentrosomal granules formed by the *in situ*-tagged full-length PCNT, these phase-separated PCNT “condensates” are also sensitive to the same aliphatic alcohol treatment. Furthermore, condensates formed by the middle segment of PCNT transition from liquid-to gel-like states over time and exhibit centrosome-like activities in cells, including selectively recruiting endogenous PCM components and nucleating MTs. Our findings that full-length PCNT condenses into dynamic, aliphatic alcohol-sensitive granules under physiologically relevant conditions, and that the CC- and LCR-rich segments of PCNT undergo concentration-dependent LLPS shed new light on the process of LLPS and the role of CCs and LCRs—two sequence features abundant in centrosome proteome—in centrosome assembly.

## Results

### Endogenously GFP-tagged PCNT forms dynamic, aliphatic alcohol-sensitive pericentrosomal granules during late G2/early M phases

To study full-length PCNT at endogenous levels in cells, we used the CRISPR technology (***Lin et al., 2014a***; ***Zhang et al., 2017***) to insert super-folder GFP sequence to the 5’ end of the *PCNT* locus in hTERT-immortalized human retinal pigment epithelial (RPE-1) cells (***Figure 1–Figure Supplement 1***). The GFP tagging at the N-terminus of PCNT did not affect cell morphology, mitotic progression, or the recruitment of another major PCM protein CDK5RAP2/CEP215 to mitotic centrosomes (***Figure 1–Figure Supplement 1***). As expected, GFP-PCNT decorated centrosomes, but upon close examination, it also formed small droplet-like granules—generally smaller than 400 nm in diameter—near centrosomes (***Figure 1A***). These pericentrosomal PCNT granules were observed predominantly during late G2/early M phases, concomitant with the process of centrosome maturation (***Figure 1B***, ***Figure 1–Figure Supplement 2***). Similar pericentrosomal PCNT granules were also observed by immunostaining endogenous, untagged PCNT during early mitosis (***Figure 1–Figure Supplement 3***). These pericentrosomal PCNT granules were highly dynamic; they appeared to fuse and split over a timescale of seconds (***Figure 1A***, ***Figure 1–video 1***, ***Figure 1–video 2***). Similar dynamic PCNT granules were also observed in another independent GFP knock-in clone (***Figure 1–video 3***). However, because the size of these granules is close to the diffraction limit of light, we could not use light microscopy to unequivocally measure their aspect ratios and determine whether they indeed have a spherical shape, a feature that would suggest a liquid form.

**Figure 1.**
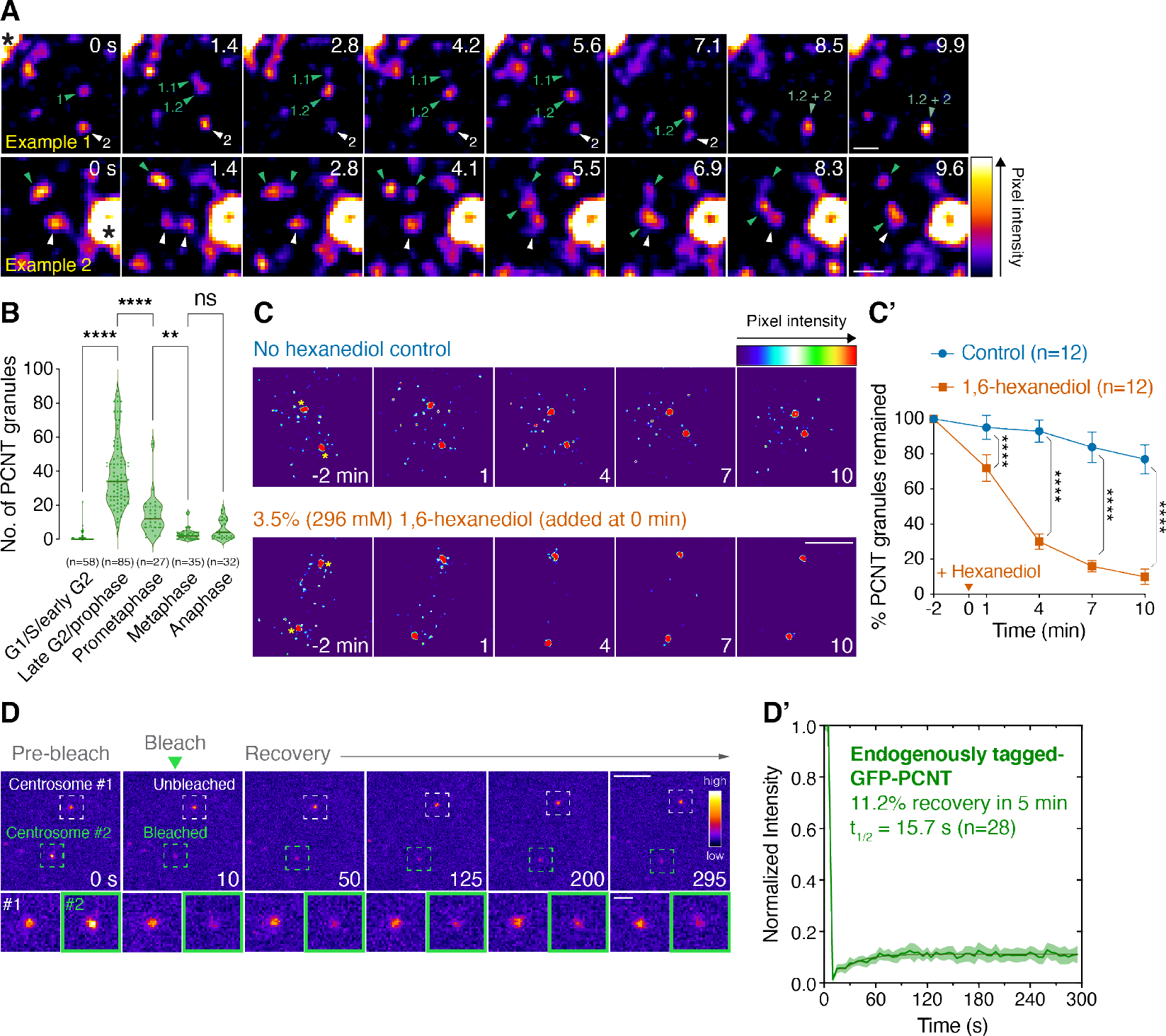
Endogenously GFP-tagged PCNT forms dynamic, aliphatic alcohol-sensitive pericentrosomal granules during late G2/early M phases before incorporating into a largely non-dynamic mitotic PCM. (**A**) Time-lapse micrographs of GFP-PCNT expressed from its endogenous locus during late G2/early M phases. Arrowheads denote the fusing and splitting events of the GFP-PCNT granules. Asterisks in time 0 denote the centrosomes. Similar results were obtained from more than three biological replicates (also see ***Figure 1–video 1***, ***Figure 1–video 2***, and ***Figure 1–video 3***). (**B**) Quantification of PCNT granule numbers at different cell cycle stages. Data are median ± the first and third quartiles. n, number of cells analyzed from more than three biological replicates. Representative images are shown in ***Figure 1–Figure Supplement 2***. (**C**) Time-lapse micrographs of pericentrosomal GFP-PCNT granules without or with the acute 3.5%/296 mM 1,6-hexanediol treatment. Time 0 is set at the time of hexanediol addition. Asterisks in time −2 min denote the centrosomes. (**C’**) Quantification of the results shown in (C). Percent of GFP-PCNT granules remained, without and with the acute 1,6-hexanediol treatment, as the function of time was plotted and represented as mean ± 95% CI from three biological replicates. The total number of cells analyzed for each condition is indicated. The arrowhead denotes the time of hexanediol addition (time 0). (**D**) FRAP analysis of endogenously GFP-tagged PCNT in RPE-1 cells during late G2/early mitosis. Only one of the centrosomes (Centrosome #2) was photobleached, and the fluorescence recovery was recorded every 5 s for 5 min. Dashed squares delineate centrosomes. (**D’**) The percent recovery and half-life (t_1/2_) after photobleaching were calculated after fitting the data with non-linear regression. Data are mean ± 95% CI. n, number of centrosomes analyzed from two biological replicates. p-values were determined by one-way ANOVA (B) or the Student’s t-test (two-tailed) (C’). **: p<0.01, ****: p<0.0001; ns, not significant. Scale bars, 0.5 µm (A), 5 µm (C), 10 µm (D), and 2 µm (D, inset). **Figure 1–Figure supplement 1.** Generation and validation of GFP-PCNT knock-in cells. **Figure 1–Figure supplement 2.** PCNT granules at different cell cycle stages. **Figure 1–Figure supplement 3.** Endogenous pericentrosomal PCNT granules at early mitosis. **Figure 1–Figure supplement 4.** Effects of different aliphatic alcohols on PCNT granules. **Figure 1–Figure supplement 5.** Validation of *TP53* knockout RPE-1 cells. **Figure 1–video 1.** PCNT granules-Example 1. **Figure 1–video 2.** PCNT granules-Example 2. **Figure 1–video 3.** PCNT granules-Another knock-in clone.

To probe the biophysical properties of these dynamic PCNT granules, we turned to aliphatic alcohol 1,6-hexanediol. 1,6-hexanediol was originally shown to weaken the permeability barrier of nuclear pore complexes (NPCs) by disrupting weak hydrophobic interactions between phenylalanine-glycine (FG) repeats of nucleoporins (***Patel et al., 2007***; ***Schmidt and Gorlich, 2015***; ***Shulga and Goldfarb, 2003***). 1,6-hexanediol is also shown to dissolve several membraneless, liquid-like cellular assemblies such as RNA-protein granules (e.g., P granules and stress granules) (***Kroschwald et al., 2015***; ***Updike et al., 2011***). Furthermore, the liquidity of these cellular assemblies is linked to their sensitivity to 1,6-hexanediol (***Kroschwald et al., 2015***, ***2017***; ***Lin et al., 2016***). This correlation has been attributed to the ability of 1,6-hexanediol to disrupt weak hydrophobic interactions through the hydrophobic effect exerted by its alkyl chain—as in the case for disrupting the NPC permeability barrier (***Patel et al., 2007***; ***Schmidt and Gorlich, 2015***; ***Shulga and Goldfarb, 2003***)—as well as the ability of 1,6-hexanediol to reduce aqueous surface tension (***Romero et al., 2007***).

Based on these previous studies, we treated the cells with 1,6-hexanediol acutely and immediately followed the fate of pericentrosomal PCNT granules by time-lapse microscopy. We found that PCNT granules were dissolved within minutes, whereas PCNT assembly at mitotic centrosomes was refractory to the treatment (***Figure 1C, C’***). These results suggest that the stability of PCNT granules is likely maintained by weak hydrophobic interactions, and that PCNT may change its modes of interaction and/or biophysical properties upon incorporation into mitotic centrosomes—e.g., through the anchoring of PCNT near the centriolar wall via its C-terminal PACT motif (***Gillingham and Munro, 2000***; ***Takahashi et al., 2002***).

We next asked whether other aliphatic alcohols also exert similar effects on PCNT granules. We found that similar to 1,6-hexanediol, several aliphatic alcohols—varying in the length of the alkyl chain and the position of hydroxyl groups—also dissolved PCNT granules, but not the PCNT assembly at centrosomes. Furthermore, the effectiveness of the aliphatic alcohol on dissolving PCNT granules was roughly proportional to its relative hydrophobicity (***Figure 1–Figure Supplement 4***), a characteristic also observed when the NPC permeability barrier and other liquid-like, membraneless cellular assemblies were exposed to various aliphatic alcohols (***Lin et al., 2016***; ***Patel et al., 2007***; ***Ribbeck and Gorlich, 2002***; ***Rog et al., 2017***; ***Schmidt and Gorlich, 2015***; ***Shulga and Goldfarb, 2003***; ***Updike et al., 2011***).

We also probed the biophysical properties of the PCNT assembly at mitotic centrosomes by a fluorescence recovery after photobleaching (FRAP) experiment, in which a bleaching laser targeted the mitotic centrosome and the recovery of fluorescence was measured to quantify exchange of PCNT between the cytoplasm and the PCM. A limited fluorescence recovery of GFP-PCNT at the mitotic centrosome after photobleaching was observed (∼11% in 5 min, ***Figure 1D, D’***), indicating that there was little exchange of PCNT at the mitotic centrosome, and/or there was a limited amount of non-centrosomal PCNT available for exchange. Thus, consistent with the aliphatic alcohol data, this FRAP result suggests that the PCNT assembly at mitotic centrosomes is largely non-dynamic, in contrast to the nearby dynamic pericentrosomal PCNT granules. Unfortunately, due to the highly dynamic nature and the small size of pericentrosomal PCNT granules, we were unable to perform similar FRAP experiments on these PCNT granules to probe their biophysical properties.

Taken together, results from these experiments suggest that under physiologically relevant conditions and predominantly during late G2/early M phases when the centrosome is maturing, PCNT condenses into dynamic, aliphatic alcohol-sensitive granules, likely through weak hydrophobic interactions between PCNT molecules. However, once PCNT is incorporated into mitotic centrosomes, the modes of interaction and/or biophysical properties of PCNT change through a yet unknown mechanism, making PCNT assembly at the PCM largely non-dynamic and resistant to the dissolution of aliphatic alcohols.

### Coiled-coils and low-complexity regions of pericentrin are more conserved than the rest of the protein sequence

If condensation of PCNT is of evolutionary significance, the sequence features that contribute to condensation should be conserved across species. To test this hypothesis, we constructed an alignment of 169 pericentrin orthologous proteins—167 from vertebrates, one each from fruit fly (D-PLP) (***Martinez-Campos et al., 2004***) and budding yeast (Spc110) (***Knop and Schiebel, 1997***; ***Sundberg and Davis, 1997***) (***Figure 2A*** and ***Figure 2–Figure Supplement 1***). The analysis shows that a number of regions are highly conserved. One is around the C-terminus, particularly at the centrosomal anchoring PACT motif (***Gillingham and Munro, 2000***; ***Takahashi et al., 2002***). Another conserved region is in the middle portion of the protein. In contrast, the N-terminus is not well conserved, with the evidence of clade-specific insertions.

**Figure 2.**
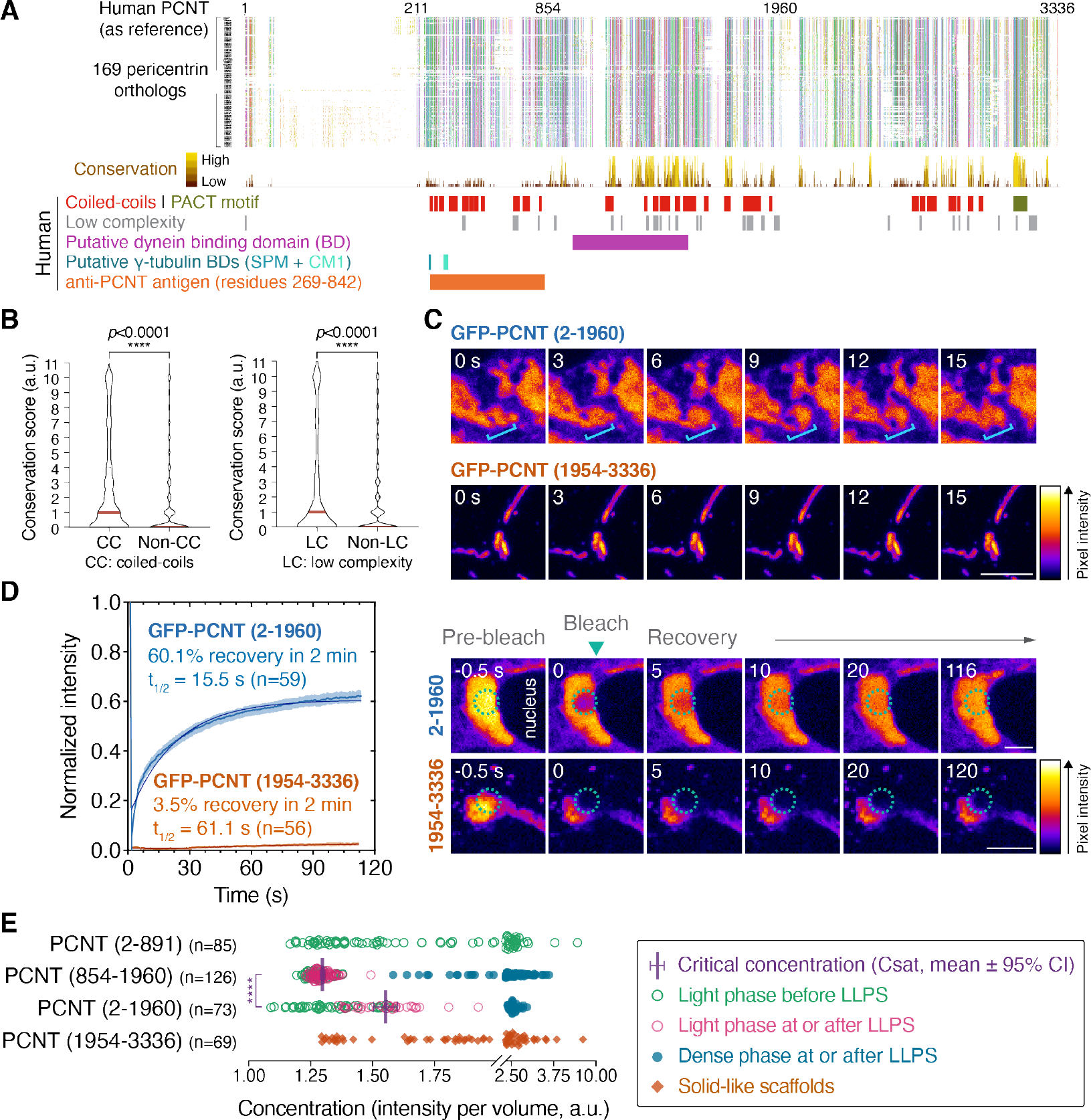
N-terminal segments of PCNT phase separate in a concentration-dependent manner in cells. **(A)** Alignments of 169 pericentrin orthologous proteins from vertebrates (167), fruit fly (1), and budding yeast (1), colored by the Clustal X color scheme in Jalview (***Figure 2–Figure Supplement 1***). Conservation scores, locations of the predicted coiled-coils (CCs), PACT motif, low-complexity (LC) regions, putative dynein and *y*-tubulin binding domains (BDs) of human PCNT, and the epitopes of the anti-PCNT antibody (Abcam, ab4448) are noted below the alignments. (**B**) Conservation scores within or outside of CC or LC regions of human PCNT. Data are median with the third quartile. (**C**) Representative time-lapse images of GFP-PCNT (2-1960) condensates and GFP-PCNT (1954-3336) scaffolds 24 h post Dox induction in RPE-1 cells. Brackets denote an area with dynamic rearrangement of GFP-PCNT (2-1960) condensates; PCNT (1954-3336) scaffolds are non-dynamic (also see ***Figure 2–video 1*** and ***Figure 2–video 2***). (**D**) FRAP analyses of GFP-PCNT (2-1960) condensates and GFP-PCNT (1954-3336) scaffolds in RPE-1 cells. Dashed circles denote the bleached sites. Data are mean ± 95% CI. n, number of condensates/scaffolds analyzed from more than three biological replicates. The percent recovery and half-life (t_1/2_) after photobleaching were calculated after fitting the data with non-linear regression. (**E**) Quantification of relative protein concentrations in live cells expressing various GFP-tagged PCNT segments after Dox induction (see ***Figure 2–Figure Supplement 3*** for details and representative images). PCNT (2-1960) and PCNT (854-1960) phase separated after reaching their respective critical concentrations (Csats), the concentrations of the light phase at which LLPS just occurred. Csats are mean ± 95% CI. n, number of cells analyzed from three biological replicates. The p-values were determined by the Student’s t-test (two-tailed). ****: p<0.0001. a.u., arbitrary unit. Scale bars, 5 µm. **Figure 2–Figure supplement 1.** Color scheme for multiple alignments. **Figure 2–Figure supplement 2.** Disorder predictions of human PCNT. **Figure 2–Figure supplement 3.** Construction of protein phase diagrams in cells. **Figure 2–video 1.** Dynamic GFP-PCNT (2-1960) condensates. **Figure 2–video 2.** Non-dynamic GFP-PCNT (1954-3336) scaffolds.

To gain insights into the properties of these conserved sequences, we did further *in silico* analyses, focusing on human PCNT (***Figure 2A***). The Ncoils (***Lupas et al., 1991***) and SEG (***Wootton, 1994***) programs respectively predict that human PCNT is enriched with coiled-coils (CCs) and low-complexity regions (LCRs)—which often overlap with intrinsically disordered sequences, a sequence feature that can mediate multivalent interactions to drive LLPS (***Boke et al., 2016***; ***Kato et al., 2012***; ***Lin et al., 2015***; ***Molliex et al., 2015***; ***Patel et al., 2015***; ***Riback et al., 2017***). Indeed some disorder predictors (e.g., PONDR, ***Peng et al., 2005***, ***2006***) predicted that human PCNT is largely disordered except for the C-terminal PACT motif (***Figure 2–Figure Supplement 2***). However, as a known limitation with current disorder predictions (***Atkins et al., 2015***), not all disorder predictors are in complete agreement, with each predictor suggesting different degrees of disorder/order tendency (e.g., IUpred, ***Dosztanyi et al., 2005***, predicted an overall more ordered structure than PONDR did). Statistical analyses further showed that CCs and LCRs in human PCNT are significantly more conserved than non-CCs and non-LCRs (***Figure 2B***). Together, these results suggest that CCs and LCRs across pericentrin orthologous proteins are likely under natural selection to preserve their molecular functions.

### The N-terminal, CC/LCR-rich segments of PCNT undergo LLPS in a concentration-dependent manner

Given that both CCs and LCRs can drive LLPS (*Berry et al., 2015*; *Boeynaems et al., 2018*; *Elbaum-Garinkle et al., 2015*; *Fang et al., 2019*; *Hennig et al., 2015*; *Lu et al., 2020*; *Molliex et al., 2015*; *Nott et al., 2015*; *Rog et al., 2017*; *Smith et al., 2016*; *Wang et al., 2018*; *Wippich et al., 2013*; *Zeng et al., 2016*; *Zhang et al., 2018*), we hypothesized that the CC/LCR-rich sequences drive LLPS of the full-length PCNT to form dynamic pericentrosomal PCNT granules observed in the knock-in cells. To test this hypothesis, control PCNT transcription tightly, and map LLPS determinants, we expressed GFP-tagged N- or C-terminal segment of human PCNT under the control of a doxycycline (Dox)-inducible promoter. We stably integrated each construct in RPE-1 cells using a *piggyBac* transposon system, which is free from limitations on insert size (*Kim et al., 2016*). Upon Dox induction, live cell imaging showed that the N-terminal segment, GFP-PCNT (2-1960), formed dynamic condensates (*Figure 2C* and *Figure 2–video 1*) with fast internal rearrangement of molecules as determined by FRAP (*Figure 2D*). In contrast, the C-terminal segment, GFP-PCNT (1954-3336), formed solid-like scaffolds with little internal rearrangement (*Figure 2C* and *Figure 2*–*video 2*) or fluorescence recovery after photobleaching (*Figure 2D*).

To further map the sequences that drive LLPS, we tested GFP-tagged PCNT (2-891) and (854-1960) constructs, which subdivide PCNT (2-1960) but do not disrupt individual CCs or LCRs. After inducing their expressions, we compared their critical concentrations, the point above which LLPS occurs (Csat) (***Asherie, 2004***). To quantitatively assess the Csat in live cells, we developed an imaging and quantification strategy to measure relative protein concentrations by fluorescence intensity per volume after 3D reconstruction (***Figure 2–Figure Supplement 3***). We found that GFP-PCNT (2-891) remained diffuse in cells as its concentration increased. However, over the same concentration range, GFP-PCNT (854-1960) suddenly formed droplet-like condensates when it reached its Csat for LLPS (***Figure 2E***, ***Figure 2–Figure Supplement 3***, and ***Figure 3***). We also validated the LLPS behavior of GFP-PCNT (2-1960)—which had a slightly higher Csat than GFP-PCNT (854-1960)—and the lack of LLPS for GFP-PCNT (1954-3336) (***Figure 2E*** and ***Figure 2–Figure Supplement 3***). Importantly, since FLAG- and mScarlet-i-tagged PCNT (854-1960) also formed similar condensates (***Figure 3–Figure Supplement 1*** and ***Figure 3–video 1***), GFP tagging did not artifactually drive LLPS. Collectively, these results suggest that the abundant CCs and LCRs within PCNT (854-1960)—which are well conserved across species (***Figure 2A, B***)—contain the key sequence elements that drive the LLPS of PCNT (2-1960) and PCNT (854-1960) segments.

**Figure 3.**
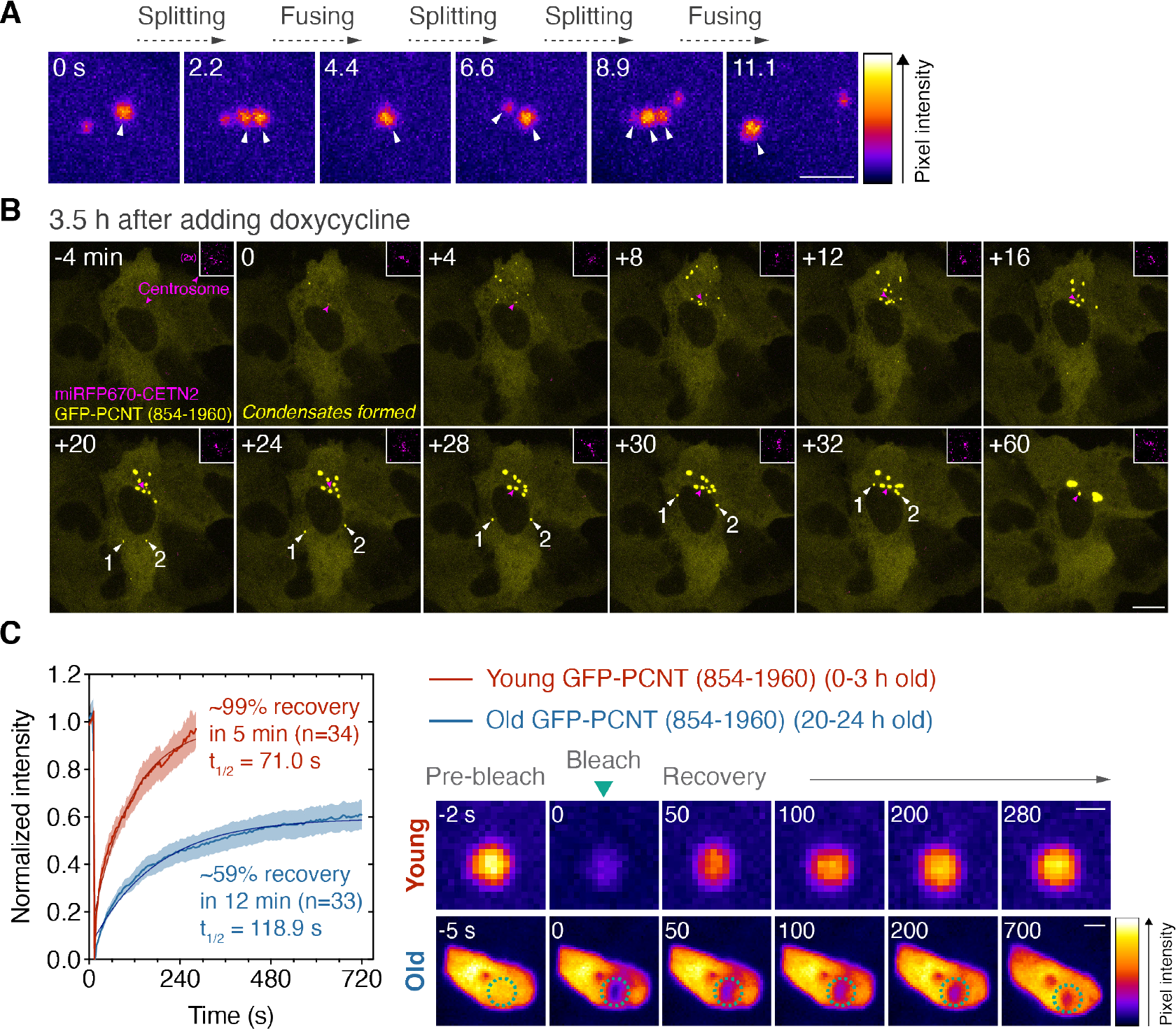
GFP-PCNT (854-1960) undergoes LLPS, coalesces, and moves toward the centrosome. **(A)** Time-lapse micrographs of GFP-PCNT (854-1960) condensates in RPE-1 cells. Arrowheads denote the fast fusing and splitting events of the spherical condensates. Similar results were obtained from three biological replicates. (**B**) Time-lapse micrographs of GFP-PCNT (854-1960) expressed in RPE-1 cells stably expressing miRFP670-CETN2 (insets; magenta arrowheads denote the centrosomes). Time-lapse imaging started 3.5 h post Dox induction. The time when the first condensates formed is marked as time 0. White arrowheads denote the examples of two condensates moving around the nucleus toward the centrosome. Similar results were obtained from more than three biological replicates (also see ***Figure 3–video 1*** and ***Figure 3–video 2***). (**C**) FRAP analyses of different ages of GFP-PCNT (854-1960) condensates in RPE-1 cells. Dashed circles delineate the bleached sites. Data are mean ± 95% CI. n, number of condensates analyzed from three biological replicates. The percent recovery and half-life (t_1/2_) after photobleaching were calculated after fitting the data with non-linear regression. Note that the highly mobile nature of young condensates prevented us from tracking the same condensates consistently beyond 5 min in the recovery phase of the FRAP assay. Scale bars, 2 µm (A), 10 µm (B), 1 µm (C, young condensates), and 2 µm (C, old condensates). **Figure 3–Figure supplement 1.** FLAG- and mScarlet-i-tagged PCNT (854-1960) fusion proteins form condensates. **Figure 3–Figure supplement 2.** Effects of 1,6-hexanediol on different condensates/scaffolds. **Figure 3–video 1.** mScarlet-i-PCNT (854-1960) undergoes LLPS. **Figure 3–video 2.** GFP-PCNT (854-1960) condensates coalesce and move toward the centrosome.

### The liquidity of different PCNT assemblies is correlated with their sensitivity to 1,6-hexanediol

By far we observed that two PCNT fragments (residues 2-1960 and 854-1960) undergo typical LLPS in a concentration-dependent manner to form liquid-like condensates with defined phase transitioning boundaries—at which the Csat can be determined (***Figure 2E***)—whereas PCNT (1954-3336) forms solid-like scaffolds in a wide range of concentrations without discernible phase transition. As the liquidity of membraneless assemblies is linked to their sensitivity to aliphatic alcohols (***Kroschwald et al., 2015***, ***2017***; ***Lin et al., 2016***), we tested how these three assemblies would react when exposed to aliphatic alcohols by live microscopy. Upon adding 3.5% 1,6-hexanediol—the same treatment shown to dissolve pericentrosomal PCNT granules (***Figure 1C, C’***)—PCNT (2-1960) and (854-1960) condensates were also dissolved in minutes, whereas PCNT (1954-3336) scaffolds were not affected (***Figure 3–Figure Supplement 2***). Therefore, the pericentrosomal PCNT granules—formed by the *in situ*-tagged GFP-PCNT—as well as the phase-separated PCNT (2-1960) and (854-1960) condensates are all sensitive to aliphatic alcohols in a similar manner. These results suggest that all these three cellular assemblies are liquid-like, and that the pericentrosomal PCNT granules may also be formed via LLPS, as in the case with PCNT (2-1960) and PCNT (854-1960) condensates.

### GFP-PCNT (854-1960) undergoes LLPS, coalesces, and moves toward the centrosome in a dynein- and MT-dependent manner

Besides phase separating at a lower concentration than GFP-PCNT (2-1960) (***Figure 2E***), GFP-PCNT (854-1960) condensates also exhibited different morphology and behaviors. In particular, early-stage GFP-PCNT (854-1960) condensates formed well-defined spherical, liquid-like droplets as they rapidly split and fused within seconds (***Figure 3A***). Over time, these GFP-PCNT (854-1960) condensates coalesced and converged at the centrosome—arcing around the nucleus in some cases—to form large pericentrosomal condensates (***Figure 3B*** and ***Figure 3–video 2***). Because PCNT (854-1960) contains the putative dynein-binding domain (***Tynan et al., 2000***) (***Figure 2A***), we tested whether the movement of PCNT (854-1960) condensates toward the centrosome is a dynein- and MT-dependent process. We treated the cells with dynein inhibitors (ciliobrevin D and dynarrestin) (***Firestone et al., 2012***; ***Hoing et al., 2018***) or nocodazole after Dox-induced condensate formation and followed condensate movement by time-lapse microscopy. To quantitatively assess the effects of these treatments, we developed a Python program to semi-automatically track and calculate each condensate’s size, number, and distance to the centrosome (miRFP670-CETN2 labeled) at single-cell resolution over time (***Figure 4***–***source code 1***). When dynein was inhibited or MTs were depolymerized, fusion of the condensates—indicated by size increase and number decrease—was impaired (***Figure 4A, B***), and their movements toward the centrosome were also significantly attenuated (***Figure 4C***). Moreover, initial LLPS also occurred closer to the centrosome in the DMSO-than in the dynein inhibitor- or nocodazole-treated cells (***Figure 4D***). Therefore, we conclude that GFP-PCNT (854-1960) and the condensates it forms move toward the centrosome in a dynein- and MT-dependent manner. As LLPS of GFP-PCNT (854-1960) takes place closer to the centrosome with intact dynein activity and MTs, this dynein- and MT-dependent transport could potentially also facilitate LLPS by concentrating and converging GFP-PCNT (854-1960) toward the centrosome along MT tracks.

**Figure 4.**
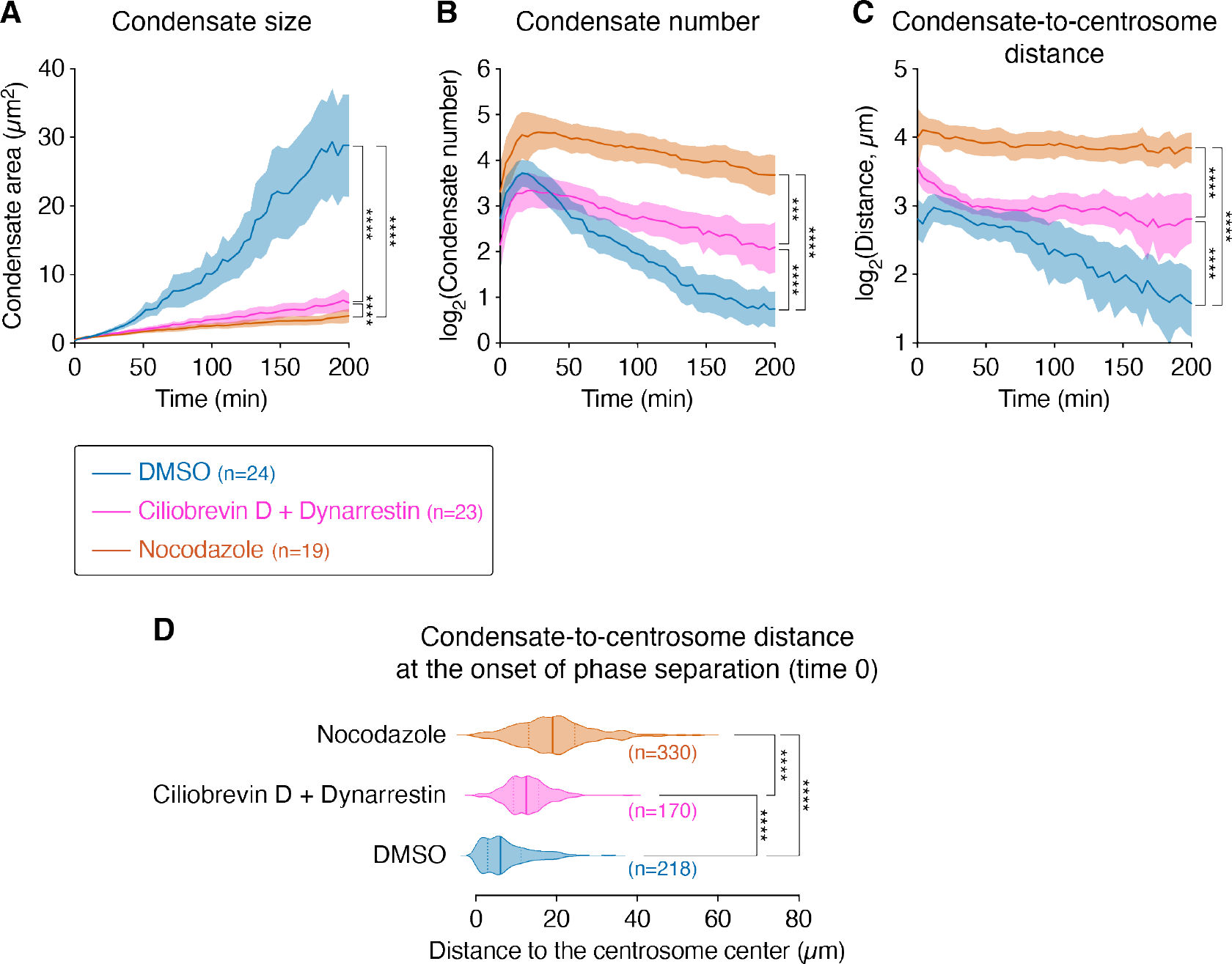
GFP-PCNT (854-1960) condensates coalesce and move toward the centrosome in a dynein- and MT-dependent manner. Quantification of the size (**A**), number (**B**), distance to the centrosome (**C**) and distance to the centrosome at time 0 (the start of phase separation) (**D**) of GFP-PCNT (854-1960) condensates over time in the cells treated with DMSO vehicle, ciliobrevin D and dynarrestin mix (50 µM each), or nocodazole (8.3 µM) (see ***Figure 4–source code 1*** for the Python script). GFP-PCNT (854-1960) was expressed from a Dox-inducible promoter in RPE-1 cells. Data were aligned at the onset of phase separation (time 0) of individual cells. Note that the fusing of PCNT (854-1960) condensates and their movement toward the centrosome were attenuated upon dynein inhibition or MT depolymerization. Initiation of phase separation also occurred closer to the centrosome with intact dynein activity and MTs (D). Data are mean ± 95% CI (A-C) or median ± the first and third quartiles (D). n, number of cells (A-C) or condensates (D) analyzed, from three (DMSO and dynein inhibition) or two (nocodazole) biological replicates. p-values were determined by the F-test that compares the slopes of fitted lines between data sets via linear regression (A-C) or by one-way ANOVA for the distance comparison at time 0 in (D). ***: p=0.0001, ****: p<0.0001. **Figure 4–source code 1.** The Python source code for tracking condensates.

### GFP-PCNT (854-1960) condensates transition from liquid-to gel-like states over time

Close examination of time-lapse data revealed that the rate of fusing and splitting decreased as PCNT (854-1960) condensates coalesced (***Figure 3B***, ***Figure 3***–***video 1***, and ***Figure 3***–***video 2***), suggesting that the “liquidity” of PCNT (854-1960) condensates decreased over time. To test this hypothesis, we used the Dox-inducible system to induce, track, and analyze young (0-3 h old) and old (20-24 h old) GFP-PCNT (854-1960) condensates by FRAP. We found that young condensates recovered fluorescence almost twice as fast as the old ones (***Figure 3C***). Some young condensates recovered 100% of their initial fluorescence intensity because they grew in size. These results suggest that GFP-PCNT (854-1960) condensates become “hardened” over time. Such molecular aging has also been reported for other proteins that phase separate *in vitro*, such as SPD-5 (***Woodruff et al., 2017***), FUS (***Patel et al., 2015***), hnRNPA1 (***Lin et al., 2015***), and Tau (***Wegmann et al., 2018***).

### GFP-PCNT (854-1960) condensates selectively recruit endogenous PCM components

Because the “hardened” SPD-5 condensates recruit tubulins and factors involved in MT nucleation *in vitro* (***Woodruff et al., 2017***), we tested whether GFP-PCNT (854-1960) condensates can also recruit PCM components, including structural (e.g., CEP215) and “client” proteins (e.g., dynein and PLK1). We found that endogenous PCNT, *γ*-tubulin, CEP215, CEP192, dynein intermediate chains (ICs), and PLK1 were significantly enriched in GFP-PCNT (854-1960) condensates, whereas the non-PCM component ribosomal protein S6 (RPS6) was excluded (***Figure 5***). Note that the antibody used to detect endogenous PCNT recognizes the epitopes before residue 854 (***Figure 2A***). Thus, this anti-body will not recognize PCNT (854-1960). Interestingly, these recruited proteins were not uniformly distributed in the condensate, with reticular patterns that resemble mitotic PCM (***Lawo et al., 2012***; ***Sonnen et al., 2012***). To exclude the possibility that any phase-separated condensates could recruit PCM components, we examined the enrichment of PCM proteins in the condensates formed by HO-Tags, the *de novo*-designed homo-oligomeric coiled-coils (***Grigoryan et al., 2011***; ***Huang et al., 2014***; ***Thomson et al., 2014***), which phase separate through multivalent interactions (***Zhang et al., 2018***). We found that PCM components *γ*-tubulin, PCNT, and CEP192 were not enriched in the HOTag condensates (***Figure 5–Figure Supplement 1***).

**Figure 5.**
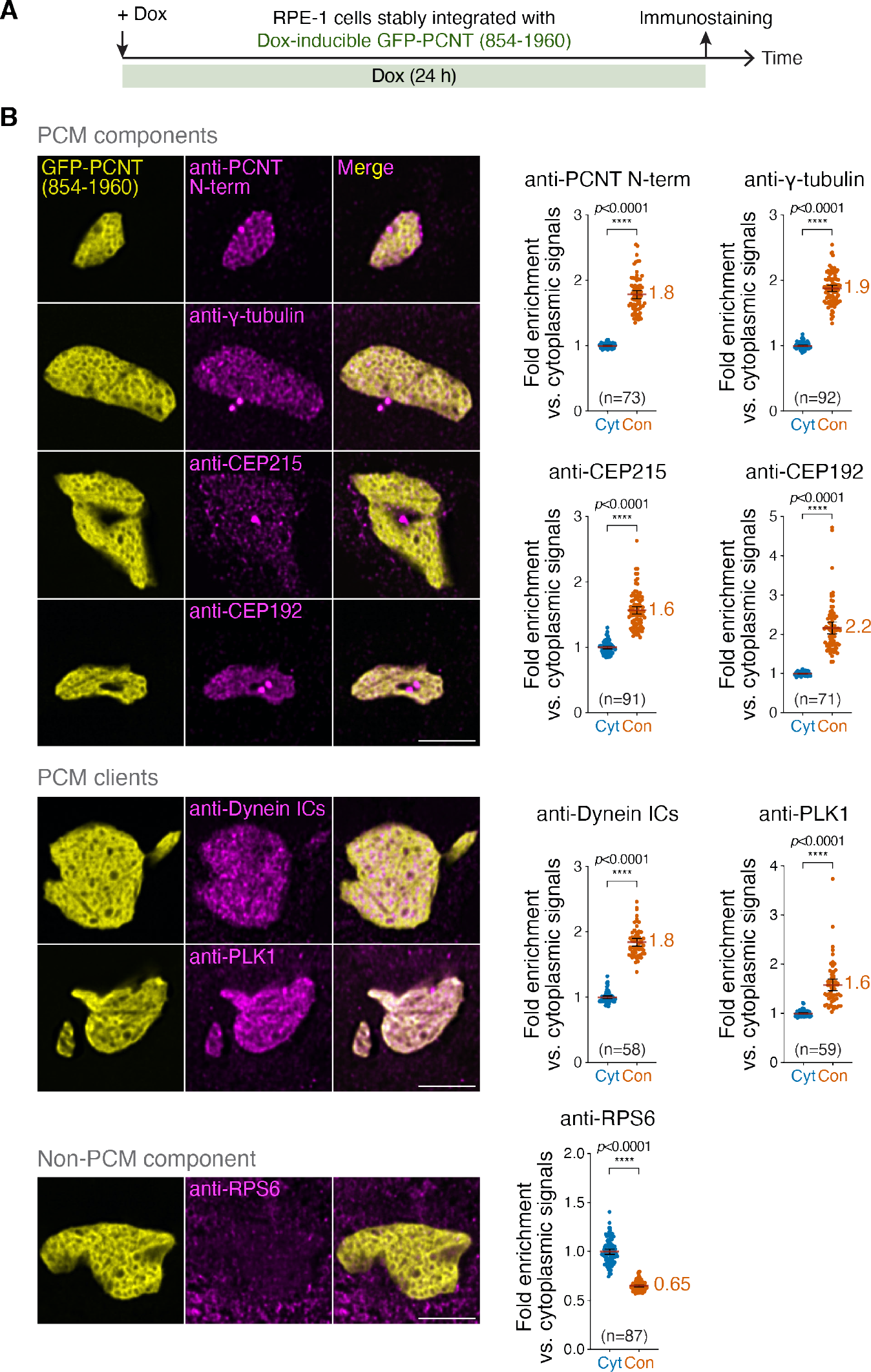
GFP-PCNT (854-1960) condensates selectively recruit endogenous PCM components and clients. (**A**) Schematic of the recruitment assay to show the timeline of Dox induction and immunostaining. Immunofluorescence of PCNT N-terminus (PCNT N-term), *y*-tubulin, CEP215, CEP192, cytoplasmic dynein intermediate chains (Dynein ICs), PLK1, or ribosomal protein S6 (RPS6) in RPE-1 cells after Dox induction to form GFP-PCNT (854-1960) condensates. Fold enrichment of fluorescence signals in the condensate (Con) relative to those in the cytoplasm (Cyt) was quantified. Data are mean ± 95% CI. The mean of fold enrichment is noted. n, number of cells analyzed from at least two biological replicates per protein. p-values were determined by the Student’s t-test (two-tailed). Scale bars, 5 µm. **Figure 5–Figure supplement 1.** Endogenous PCM proteins are not enriched in the HOTag condensates. **Figure 5–Figure supplement 2.** Generation of *PCNT* knockout cells. **Figure 5–Figure supplement 3.** Differential requirements of endogenous PCNT for PCNT (854-1960) condensates to recruit PCM components and clients.

Because PCNT (854-1960) condensates also recruited endogenous full-length PCNT (***Figure 5***), it raised the question of whether the recruitment of other PCM components is mediated through endogenous PCNT. To test this, we repeated the recruitment assays in the presence or absence of endogenous PCNT (i.e., between the parental and *PCNT* knockout cells, ***Figure 5–Figure Supplement 2***). We found that the recruitment of PCM components to PCNT (854-1960) condensates may or may not depend on endogenous PCNT. For example, the recruitment of CEP215 was strictly dependent on endogenous PCNT (***Figure 5–Figure Supplement 3***, ***Group I***), whereas the recruitment of dynein ICs and PLK1 was not (***Figure 5–Figure Supplement 3***, ***Group II***). For *γ*-tubulin and CEP192, endogenous PCNT was not required but facilitated their recruitment to PCNT (854-1960) condensates (***Figure 5–Figure Supplement 3***, ***Group III***). Taken together, these results indicate that PCNT (854-1960) condensates possess unique properties that enable them to selectively recruit endogenous PCM proteins and clients, including endogenous PCNT—which is responsible for the recruitment of some, but not all, PCM proteins to PCNT (854-1960) condensates.

### GFP-PCNT (854-1960) condensates nucleate MTs in cells

Because PCNT (854-1960) condensates recruit *γ*-tubulin (*Figure 5*), the protein critical for MT nucleation (*Felix et al., 1994*; *Hannak et al., 2002*; *Joshi et al., 1992*; *Oakley et al., 1990*; *Stearns et al., 1991*; *Stearns and Kirschner, 1994*; *Zheng et al., 1991*), we tested whether these condensates can also nucleate MTs by MT renucleation assays (*Jao et al., 2017*; *Sanders et al., 2017*). In the MT renucleation assay, we depolymerized MTs by nocodazole, washed out the drug, and monitored MT renucleation by anti-*α*-tubulin immunostaining. We found that MTs were renucleated not only from the centrosome as expected, but also from the interior and surface of PCNT (854-1960) condensates (*Figure 6B*, *arrows*). The condensates also recruited endogenous PCNT as observed before (*Figure 6B* and *Figure 5*). Some small PCNT (854-1960) condensates also recruited endogenous PCNT and nucleated MTs (*Figure 6C*, *asterisks in insets*). Quantification of *α*-tubulin density in condensates and their surrounding cytoplasm confirmed that PCNT (854-1960) condensates had a significantly higher MT renucleation activity than the surrounding cytoplasm (*Figure 6D*).

**Figure 6.**
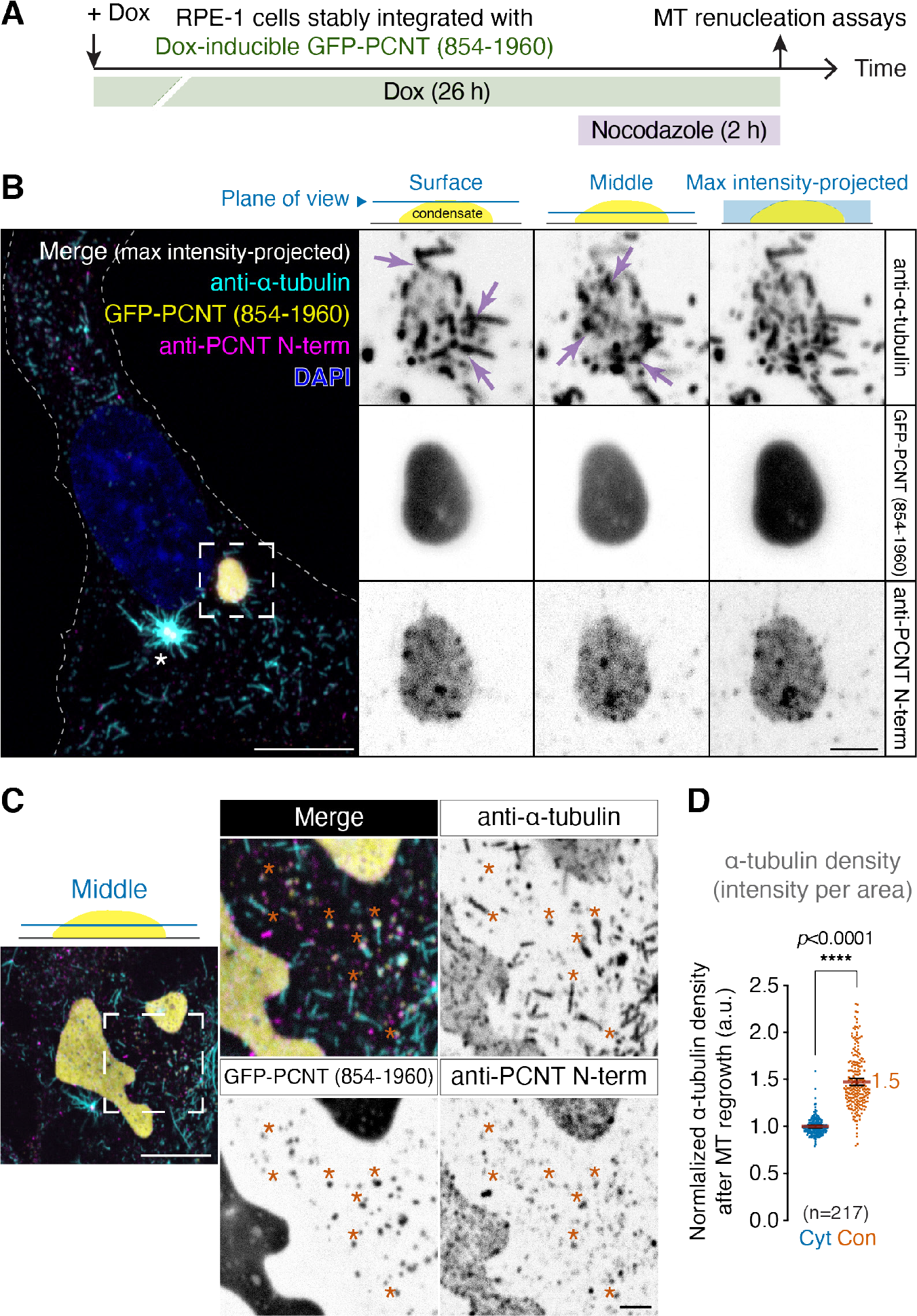
GFP-PCNT (854-1960) condensates nucleate MTs in cells. (**A**) Schematic of the MT renucleation assay to determine whether GFP-PCNT (854-1960) condensates nucleate MTs. (**B**) Anti-*a*-tubulin immunofluorescence of the cells containing GFP-PCNT (854-1960) condensates after MT renucleation in maximum intensity-projected and single optical section views. Note that MTs were renucleated within and on the surface of the condensate (arrows). The asterisk in the merged channels denotes the centrosome in which MT renucleation was robust. Similar MT renucleation in GFP-PCNT (854-1960) condensates was also observed in live cells (see ***Figure 6–Figure Supplement 3*** and ***Figure 6–video 1***). (**C**) MT renucleation also occurred in small PCNT condensates (asterisks), which also recruited endogenous PCNT. (**D**) Quantification of *a*-tubulin density (intensity per area) in GFP-PCNT (854-1960) condensates (Con) and in the surrounding cytoplasm (Cyt) after MT renucleation. Data are mean ± 95% CI. The mean of fold enrichment is noted. n, number of condensates analyzed from three biological replicates. The p-value was determined by the Student’s t-test (two-tailed). a.u., arbitrary unit. Scale bars, 10 µm and 2 µm (inset). See ***Figure 6–Figure Supplement 1*** for the quantification of *a*-tubulin density in GFP-PCNT (854-1960) condensates in the absence endogenous PCNT. **Figure 6–Figure supplement 1.** PCNT (854-1960) condensates can still nucleate MTs in the absence of endogenous PCNT. **Figure 6–Figure supplement 2.** PCNT (854-1960) condensates, but not HOTag condensates, nucleate MTs. **Figure 6–Figure supplement 3.** GFP-PCNT (854-1960) condensates nucleate MTs in live cells. **Figure 6–video 1.** A time-lapse movie of a GFP-PCNT (854-1960) condensate nucleating MTs.

We further found that the MT renucleation activity of PCNT (854-1960) condensate did not require endogenous PCNT (***Figure 6–Figure Supplement 1***), consistent with the results that in the absence of endogenous PCNT, many PCM components, including *γ*-tubulin, were still recruited to PCNT (854-1960) condensates (***Figure 5–Figure Supplement 3***, ***Group II and III***). As a negative control, we performed the MT renucleation assay in the cells with either GFP-PCNT (854-1960) or GFP-HOTag condensates (the condensates were induced as in the experiment shown in ***Figure 5-Figure supplement 1***). We found that only GFP-PCNT (854-1960) condensates, but not GFP-HOTag condensates, nucleated MTs (***Figure 6–Figure Supplement 2***). Thus, the GFP moiety in the fusion protein did not artifactually contribute to the MT nucleation activity of GFP-PCNT (854-1960) condensates. These results were also consistent with the observation that only PCNT condensates, but not HOTag condensates, recruited PCM components (***Figure 5–Figure Supplement 1***).

A similar MT renucleation assay was also performed in live cells, in which EB3-tdTomato was used to track the growing MT plus ends. Live cell imaging showed that some PCNT (854-1960) condensates nucleated MTs as EB3-tdTomato signals were emanating from the surface of PCNT condensates (***Figure 6–Figure Supplement 3*** and ***Figure 6***–***video 1***). Together, from these MT renucleation assays, we conclude that GFP-PCNT (854-1960) condensates possess the centrosome-like MT nucleation activity in cells, although this activity is significantly lower than that of the centrosome (e.g., compare the MT nucleation activities between the centrosome and the condensates shown in ***Figure 6B and Figure 6-Figure supplement 2***).

## Discussion

Our work shows that endogenously expressed PCNT, a core PCM protein important for centrosome maturation, forms dynamic pericentrosomal granules before incorporating into mitotic centrosomes in human cells. These PCNT granules are likely formed through LLPS because (1) they are sensitive to various aliphatic alcohols that are known to disrupt phase-separated cellular assemblies (***Kroschwald et al., 2017***; ***Larson et al., 2017***; ***Rog et al., 2017***; ***So et al., 2019***; ***Strom et al., 2017***); (2) the CC/LCR-rich portion of PCNT undergoes concentration-dependent condensation *in cellulo* that obeys characteristics of LLPS—including a defined phase transition boundary, condensate coalescence, deformability, fast recovery in FRAP experiments, and sensitivity to the 1,6-hexanediol treatment, the same treatment that also disrupts pericentrosomal PCNT granules. Recent theoretical modeling (***Zwicker et al., 2014***) and *in vitro* reconstitution studies (***Woodruff et al., 2017***) suggest that LLPS underlies centrosome assembly in *C. elegans*. To our knowledge, our study provides the first *in cellulo* evidence to support such a model and suggests that LLPS may also underlie the assembly of vertebrate centrosomes with at least one protein, PCNT, directly involved in this process.

### Is co-translational targeting of PCNT linked to its condensation during late G2/early mitosis?

The dynamic PCNT granules are predominantly observed during late G2/early mitosis when cotranslational targeting of PCNT to the centrosome peaks (***Sepulveda et al., 2018***). This raises an intriguing question of whether co-translational targeting facilitates the condensation of PCNT during this period. In the co-translational targeting model, multiple nascent PCNT polypeptides emerge from each polysome complex with a single large *PCNT* mRNA and are transported along the MT tracks. In principle, this process could effectively bring multiple N-terminal, LLPS-driving PCNT polypeptides in close proximity. A proximity-driven LLPS can thus be envisioned as LLPS is a concentration-dependent process. This could also explain why these dynamic PCNT granules are observed predominantly during late G2/early mitosis when PCNT production peaks. This proximity-driven phase separation model is also consistent with the results that the process of LLPS of the N-terminal (2-1960) and middle (854-1960) segments of PCNT can be reconstituted in a concentration-dependent manner in the cytoplasm, regardless of cell cycle stages. This model is also consistent with the observation that centrosomal targeting of PCNT (854-1960) condensates is a dynein- and MT-dependent process, and PCNT (854-1960) would phase separate closer to the centrosome with unperturbed dynein activities and intact MTs (***Figure 4D***). An important future goal will be to determine whether co-translational transport of PCNT and its condensation are indeed mechanistically linked, with the former facilitating the latter.

Many phase-separated cellular assemblies contain RNA and protein (*Elbaum-Garfinkle et al., 2015*; *Langdon et al., 2018*; *Lee et al., 2020*; *Lin et al., 2015*; *Maharana et al., 2018*; *Molliex et al., 2015*; *Schwartz et al., 2013*; *Zhang et al., 2015*), and RNA can promote or inhibit phase separation (*Fernandes and Buchan, 2020*; *Jain and Vale, 2017*; *Ma et al., 2021*; *Maharana et al., 2018*). Therefore, another important future goal is to determine whether RNA—e.g., *PCNT* mRNA, ribosomal RNA in the *PCNT* polysome, or other RNA—plays an active role in the formation or regulation of pericentrosomal PCNT granules under physiological conditions.

### Functional significance of PCNT condensation

What is the physiological significance of PCNT condensation? Would it facilitate the centrosomal targeting and incorporation of PCNT? Or more broadly, is the formation of dynamic PCNT granules a prerequisite for proper centrosome assembly? Given that PCNT (854-1960) condensates move toward the centrosome in a dynein- and MT-dependent manner (***Figure 4***), as in the case of cotranslational targeting of *PCNT* polysomes to the centrosome (***Sepulveda et al., 2018***), it is tempting to speculate that by combining co-translational targeting and protein condensation with motor-mediated active transport, the cell is thus able to target PCNT (and likely other PCM proteins) to mitotic centrosomes in an efficient and “protected” manner. For example, condensation could fight the force of diffusion, and this orchestrated transport along MT tracks could limit undesired interactions in the crowded cytoplasm before PCNT reaches the centrosome.

Woodruff and colleagues show that in the presence of crowding reagents, purified centrosomal protein SPD-5 forms liquid-like spherical condensates *in vitro*, which then rapidly “mature” into a gel- or solid-like state (***Woodruff et al., 2017***). Strikingly, only these spherical SPD-5 condensates can nucleate MTs, but not the solid-like SPD-5 scaffolds formed in the absence of crowding reagents (***Woodruff et al., 2017***, ***2015***). Their data thus suggest that formation of a condensate with liquid-like properties—even only transiently—might be important to allow centrosomal proteins to be properly assembled to possess the MT nucleation activity (***Raff, 2019***; ***Woodruff et al., 2017***). A similar scenario could happen here—the formation of liquid-like pericentrosomal PCNT granules may enable the proper assembly of the human centrosome to organize MTs, for example, by allowing various PCM components to “morph” into the desired configuration before becoming a gel/solid-like state (***Figure 1D, D’***). This “transitioning step” might be particularly important in forming large, micron-sized, membraneless assemblies such as the mitotic PCM.

Unfortunately, the ability to directly assess the biophysical properties of these small pericentrosomal PCNT granules is limited. However, new insights into the process of PCNT/PCM assembly have been obtained from studying the PCNT (854-1960) segment and the condensate it forms *in cellulo*. This segment is one of the most conserved regions of PCNT (***Figure 2***) and contains the sequence elements that drive LLPS (***Figure 3***). PCNT (854-1960) condensates show a molecular aging process (***Figure 3***) that resembles the possible liquid-to-gel/solid-like transition of the *in situ*-tagged GFP-PCNT granules. PCNT (854-1960) condensates also move toward the centrosome in a dynein- and MT-dependent manner (***Figure 4***), similar to how the *PCNT* polysomes are co-translationally transported to the mitotic centrosome (***Sepulveda et al., 2018***). Morphologically, the internal organization of PCNT (854-1960) condensates—with an inhomogeneous, porous appearance (***Figure 5***)—also resembles that of salt-stripped mitotic centrosomes purified from flies and clams in the electron tomography studies (***Moritz et al., 1995a***; ***Schnackenberg et al., 1998***), in which the PCM is shown as a fibrous, solid-like scaffold surrounding the centrioles. Moritz and colleagues further demonstrate that upon adding bovine tubulins, MTs regrow from the PCM of the salt-stripped centrosomes, with MT nucleation sites distributed throughout the PCM and MTs oriented in different directions (***Moritz et al., 1995a***). Interestingly, in our MT renucleation assays, we also observed a similar MT renucleation pattern in the PCNT (854-1960) condensate—MTs were nucleated throughout the condensate and regrown into different directions (***Figure 6***, ***Figure 6–Figure Supplement 3***, and ***Figure 6***–***video 1***). PCNT (854-1960) condensates and the isolated, reconstructed PCM scaffolds thus share a similar gross morphology and possess a centrosome-like MT nucleation activity. Taken together, results from our studies of the *in situ*-tagged full-length PCNT and phase-separating PCNT (854-1960) segment suggest that proper centrosome function may pivot on LLPS and liquid-to-gel/solid-like phase transition during the process of centrosome assembly.

An important future goal is to rationally design phase separation-deficient (and -rescuing) PCNT variants to determine the functional significance of phase separation *per se* in centrosome function. It is also important to determine whether other PCM components are co-condensed with pericentrosomal PCNT granules and/or undergo a similar condensation process during centrosome assembly.

### How is PCNT (854-1960) capable of recruiting endogenous PCM components and nucleating MTs?

It is surprising that the PCNT condensate formed by only one-third of PCNT (i.e., residues 854-1960) can selectively recruit endogenous centrosomal proteins and nucleate MTs *in cellulo*. This region is particularly enriched with coiled-coils (CCs) and low-complexity regions (LCRs) (***Figure 2A***). However, this region does not contain the putative *γ*-tubulin-binding domains, which are within the first 350 residues of human PCNT (***Lin et al., 2014b***) (***Figure 2A***), nor the CEP215-binding site, which is mapped to the C-terminus of human PCNT (residues 2390-2406) (***Kim and Rhee, 2014***). Yet both *γ*-tubulin and CEP215 (and several other PCM proteins) are recruited to the PCNT (854-1960) condensate (***Figure 5***). One explanation is that their recruitments are mediated through the endogenous PCNT, which is also recruited to the condensate. Indeed, in the absence of endogenous PCNT, CEP215 is no longer recruited to the PCNT (854-1960) condensate (***Figure 5–Figure Supplement 3***). However, without endogenous PCNT, most of other PCM proteins or clients we examined can still be recruited to the PCNT (854-1960) condensate—some proteins are recruited to a lesser extent (e.g., *γ*-tubulin and CEP192), while others are still recruited to the similar level as in the cells with endogenous PCNT (e.g., dynein and PLK1) (***Figure 5–Figure Supplement 3***). These results suggest that PCNT (854-1960) condensates recruit PCM proteins either indirectly (e.g., via the endogenous PCNT they also recruit) or directly (e.g., through yet to be identified binding sites for certain PCM proteins or clients). It is also possible that after PCNT (854-1960) phase separates, the resulting condensates gain new biophysical properties (e.g., a new binding environment) that are not present in PCNT (854-1960) monomers, thus enabling them to specifically recruit certain PCM proteins or clients. Combining mutagenesis and *in vitro* reconstitution experiments will help dissect the mechanisms underlying the selective recruitment of different PCM proteins and clients to PCNT (854-1960) condensates.

### Re-evaluating the role of coiled-coils in centrosome assembly

Coiled-coils are often enriched with low-complexity sequences (***Romero et al., 1999***). They are frequently predicted to be disordered as monomers but become folded upon formation of quaternary structures (coupled folding and binding) (***Anurag et al., 2012***; ***Szappanos et al., 2010***; ***Uversky et al., 2000***). Due to these unique properties, coiled-coils are known to adapt vast structural variations with different superhelical stabilities to exert a wide range of biological functions (***Grigoryan and Keating, 2008***; ***Li et al., 2003***; ***Rose and Meier, 2004***).

It has long been recognized that coiled-coil proteins are enriched at the centrosome and function as parts of the “centromatrix” for the recruitment of other proteins (***Doxsey, 2001***; ***Salisbury, 2003***; ***Schnackenberg and Palazzo, 1999***). Recent *in vitro* reconstitution studies of the coiled-coil PCM proteins SPD-5 (*C. elegans*) and Cnn (*D. melanogaster*) provide strong evidence to support a polymer-based mechanism of PCM assembly (***Feng et al., 2017***; ***Woodruff et al., 2017***, ***2015***). However, the exact mechanism underlying this polymer-based assembly is still under debate. It also remains unclear whether this model is applicable to vertebrate systems (***Gupta and Pelletier, 2017***; ***Raff, 2019***).

We found that in cultured human cells, not only endogenously expressed full-length PCNT condenses into dynamic granules, the coiled-coil-rich PCNT segments alone can undergo typical LLPS to form bioactive condensates with centrosome-like activities. These findings illuminate the fundamental principle of centrosome assembly and join a growing list of studies in which coiled-coilmediated phase separation participates in a variety of biological functions (***Fang et al., 2019***; ***Lu et al., 2020***; ***Rog et al., 2017***; ***Vega et al., 2019***; ***Zeng et al., 2016***).

Notably, coiled-coils (CCs) and low-complexity regions (LCRs) of human PCNT are enriched in the regions that are evolutionarily conserved, suggesting that these sequence features are under natural selection to preserve critical functions. We propose that PCNT is a linear multivalent protein that can undergo LLPS through its conserved CCs and LCRs to become spatially organized condensates that scaffold PCM assembly. This process is likely initiated during its co-translational targeting to the centrosome when the nascent PCNT polypeptides are in close proximity in the polysome (***Figure 7***). We propose that PCNT phase separation can achieve two main goals. First, it concentrates PCM proteins and clients as the PCNT condensates selectively recruit them. This will facilitate their incorporation into the centrosome and limit the biochemical reactions at the centrosome (e.g., MT nucleation, kinase activities) from taking place elsewhere in the cytoplasm. Second, it enables a liquid-to-gel/solid-like transitioning process during centrosome assembly. This process provides the PCM proteins a thermodynamically favorable pathway to assemble into a micron-sized, membraneless, and yet spatially organized PCM.

**Figure 7.**
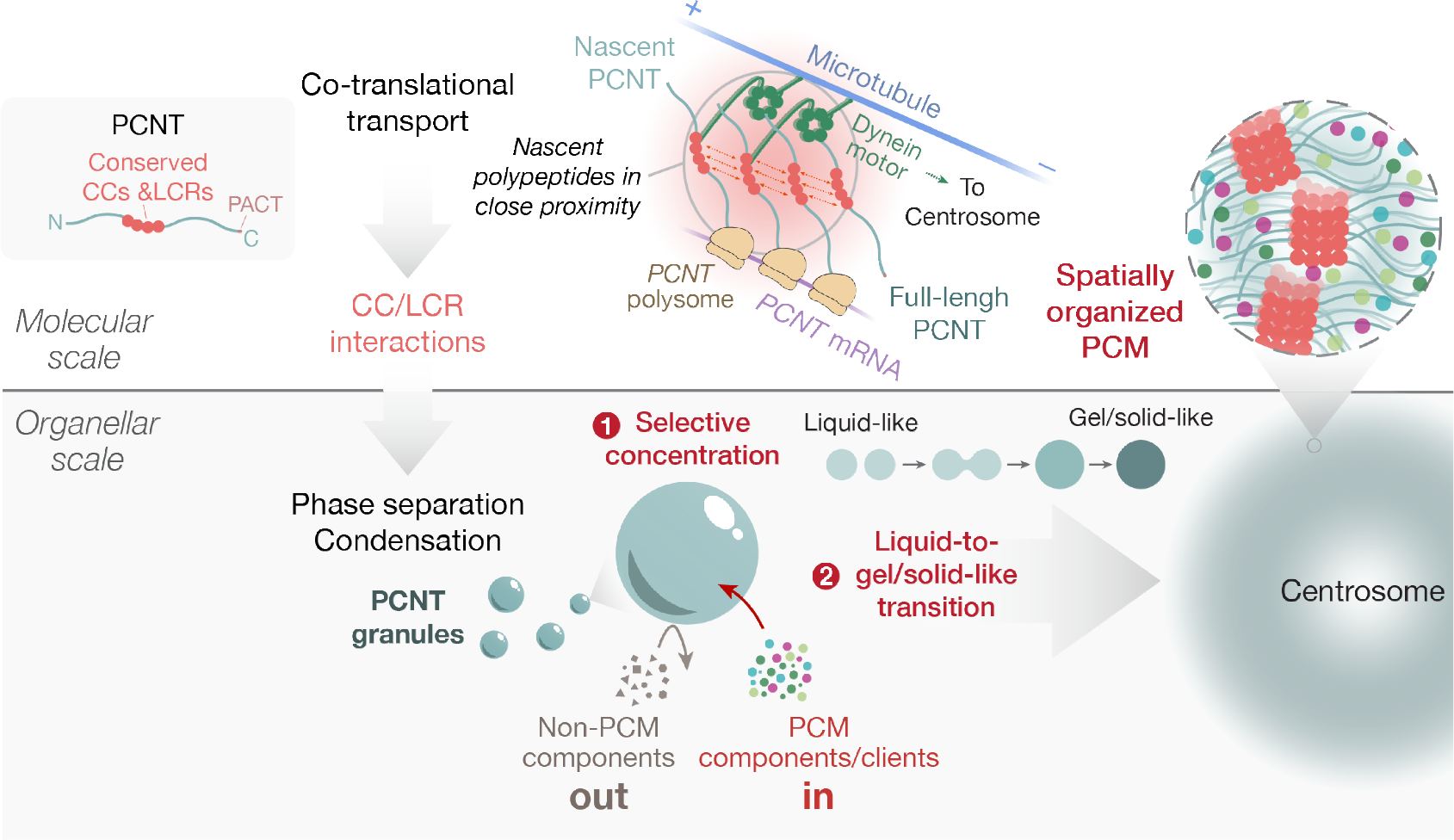
Model for PCNT phase separation in centrosome assembly. PCNT is a linear multivalent protein that phase separates through its conserved coiled-coils (CCs) and low-complexity sequence regions (LCRs) during its co-translational targeting to the centrosome when the nascent *PCNT* polypeptides are in close proximity in the polysome. The resulting PCNT granules/condensates promote PCM assembly by (1) selectively concentrating PCM components and clients; this will facilitate PCM assembly and limit thebiochemical reactions at the centrosome (e.g., MT nucleation, kinase activities) from occurring elsewhere in the cytoplasm and (2) enabling a liquid-to-gel/solid-like transitioning process during centrosome assembly; this process provides the PCM components a thermodynamically favored pathway to assemble into a micron-sized, membraneless, and yet spatially organized PCM.

Although CCs mediate other phase separation-independent activities, LLPS mediated by CC- and LCR-rich sequences observed in our study might be widespread among other CC-rich centrosomal proteins as previously suggested (***Woodruff et al., 2017***). Our results encourage future studies to rethink the conceptional framework regarding CC proteins and LLPS in centrosome assembly and to determine whether a unified mechanism is applied across metazoans.

## Methods and Materials

### Key resources table

| Reagent type | Reagent | Source | Cat. no. | Additional Information |
| --- | --- | --- | --- | --- |
| Chemical compound, drug | 1,7-heptanediol | Tokyo Chemical Industry Co., LTD., Tokyo, Japan | H0028 |  |
| Chemical compound, drug | 1,2-hexanediol | Tokyo Chemical Industry Co., LTD., Tokyo, Japan | H0688 |  |
| Chemical compound, drug | 1,6-hexanediol | MilliporeSigma, Burlington, MA | 240117 |  |
| Chemical compound, drug | 1,5-pentanediol | Spectrum Chemical Manufacturing Corporation, New Brunswick, NJ | P2088 |  |
| Chemical compound, drug | Ciliobrevin D | MilliporeSigma, Burlington, MA | 250401 |  |
| Chemical compound, drug | Doxycycline hyclate | MilliporeSigma, Burlington, MA | D9891 |  |
| Chemical compound, drug | Dynarrestin | MilliporeSigma, Burlington, MA | SML2332 |  |
| Chemical compound, drug | Nocodazole | MilliporeSigma, Burlington, MA | M1404 |  |
| Chemical compound, drug | Rapamycin | LC Laboratories, Woburn, MA | R-5000 |  |
| Chemical compound, drug | RO-3306 | R & D Systems, Minneapolis, MN | 4181 |  |

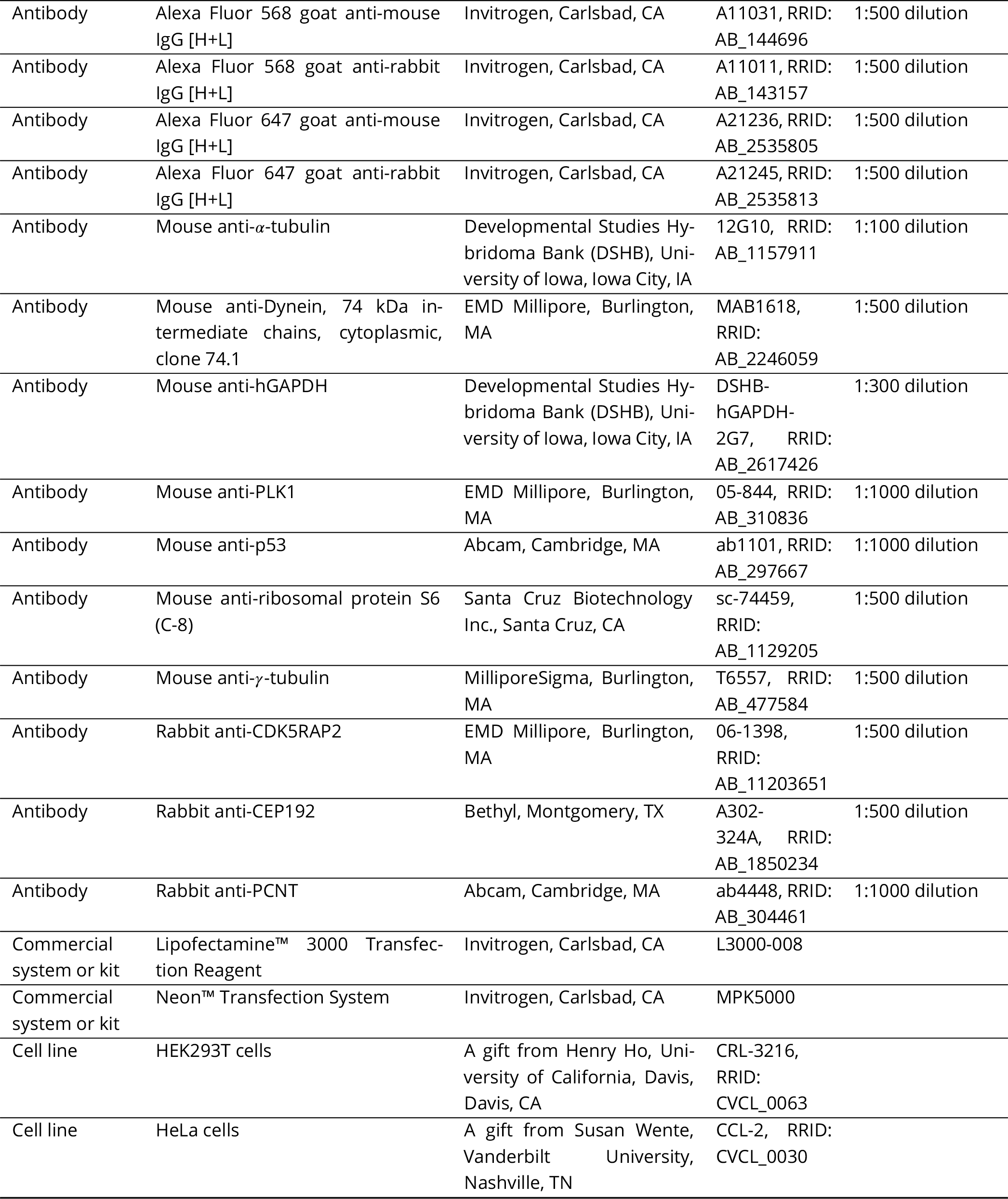

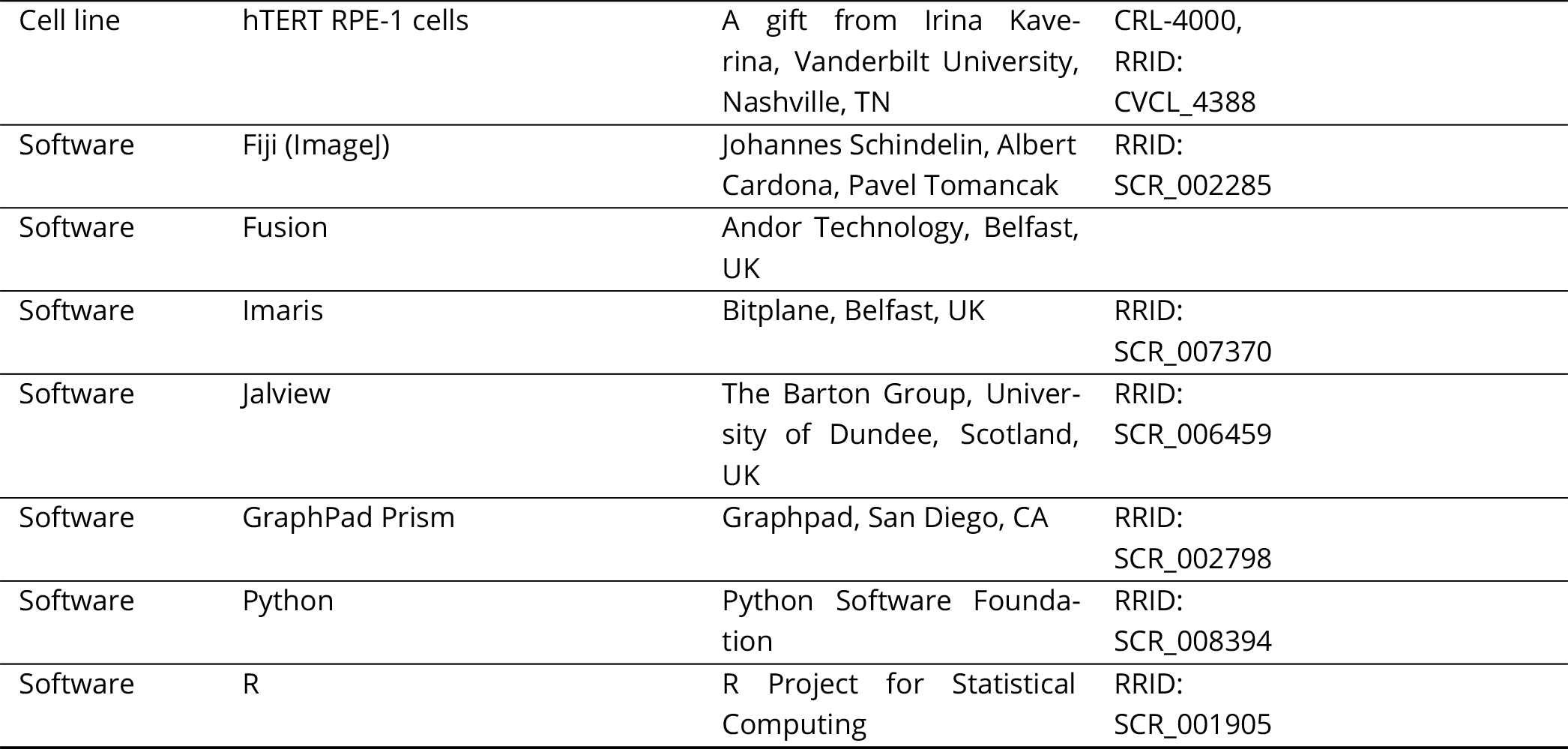

### Constructs

To generate constructs for stable expression of GFP- or mScarlet-i-PCNT segments controlled by a doxycycline (Dox)-inducible promoter, each PCNT segment was first amplified by PCR from pCMV-3xFLAG-EGFP-PCNT-Myc plasmid (a kind gift from Kunsoo Rhee, Seoul National University, Seoul, South Korea) (***Lee and Rhee, 2011***) and assembled into a vector with sfGFP or mScarlet-i by Gibson assembly (***Gibson et al., 2009***). The final constructs with the *piggyBac* transposon elements and Dox-inducible promoter were made by subcloning the sfGFP- or mScarlet-i-PCNT fragment into PB-TA-ERN (a gift from Knut Woltjen, Addgene plasmid #80474) (***Kim et al., 2016***) using the Gateway cloning system (Thermo Fisher Scientific, Waltham, MA). The following *piggyBac* transposon constructs, with amino acid sequences of human PCNT in the parentheses, were used in this study: PB-TA-sfGFP-PCNT (2-891), PB-TA-sfGFP-PCNT (854-1960), PB-TA-sfGFP-PCNT (2-1960), PB-TA-sfGFP-PCNT (1954-3336), and PB-TA-mScarlet-i-PCNT (854-1960).

To make the construct for labeling microtubule (MT) plus ends, the EB3-tdTomato fragment was amplified by PCR from EB3-tdTomato (a gift from Erik Dent, Addgene plasmid #50708) (***Merriam et al., 2013***) and cloned into a lentiviral targeting plasmid pLVX-EF1*α*-mCherry-N1 without the mCherry portion (631986, Takara Bio, Mountain View, CA) by Gibson assembly. The resulting plasmid, pLVX-EF1*α*-EB3-tdTomato, was used to make lentiviruses to transduce cultured cells.

To make the construct for labeling DNA, mScarlet-i-H2A construct was amplified by PCR from pmScarlet-i_H2A_C1 (a gift from Dorus Gadella, Addgene plasmid #85053) (***Bindels et al., 2017***) and cloned into the same lentiviral targeting construct by Gibson assembly as described above to generate pLVX-EF1*α*-mScarlet-i-H2A. The mScarlet-i-H2A construct was also cloned into a *piggybac* vector with the cDNA under the control of EF1*α* promoter (PB-EF1*α*-GW, a gift from Henry Ho) by the Gateway cloning system to generate PB-EF1*α*-mScarlet-i-H2A.

To make the construct for labeling the centrioles with far red fluorescence, the coding sequence of human centrin 2 (CETN2) was first cloned from total RNA of HEK293T cells using the SuperScript® III One-Step RT-PCR System (Invitrogen). The miRFP670 was amplified by PCR from pmiRFP670-N1 (a gift from Vladislav Verkhusha, Addgene plasmid #79987) (***Shcherbakova et al., 2016***). miRFP670 and CETN2 were then assembled into the same lentiviral targeting construct by Gibson assembly as described above to generate pLVX-EF1*α*-miRFP670-CETN2. The miRFP670-CETN2 construct was also cloned into PB-EF1*α*-GW described above by the Gateway cloning system to generate PB-EF1*a*-miRFP670-CETN2.

To generate the Cas9 tagged with the nuclear localization signal (NLS) at both the N- and C-termini for expression in mammalian cells, an NLS from SV40 large T-antigen (***Kalderon et al., 1984***) was synthesized and added to hCas9, a construct encoding a human codon-optimized Cas9 (hCas9) with an NLS at its C-terminus (a gift from George Church, Addgene plasmid #41815) (***Mali et al., 2013***), by PCR. The final construct, pCMVSP6-NLS-hCas9-NLS-polyA_Tol2pA2, with NLS-hCas9-NLS under the control of the cytomegalovirus (CMV) immediate-early enhancer and promoter was generated by the Gateway cloning system using the components from the Tol2kit (***Kwan et al., 2007***).

### Cell culture

RPE-1 cells (a gift from Irina Kaverina, Vanderbilt University) were maintained in Dulbecco’s Modification of Eagles Medium/Ham’s F-12 50/50 Mix (DMEM/F-12) (10-092-CV, Corning, Corning, NY). HeLa cells (ATCC CCL-2, a gift from Susan Wente, Vanderbilt University, Nashville, TN) and HEK293T cells (a gift from Henry Ho, University of California, Davis, Davis, CA) were maintained in DMEM (10-017-CV, Corning). All cell lines were supplemented with 10% fetal bovine serum (FBS) (12303C, lot no. 13G114, Sigma-Aldrich, St. Louis, MO), 1x Penicillin Streptomycin (30–002 CI, Corning), and maintained in a humidified incubator with 5% CO_2_ at 37°C.

Cell lines used in this study were not further authenticated after obtaining from the sources. All cell lines were tested negative for mycoplasma using a PCR-based testing with the Universal Mycoplasma Detection Kit (30-1012K, ATCC, Manassas, VA). None of the cell lines used in this study was included in the list of commonly misidentified cell lines maintained by International Cell Line Authentication Committee.

### Generation of GFP-PCNT knock-in cell lines

The CRISPR/Cas9 technology with a double-cut homology directed repair (HDR) donor was used to knock in sfGFP into the *PCNT* locus of RPE-1 cells (***Lin et al., 2014a***; ***Zhang et al., 2017***). Because it was unclear if knocking in sfGFP would perturb centrosome integrity that might lead to p53-dependent cell cycle arrest (***Mikule et al., 2007***), making it unfavorable to isolate the knock-in clones, we first generated a *TP53* knockout RPE-1 cell line before the knock-in experiment (***Figure 1–Figure Supplement 5***). Knocking out *TP53* was achieved through the CRISPR-mediated gene editing by co-expression of Cas9 protein with the gRNA targeting *TP53* (5’-GGGCAGCTACGGTTTCCGTC-3’) using the method described by the Church group (***Mali et al., 2013***). The gRNA template was cloned into the gRNA Cloning Vector (Addgene plasmid # 41824) via Gibson assembly. 1 µg Cas9 plasmid (pCMVSP6-NLS-hCas9-NLS-polyA_Tol2pA2) and 1 µg gRNA plasmid were transfected into RPE-1 cells using the Lipofectamine 3000 reagent according to the manufacturer’s instruction (Invitrogen, Carlsbad, CA). Cells were expanded, isolated as single colonies, and screened for frameshift mutations in both *TP53* alleles by high-throughput Illumina sequencing. The loss of TP53 expression was further confirmed by Western blot analysis. A *TP53^-/-^* RPE-1 cell line (RPE-1_1-1) was used in this study (***Figure 1–Figure Supplement 5***).

To knock in sfGFP sequence into the *PCNT* locus, crRNA/tracrRNA (i.e., the Alt-R system, Integrated DNA Technologies, Coralville, IA) were used to target a sequence near the start codon of *PCNT* (CGCGCGGAGTCTGAGGGAGA). The double-cut HDR donor contains the sfGFP-PCNT cassette with 600-bp homology arms flanked by the same guide RNA target sequence (synthesized and cloned into pUC57-Kan vector, Genewiz, South Plainfield, NJ) (***Figure 1–Figure Supplement 1***). Annealed crRNA/tracrRNA and Cas9 protein (a kind gift from Fuguo Jiang and Jennifer Doudna, (***Jiang et al., 2016***, ***2015***) were incubated in 30 mM HEPES, pH 7.5, 1 mM MgCl_2_, 200 mM KCl, and 1 mM TCEP at 37°C for 10 min to form the Cas9 ribonucleoprotein (RNP) complex. Before electroporation, the HDR donor plasmid was mixed with 2 × 10^5^ of *TP53^-/-^* RPE-1 cells synchronized to early M phase using RO-3306 as done before (***Sepulveda et al., 2018***). The Cas9 RNP complexes were then mixed with the cell/donor vector mix, followed by electroporation using the Neon electroporation system with a 10-µl tip according to the manufacturer’s instruction (Pulse voltage: 1200 V, pulse width: 25, pulse number: 4, Invitrogen). The final concentrations of the annealed crRNA/tracrRNA, Cas9 protein, and HDR donor plasmid are 3 µM, 2 µM, and 120 nM, respectively, in a total volume of 10 µl. After electroporation, the cells were grown for 10-14 days at low density. Individual clones were isolated and screened for the presence of GFP-positive centrosomes. The GFP-positive clones were further confirmed by anti-PCNT immunostaining and sequencing the junctions of the sfGFP integration site (primer sequences are in ***Supplementary ile 1***).

### Generation of *PCNT* knockout cell lines

Disrupting *PCNT* was achieved by electroporating the Cas9 RNP complex into *TP53^-/-^* RPE-1 cells as done for generating the GFP-PCNT knock-in cells described above, except that no donor plasmid was included the RNP complex. The gRNA target sequence is near the start codon of *PCNT* (AGAGCAGCGGCGCAGAAAGG). After electroporation, the cells were grown for 10-14 days at low density. Individual clones were isolated and screened for the loss of PCNT by anti-PCNT immunostaining. The *PCNT* knockout clones were further confirmed by anti-PCNT Western blotting (***Figure 5–Figure Supplement 2***).

### Generation of stable cell lines by *piggybac* transposon-mediated integration

We used a previously described *piggyBac* transposon system (***Kim et al., 2016***) to generate stable cell lines that express proteins under the control of a Dox-inducible promoter (e.g., various GFP- and mScarlet-i-PCNT fusion proteins) or of the EF1*α* promoter (e.g., miRFP670-CETN2 and mScarlet-i-H2A fusion proteins). In brief, the *piggyBac* transposon plasmid that contains the desired transgene and a *piggyBac* transposase plasmid (PB210PA-1, System Biosciences, Palo Alto, CA) were electroporated into *TP53^-/-^* RPE-1 cells simultaneously using the Neon electroporation system according to the manufacturer’s instruction (Invitrogen). After 8-10 day of 200 mg/ml G418 treatment, transgene-integrated cells were screened by fluorescence signals. Sometimes fluorescence activated cell sorting (FACS) was further performed to obtain cells with desired, more uniform expression of the fusion proteins.

### Generation of stable cell lines by lentiviral transduction

To generate recombinant lentiviruses expressing EB3-tdTomato, pLVX-EF1*α*-EB3-tdTomato plasmid was co-transfected with the following third-generation packaging plasmids (gifts from Didier Trono): pMDLg/pRRE, pRSV-Rev, and pMD2.G (Addgene plasmids #12251, #12253, and #12259, respectively) (***Dull et al., 1998***) into HEK293T cells. Viral supernatants were collected from media 24-48 h post transfection, filtered by a 0.22-µm filter, and were used to infect the inducible GFP-PCNT (854-1960) *TP53^-/-^* RPE-1 cells with 8 µg/ml polybrene. 18-24 h post infection, the viruses were removed, and the cells were expanded. To minimize the impact on MT dynamics, the cells expressing low levels of EB3-tdTomato (gated and collected by FACS) were used for experiments. To generate lentiviruses expressing mScarlet-i-H2A and miRFP670-CETN2 fusion proteins, the same lentiviral packaging procedure was performed as above except for using the targeting vectors pLVX-EF1*α*-mScarlet-i-H2A and pLVX-EF1*α*-miRFP670-CETN2, respectively. The resulting viral supernatants were then used to infect the cells of interest. Sometimes FACS was further performed to obtain cells with desired, more uniform expression of the fusion proteins.

### Immunostaining

Immunostaining was performed as previously described (***Sepulveda et al., 2018***; ***Jiang et al., 2019***). In brief, cells were fixed for 15 min in 4% paraformaldehyde in 1xPBS at room temperature or 5 min in ice-cold 100% methanol at −20°C. Cells were then washed twice with 1x PBS and incubated with Blocking Solution (2% goat serum, 0.1% Triton X-100, and 10 mg/ml bovine serum albumin in 1x PBS) for 1 h at room temperature. Cells were then incubated with Blocking Solution containing diluted primary antibody for 1 h at room temperature, washed three times with 1x PBS, and incubated with Blocking Solution containing diluted secondary antibody for 1 h at room temperature. Cells were washed three times with 1x PBS, and nuclei were counterstained with 0.05 mg/ml 4’,6-diamidino-2-phenylindole (DAPI) in 1x PBS for 30 min at room temperature before mounting.

### Microscopy

Microscopy was performed using a spinning disk confocal microscope system (Dragonfly, Andor Technology, Belfast, UK) with 63x/1.40 (magnification/numerical aperture) or 100x/1.40 HC PL APO objectives (Leica, Wetzlar, Germany), coupled with 1x, 1.5x, or 2x motorized magnification changer. Image acquisition was controlled by Fusion software (Andor Technology) and images were captured with an iXon Ultra 888 EMCCD or Zyla sCMOS camera (Andor Technology). Sometimes deconvolution of the images was also performed using the Fusion software (Andor Technology).

All live cell imaging was performed with cells seeded in 35-mm glass-bottom dishes (P35G-1.5-10-C, MatTek Corp., Ashland, MA or D35C4-20-1.5-N, Cellvis, Mountain View, CA) mounted in a humidified chamber supplied with 5% CO_2_ inside a wrap-around environmental incubator (Okolab, Pozzuoli, Italy) with temperature set at 37°C.

For acute treatments of live cells with aliphatic alcohols, the aliphatic alcohol was prepared in 10% FBS DMEM/F-12 media, pre-warmed to 37°C, and added onto cells mounted on the microscope stage. Time lapse microscopy was performed before and immediately after the aliphatic alcohol addition. For the control, cells were imaged under the same acquisition conditions except that the cells were treated with 10% FBS DMEM/F-12 media alone.

### Quantification of PCNT granule numbers at different cell cycle stages

For counting PCNT granules, confocal images of GFP-PCNT knock-in cells stably expressing mScarlet-i-H2A and miRFP670-CETN2 were converted to 8-bit color space after maximum intensity projection using Fiji (***Schindelin et al., 2012***). An intensity threshold was then applied to separate the GFP-PCNT granules from background signals. The number of PCNT granules was then counted using the *Analyze Particles* function in Fiji.

The cell cycle stages of these cells were determined by analyzing the distance between centrosomes, the number of centrin dots, and the DNA morphology. A G1/early S-phase cell is defined as the cell that contains one centrosome with two centrin dots and its DNA is not condensed; a S/early G2-phase cell is defined as the cell that contains two centrosomes close together (less than 1 µm apart), each with a pair of centrin dots (i.e., usually one brighter than the other, representing the mother and daughter centrioles), and its DNA is not condensed; a late G2-phase cell is defined as the cell that contains two centrosomes greater than 1 µm apart, each with a pair of centrin dots, and its DNA is not condensed, but the PCM has started to expand; a prophase cell is defined as the cell that contains condensed DNA, but the nuclear envelope (NE) is still intact; a prometaphase cell is defined as the cell that contains condensed DNA and the NE has broken down; a metaphase cell is defined as the cell with condensed DNA aligned along the equator of the cell; an anaphase cell is defined as the cell with its condensed chromosomes just starting to segregate toward the two centrosome/spindle poles.

### Analysis of the movement of PCNT condensates upon dynein inhibition or MT de-polymerization

#### Drug treatments and microscopy

To assess the movement of PCNT condensates upon dynein inhibition, the *TP53^-/-^* RPE-1 cells with stably integrated GFP-PCNT (854-1960) constructs under the control of a Dox-inducible promoter (herein named Tet-ON-GFP-PCNT (854-1960) cells) were first incubated with 1 µg/ml Dox for 2-3 h, followed by incubation with ciliobrevin D and dynarrestin mix (50 µM each) and 1 µg/ml Dox for 1 h before the start of time-lapse microscopy. To assess the movement of PCNT condensates upon MT depolymerization, a mix of 1 µg/ml Dox and 8.3 µM nocodazole was added to Tet-ON-GFP-PCNT (854-1960) cells for 2-3 h before the start of time-lapse microscopy. For the control, the same experimental procedure was performed except that the cells were treated with the DMSO vehicle alone. Cells were imaged at 4-min intervals for 6-10 h in the presence of 5% CO_2_ at 37°C for all conditions.

#### Quantification

Confocal images after maximum intensity projection were split into individual channels for GFP-PCNT (854-1960) (condensate), mScarlet-i-H2A (nucleus), and miRFP670-CETN2 (centrosome). For each channel, intensity and size thresholds were applied to identify the objects of interest—i.e., condensates, nuclei, and centrosomes—as “masked” objects. *Analyze Particle* function of Fiji was then applied to assign each masked object a unique identification (ID) number across all time frames. The (X,Y) coordinates (based on the center of mass) and the area of each masked object were also calculated using Fiji.

Because ID numbers are unique, the same masked object will be represented by different ID numbers across time frames. To track the same object with different ID numbers over time, we considered that cells only moved slightly between time points with 4-min intervals, and that the nucleus was the least mobile among these three masked objects. Therefore, the ID numbers representing the same nucleus will also have the (X, Y) coordinates shifted the least between any two consecutive time points. Using this feature, we developed a Python script (***Figure 4***–***source code 1***) to automatically assign a set of ID numbers to a given nucleus across time frames so that the same nuclei could now be “tracked” frame by frame. In each time frame, this Python script also paired the condensate and centrosome objects with the nucleus of each cell in the field. Together, we were able to track all three masked objects simultaneously in each cell across time frames. After executing the script, we manually confirmed the accuracy of the pairing process and corrected any errors.

Once the tracking of all three objects across time frames was completed, the condensate size, condensate number, and its distance to the centrosome in each cell over time were calculated. Data computation was done using Pandas (***Reback et al., 2020***) and NumPy (***Harris et al., 2020***). Data visualization was performed with Matplotlib (***Hunter, 2007***) and GraphPad Prism (GraphPad, San Diego, CA). Although phase separation occurred asynchronously, data were aligned and plotted from the start of phase separation (time 0) for any given condensate (***Figure 4***).

### Fluorescence recovery after photobleaching (FRAP) experiments

FRAP experiments were performed using a FRAP photoablation module with a computer-controlled fiber optically pumped dye laser to bleach a region of interest (ROI) (∼2 µm in diameter) on the condensate or the centrosome after a few pre-bleach images were acquired. After photobleaching, the same ROI continued to be imaged at 2- to 5-s intervals for 5 to 12 min. Images were acquired using a Zeiss AxioObserver with a 60x objective coupled with a Yokogawa CSU-10 spinning disk confocal system and a Photometrics CoolSNAP HQ2 cooled CCD camera (BioImaging Solutions, San Diego, CA). The microscope system was enclosed in an environmental chamber with temperature set at 37°C. The photoablation and image acquisition were controlled by SlideBook software (Intelligent Imaging Innovations, Denver, CO). Images and data were analyzed using ImageJ, Microsoft Excel, and GraphPad Prism (GraphPad).

### Measurement of relative protein concentrations in cells

Dox-inducible cell lines expressing individual GFP-PCNT segments were seeded on glass-bottom 35-mm dishes and imaged after addition of 1 µg/ml Dox. Imaging of different cell lines was performed with the same acquisition setting and conditions (5% CO_2_ at 37°C) using a spinning disk confocal microscope system (Dragonfly, Andor Technology). To estimate the relative protein concentrations of GFP-PCNT in cells (outlined in ***Figure 2–Figure Supplement 3***), the volume of individual condensates and their surrounding cytoplasm were first determined from confocal voxels. This was done by performing surface rendering of the GFP signals of the condensates (dense phase) and of the whole cytoplasm (dense and light phases) with different thresholds using the *Surface* reconstruction function of Imaris (Bitplane, Belfast, UK). Depending on the GFP expression level in each cell, threshold values were manually adjusted for rendering. The volume of the rendered surface and and intensity sum of the GFP signals within the rendered surface were then calculated. The volume and intensity sum in the light phase were calculated by subtracting the value in the dense phase from that in the whole cytoplasm. Relative protein concentrations were calculated as intensity sum per volume. The critical concentration of phase-separated PCNT segments was determined as the concentration of the light phase when phase separation just occurred.

### Recruitment assays of proteins to the condensates

To test whether the PCNT condensates recruit PCM proteins, about 5 × 10^4^ of Tet-ON-GFP-PCNT (854-1960) cells were seeded onto each 12-mm circular coverslip (72230-01, Electron Microscopy Sciences, Hatfield, PA) in a 24-well plate and treated with 1 µg/ml Dox for 24 h to induce the formation of PCNT (854-1960) condensates. The cells were then fixed and immunostained against various PCM components or clients.

To compare the recruitment of PCM proteins between the PCNT and non-PCNT ondensates, the condensate formed by the HOTags was chosen as a non-PCNT condensate control (***Zhang et al., 2018***). The HOTag condensates were formed by transfecting pcDNA3-FKBP-EGFP-HOTag3 and pcDNA3-FRB-EGFP-HOTag6 plasmids (gifts from Xiaokun Shu; Addgene plasmid #106924 and #106919, respectively) into *TP53^-/-^* RPE-1 cells using the Lipofectamine 3000 reagent according to the manufacturer’s instruction (Invitrogen). 11 h post transfection, transfected cells were treated with 100 nM rapamycin for 1 h to induce the formation of HOTag condensates. In parallel, Tet-ON-GFP-PCNT (854-1960) cells were treated with 1 µg/ml Dox for 6 h to induce the formation of PCNT condensates—which are similar in size to the HOTag condensates at this stage—before immunostaining. In the last hour of Dox induction, 100 nM rapamycin was also added along with Dox to control for the potential effect caused by rapamycin.

To quantify the fold enrichment of various proteins in the condensates, three randomly chosen fixed areas in the condensate and in the cytoplasm per cell (2.25 µm^2^ for data in ***Figure 5***, 0.159 µm^2^ for data in ***Figure 5–Figure Supplement 1*** were selected for quantification using Fiji. The mean intensity values from the condensate or cytoplasm (from three areas each) in each cell were averaged. The fold enrichment of a given protein in the condensate is calculated as the ratio of protein intensity mean in the condensates to the overall protein intensity mean in the cytoplasm across all cells. For cells with centrosomes embedded in the PCNT condensates, PCM protein signals at the centrosomes within the PCNT condensate were excluded from quantification.

### MT renucleation assay

MT renucleation assay was adapted from previous studies (***Jao et al., 2017***; ***Sanders et al., 2017***). In brief, cells were treated with 8.3 µM nocodazole for 2 h to depolymerize MTs. The 24-well plate containing the treated cells was then placed on ice and the cells were washed with ice-cold media for 8 times to remove nocodazole. To allow MTs to renucleate, the plate was then placed in a 37°C water bath while the cells were incubated with pre-warmed media for 60-120 s (the optimal regrowth period varied between experiments), followed by a 10-s incubation with pre-warmed Extraction Buffer (60 mM PIPES, pH 6.9, 25 mM HEPES, 10 mM EGTA, 2 mM MgCl_2_, 0.1% Saponin, 0.25 nM nocodazole, 0.25 nM taxol). Immediately after the Extraction Buffer treatment, the cells were fixed with 4% paraformaldehyde and 0.025% glutaraldehyde in Cytoskeleton Buffer (10 mM MES, 150 mM NaCl, 5 mM MgCl_2_, 5 mM EGTA, 5 mM D-glucose) for 10 min at room temperature. The cells were then incubated with 0.2% NaBH4 in 1x PBS for 10 min at room temperature to quench the autofluorescence before immunostaining.

For the live MT renucleation assay, the nocodazole-treated EB3-tdTomato-expressing cells with PCNT condensates were mounted on the microscope stage, first imaged in the presence of nocodazole, followed by washes with pre-warmed media 5 times on the microscope stage. After washes, the cells were imaged immediately again at 1-min intervals to monitor MT renucleation.

### Quantification of ***a***-tubulin density in the MT regrowth assay

To quantify the *α*-tubulin density in the condensate, the condensate was first manually outlined as the first ROI using the freehand selection tool in Fiji. The second ROI was then defined as the cytoplasm space between the outline of the first ROI and the outline 1 µm larger than the first ROI. The area and intensity sum of anti-*α*-tubulin signals in the first (condensate) and second (cytoplasm) ROIs were then measured in Fiji. The *α*-tubulin density was calculated as anti-*α*-tubulin intensity sum per area. The normalized *α*-tubulin density was calculated as the ratio of *α*-tubulin density in the condensate to the averaged *α*-tubulin density in the cytoplasm. The presence of MT regrowth was also confirmed by confocal imaging.

### Multiple protein sequence alignments

Protein sequences of pericentrin orthologs from a diverse group of vertebrates, as well as two functional homologs of pericentrin from budding yeast (Spc110) and fruit fly (pericentrin-like protein, D-Plp), were retrieved from Ensembl genome database (http://uswest.ensembl.org), resulting in a total of 233 sequences. We further filtered sequences to eliminate those with low-quality sequence reads, incomplete annotations, and those inducing large gaps (e.g., due to the insertions specific to small numbers of species), resulting in a final list of 169 sequences. These 169 sequences were then aligned using MUSCLE (***Edgar, 2004***) and colored with the default Clustal X color scheme in Jalview (***Figure 2–Figure Supplement 1***) (***Waterhouse et al., 2009***). The conservation of each residue was scored using the AMAS method of multiple sequence alignment analysis in Jalview (***Livingstone and Barton, 1993***).

### Statistical analysis

Statistical analysis was performed using GraphPad Prism (GraphPad). Each sample size (n value) is indicated in the corresponding figure or figure legend. Significance was assessed by performing the Student’s t-test, the F-test, or one-way ANOVA, as indicated in individual figures. The experiments were not randomized. The investigators were not blinded to allocation during experiments and outcome assessment.

## Acknowledgments

We thank Dr. Bo Huang for discussion with the CRISPR knock-in strategy; Dr. Kunsoo Rhee for human PCNT cDNA; Dr. Rick McKenney for the sfGFP construct; Dr. Henry Ho for HEK293T cells, pLVX-EF1*α*-mCherry-N1, *piggybac* constructs, lentiviral packaging plasmids, and critical reading of the manuscript; Dr. Susan Wente for HeLa cells; Dr. Irina Kaverina for RPE-1 cells; Drs. Fuguo Jiang and Jennifer Doudna for Cas9 protein; Dr. Megan Dennis for analyzing Illumina sequencing results; Stephen (Evan) Brahms, Marvin Orellana, Hashim Shaikh, Janice Tam, and Alan Zhong for technical help with cell culture work; all members of the Jao and Ho labs for discussion; Emily Jao for help with illustration of the model figure. Experiments were performed in part through the use of Davis Campus Shared Flow Cytometry Resource. The article is dedicated to the memory of Dr. Fuguo Jiang, who recently passed away. Fuguo purified the Cas9 protein and kindly shared it with us for this study.

---

**Figure 1–Figure supplement 1.**
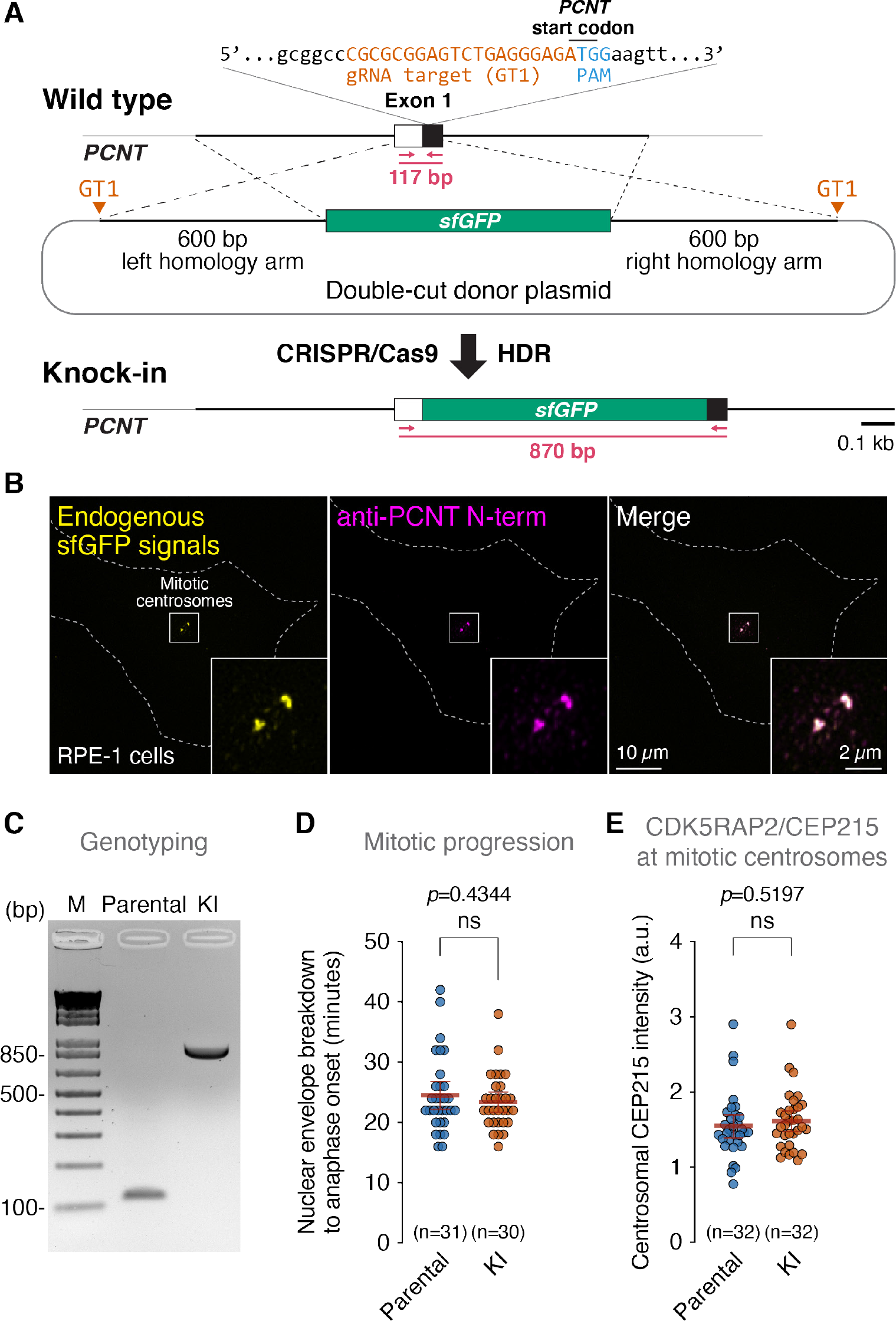
Generation and validation of GFP-PCNT knock-in cells. (**A**) Schematic of knocking in *sfGFP* into the *PCNT* locus of RPE-1 cells using the CRISPR/Cas9 technology with a double-cut homology-directed repair (HDR) donor. Arrows indicate the locations of genotyping primers. (**B**) Immunofluorescence images of the GFP-PCNT knock-in (KI) cells. Similar results were obtained from three biological replicates. (**C**) Genotyping results of one of the GFP-PCNT KI clones and the parental cells by PCR using the primers indicated in (A). Similar results were obtained from three biological replicates. (**D**) Quantification of mitotic progression of the parental and GFP-PCNT KI RPE-1 cells stably expressing miRFP670-CETN2 and mScarlet-i-H2A (which mark the centrosome and DNA, respectively). (**E**) Quantification of CDK5RAP2/CEP215 levels at mitotic centrosomes of the parental and GFP-PCNT KI RPE-1 cells. Data are mean ± 95% CI. The p-value was determined by the Student’s t-test (two-tailed). ns, not significant. n, number of cells analyzed from three (D) and two (E) biological replicates. a.u., arbitrary unit.

**Figure 1–Figure supplement 2.**
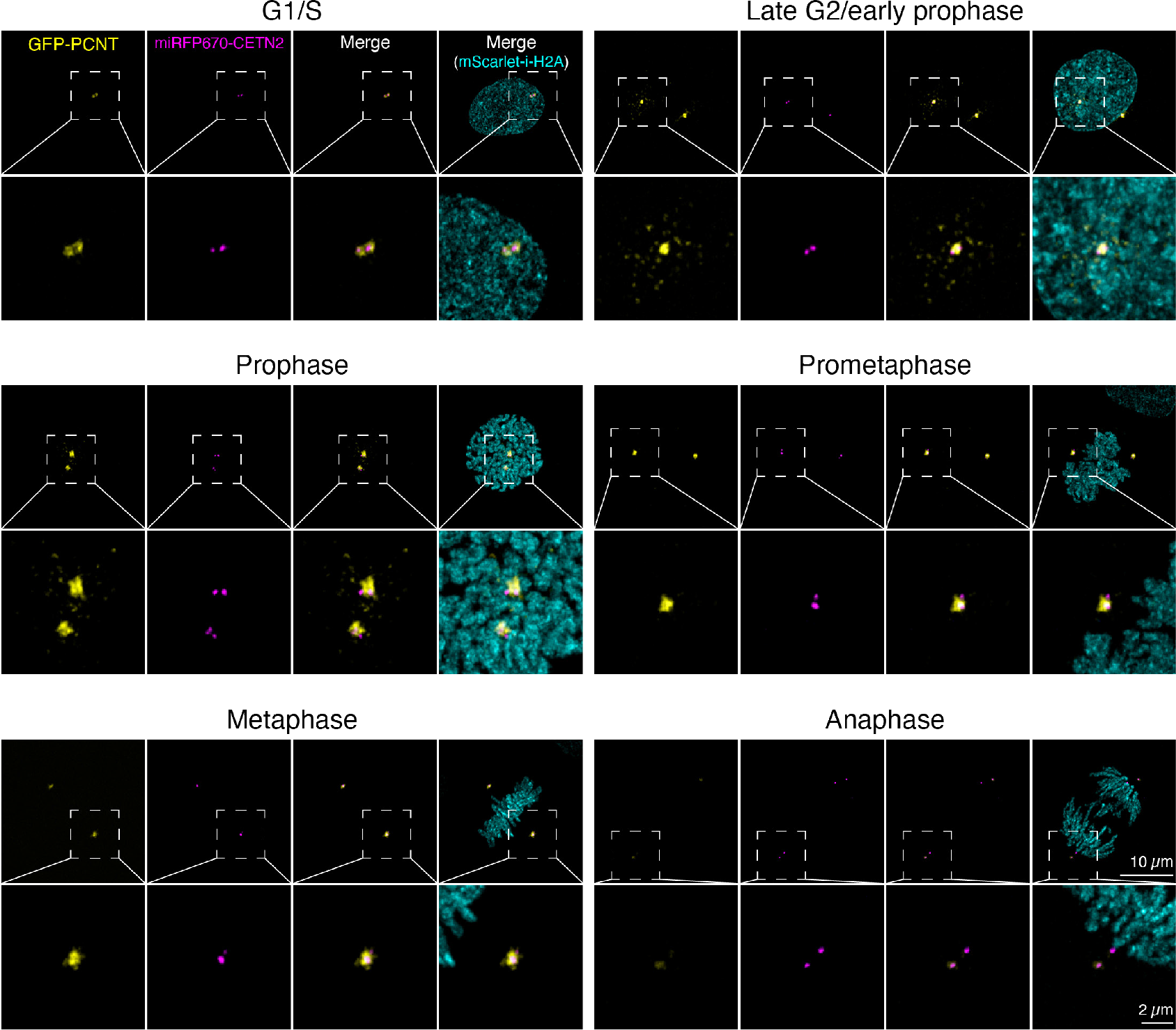
Representative confocal images of the data quantified in *Figure 1B*.

**Figure 1–Figure supplement 3.**
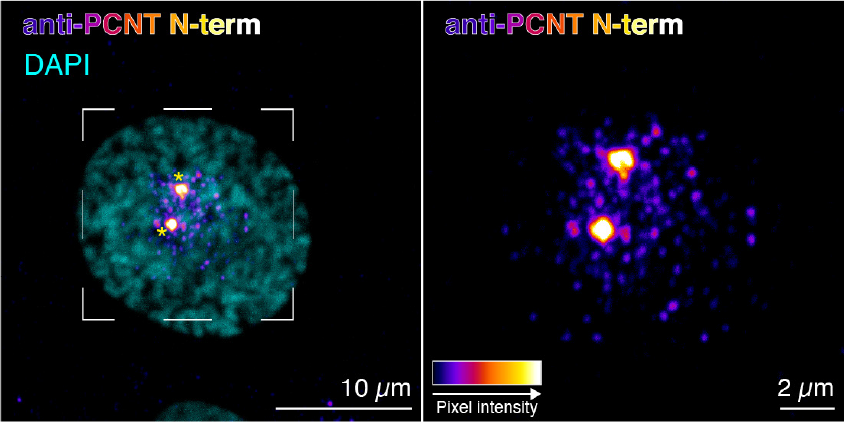
Immunofluorescence of a prophase RPE-1 cell using the anti-PCNT N-terminus antibody (Abcam, ab4448) to detect endogenous PCNT proteins. Asterisks denote the centrosomes. Similar results were obtained from three biological replicates.

**Figure 1–Figure supplement 4.**
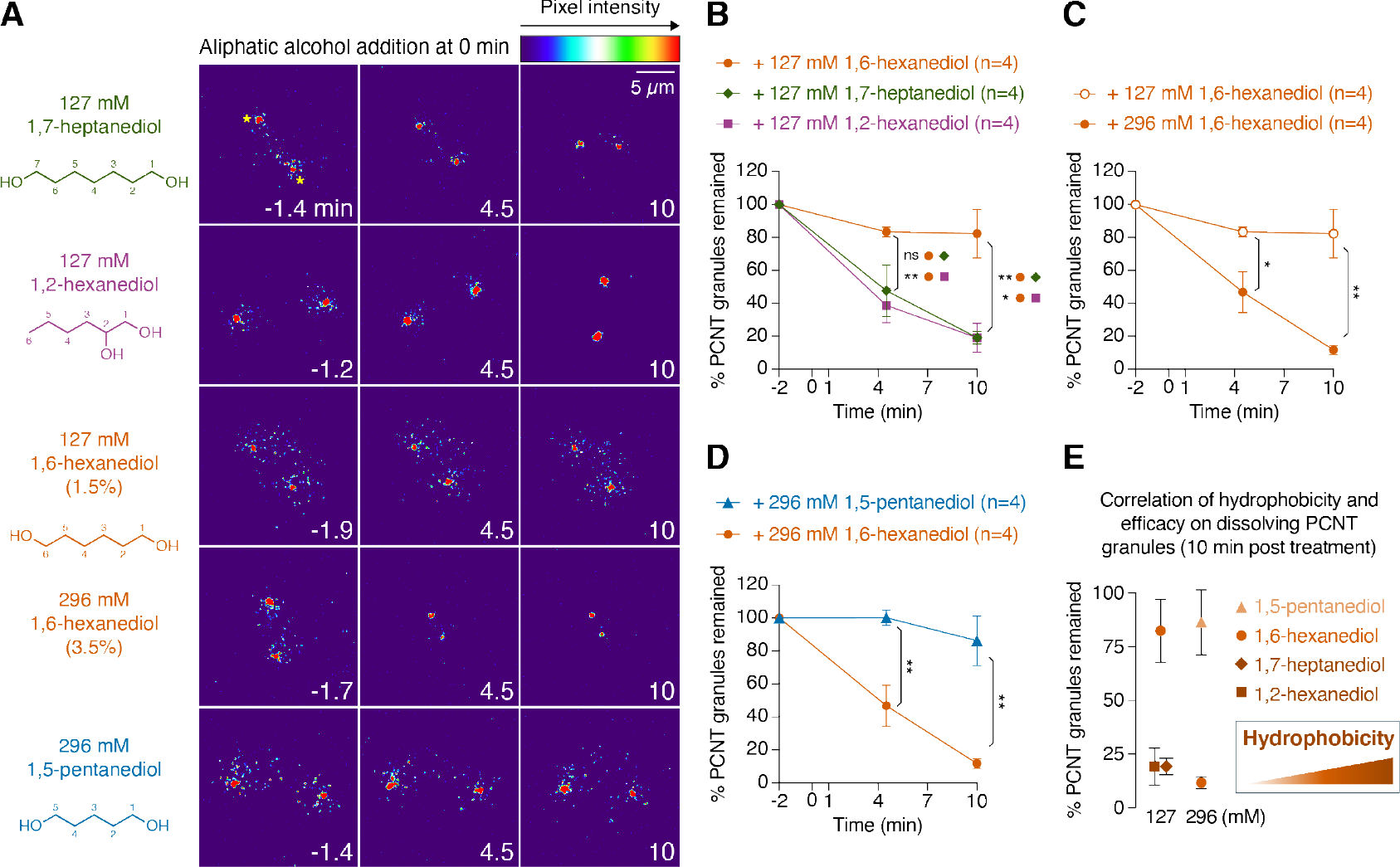
(**A**) Time-lapse micrographs of late G2/early M cells showing peri-centrosomal GFP-PCNT granules before and after acute treatments of various aliphatic alcohols at 127 or 296 mM. Note that 296 mM (or 3.5%) of 1.6-hexanediol was also the concentration used in the experiment shown in ***Figure 1C, C***’. Time 0 is the time of aliphatic alcohol addition. Two large red dots in each frame are the centrosomes depicted, for example, by asterisks in time −1.4 min on the top panel (127 mM 1,7-heptanediol). (**B-E**) Quantification of the results from the experiments shown in (A). After treatments of various aliphatic alcohols at 127 mM or 296 mM, percent of GFP-PCNT granules remained as the function of time was plotted and represented as mean ± SEM from two to three biological replicates. The results after 10 min of treatment were summarized in (E). The relative hydrophobicity of different aliphatic alcohols is represented as an orange gradient; the darker the color, the higher the hydrophobicity roughly is. The total number of cells analyzed for each condition is indicated. p-values were determined by thte Student’s t-test (two-tailed). *: p<0.05, **: p<0.01, ns: not significant.

**Figure 1–Figure supplement 5.**
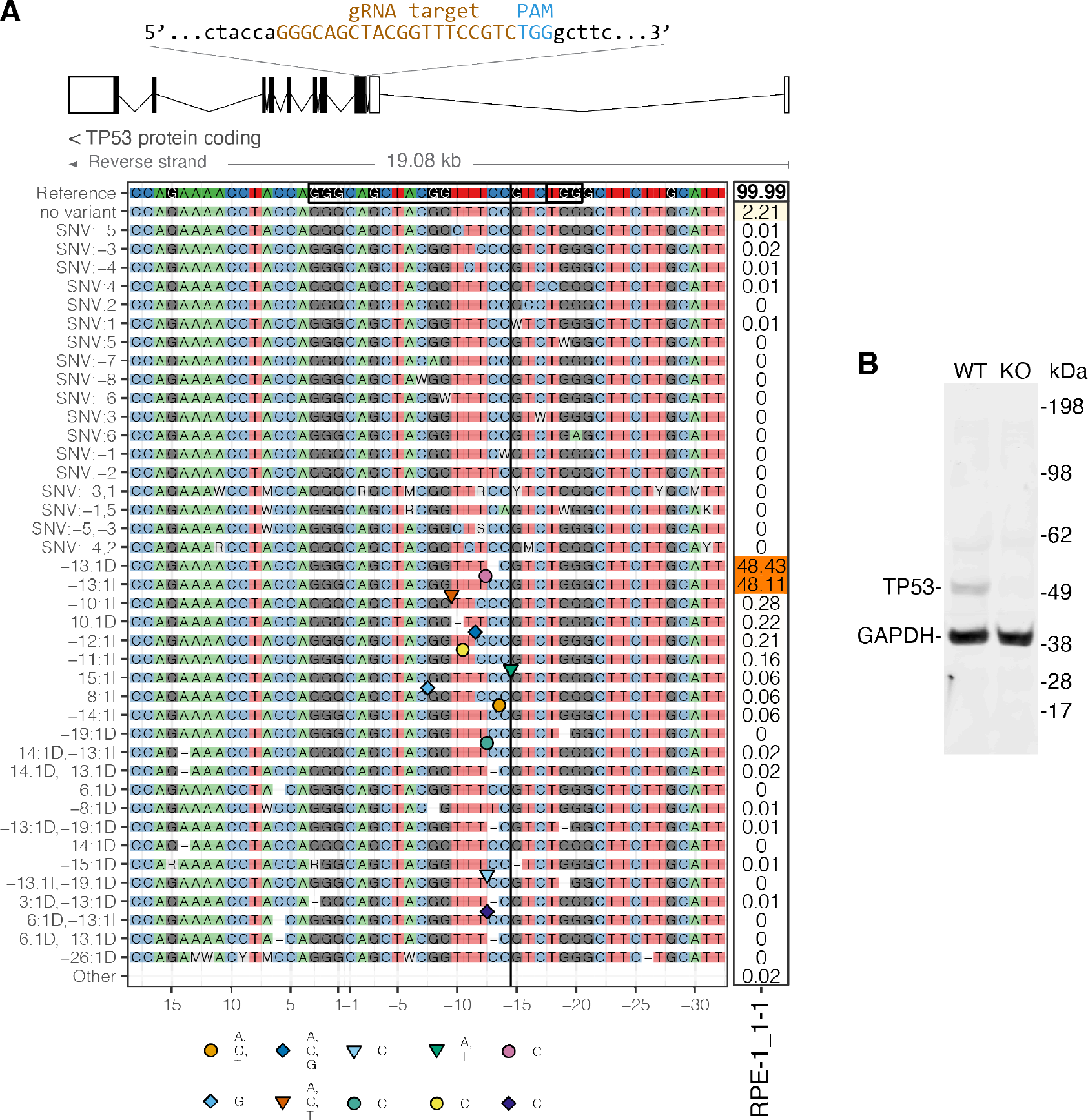
(**A**) Schematic to show the gRNA target to disrupt *TP53* in RPE1 cells. Illumina sequencing confirmed that one of the CRISPR-edited RPE-1 cell lines, RPE-1_1-1, has frameshift mutations at the gRNA target site in both *TP53* alleles (a 1-bp deletion and a 1-bp insertion). Sequencing data were analyzed and illustrated by an R-based toolkit, CrispRVariants (***Lindsay et al., 2016***). (**B**) Western blot analysis confirmed the loss of TP53 protein in RPE-1_1-1 cells. Anti-GAPDH staining served as the loading control. WT: parental RPE-1 cells; KO: RPE-1_1-1 cells.

**Figure 2–Figure supplement 1.**
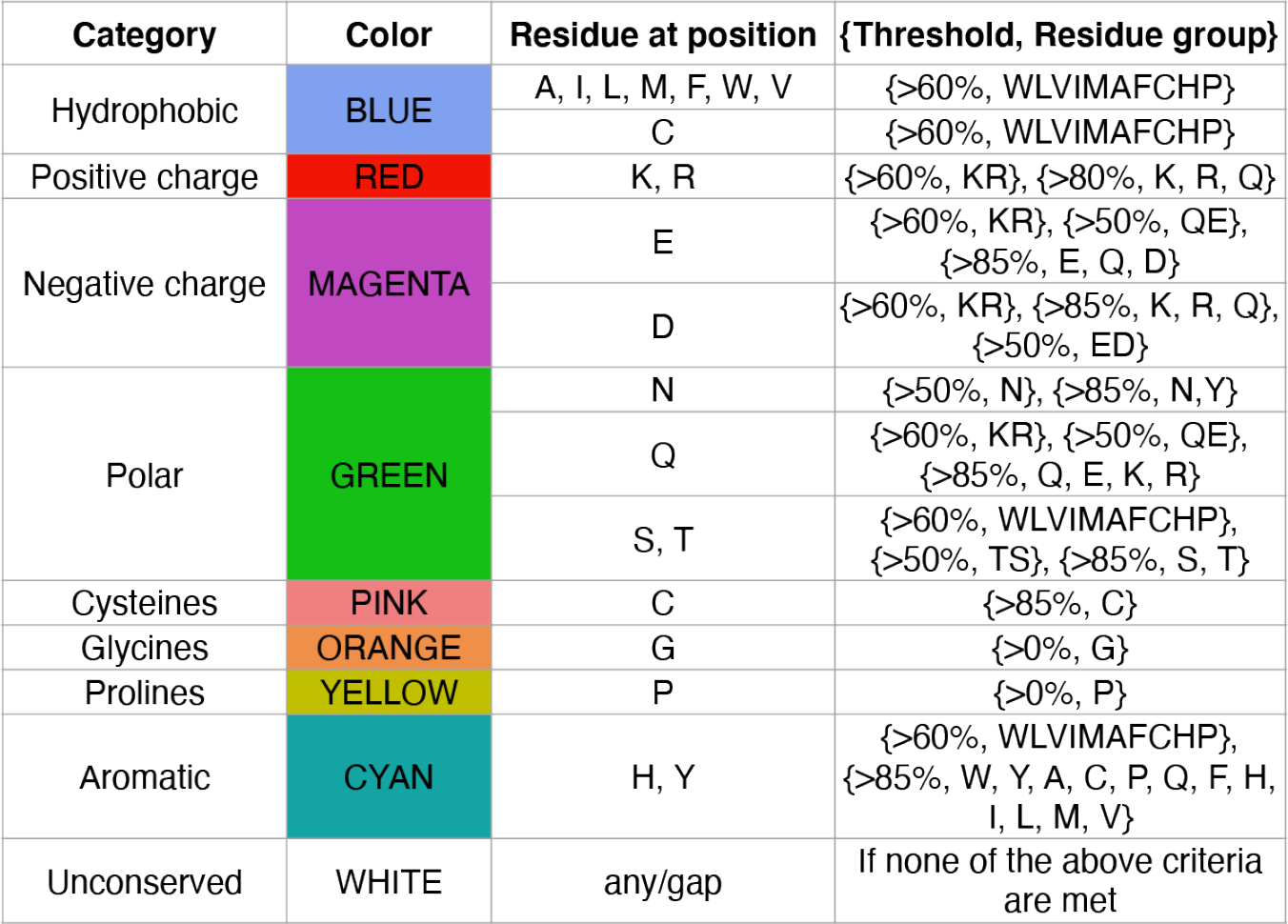
Clustal X coloring scheme in Jalview (***Waterhouse et al., 2009***) used in the multispecies alignments in *Figure 2A*.

**Figure 2–Figure supplement 2.**
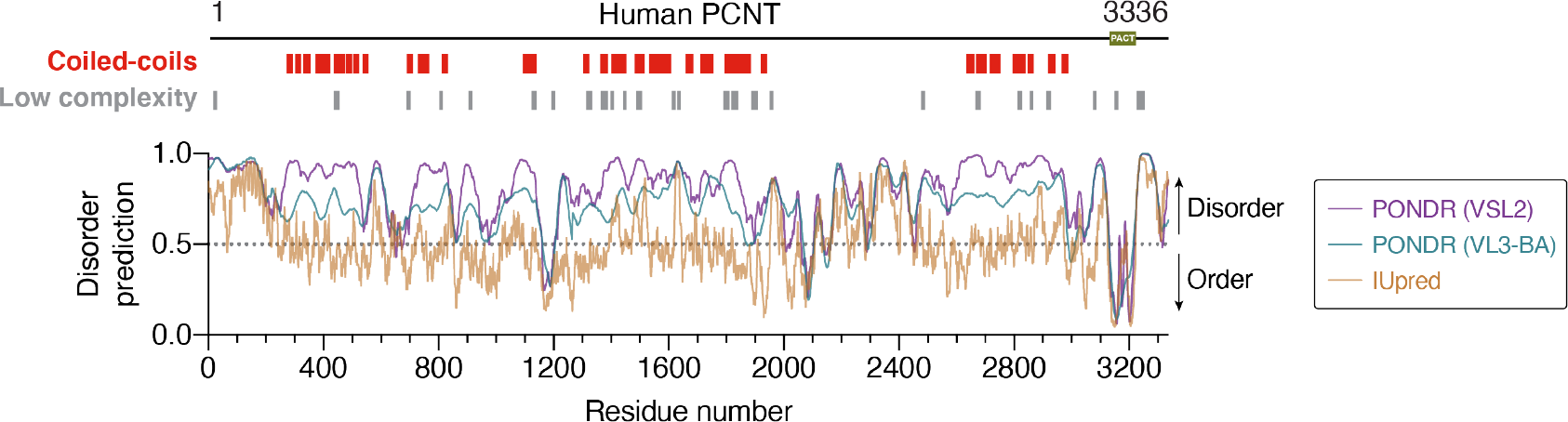
Predicted coiled-coils (CCs) and low-complexity regions (LCRs) of human PCNT aligned with disorder prediction results from three disorder prediction algorithms—PONDR VSL2, PONDR VL3-BA, and IUpred (***Peng et al., 2006***, ***2005***; ***Dosztanyi et al., 2005***).

**Figure 2–Figure supplement 3.**
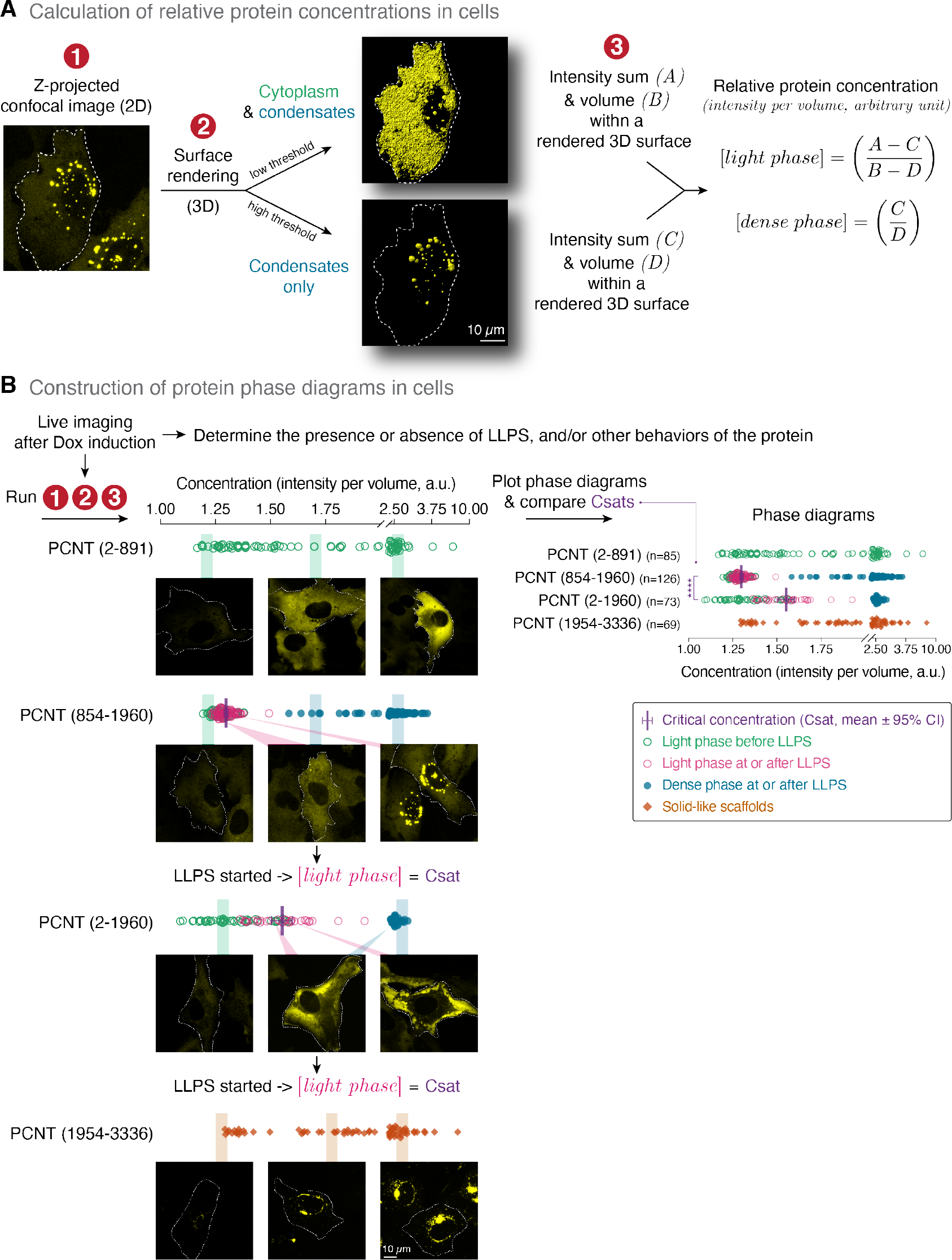
Worklow of determining the relative protein concentration and critical concentration (Csat) for phase separation in live cells (related to *Figure 2E*). (**A**) Schematic depicts the rendering of z-stack confocal images into 3D voxels—either as the combined cytoplasmic and condensate volumes or as the condensate volumes only—using the Imaris software. The intensity sum and the volume of the rendered 3D voxels were then measured and used to calculate the relative protein concentrations in the light and dense phases. (**B**) Live cell imaging was used to determine the presence or absence of LLPS after Dox-induced protein expression. The relative protein concentration was calculated using the workflow described in (A). Representative confocal images were shown. The Csat is the concentration of the light phase when LLPS just started.

**Figure 3–Figure supplement 1.**
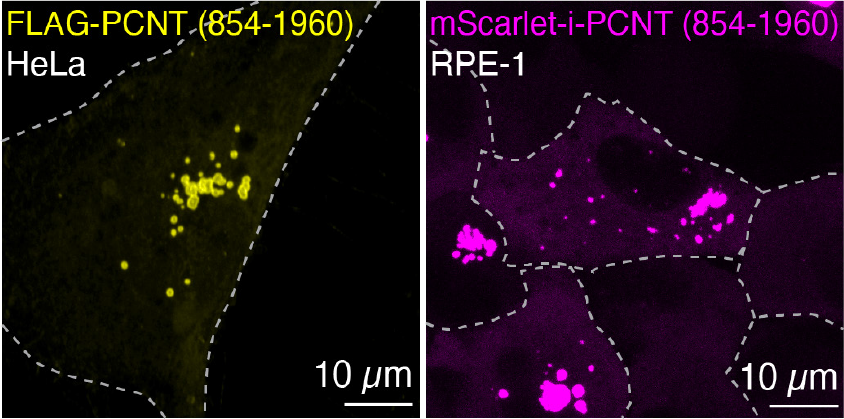
(**A**) Anti-PCNT N-terminus immunofluorescence of HeLa cells transiently expressing FLAG-PCNT (854-1960). Similar results were obtained from two biological replicates. (**B**) mScarlet-i-PCNT (854-1960) condensates in live RPE-1 cells. Dashed lines delineate the cell boundaries. Similar results were obtained from three biological replicates.

**Figure 3–Figure supplement 2.**
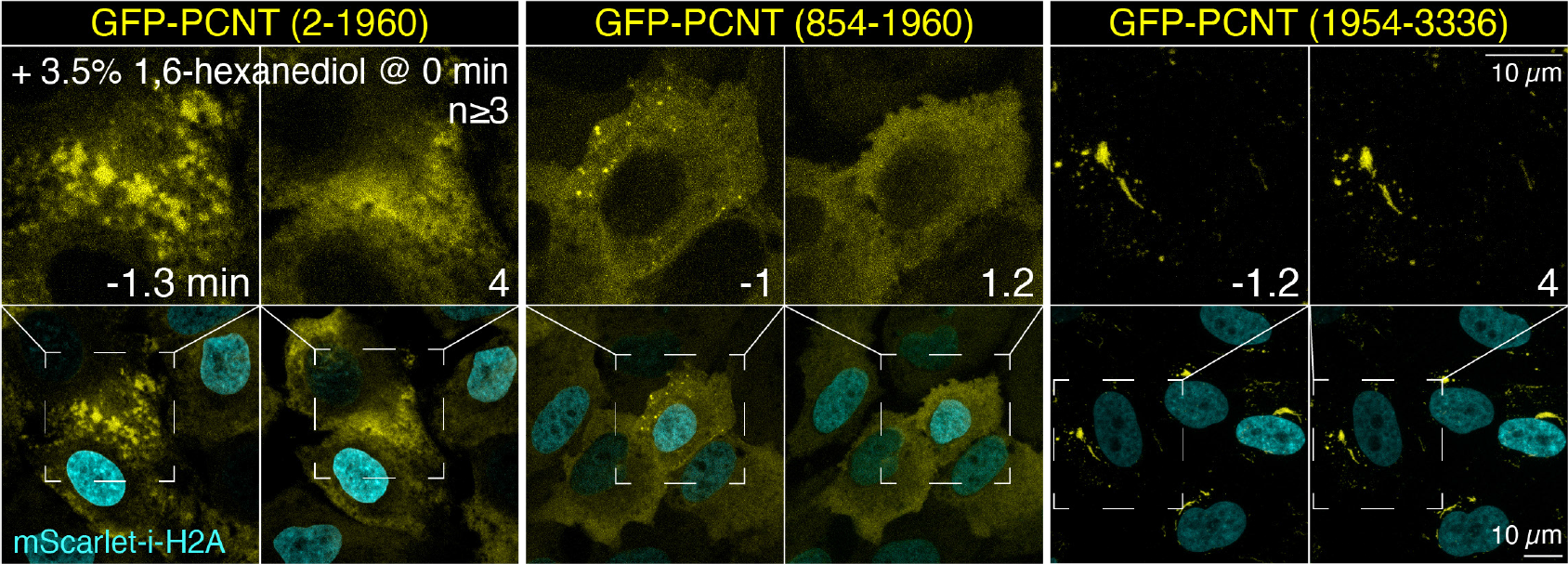
Representative time-lapse images of RPE-1 cells expressing GFP-PCNT (2-1960), GFP-PCNT (854-1960) condensates, or GFP-PCNT (1954-3336) scaffolds before and after the acute treatment of 3.5%/296 mM 1,6-hexanediol at time 0. Note that only the phase-separated, dynamic GFP-PCNT (2-1960) and GFP-PCNT (854-1960) condensates—but not the solid-like GFP-PCNT (1954-3336) scaffolds—were dissolved acutely by 1,6-hexanediol. Similar results were obtained from more than three biological replicates.

**Figure 5–Figure supplement 1.**
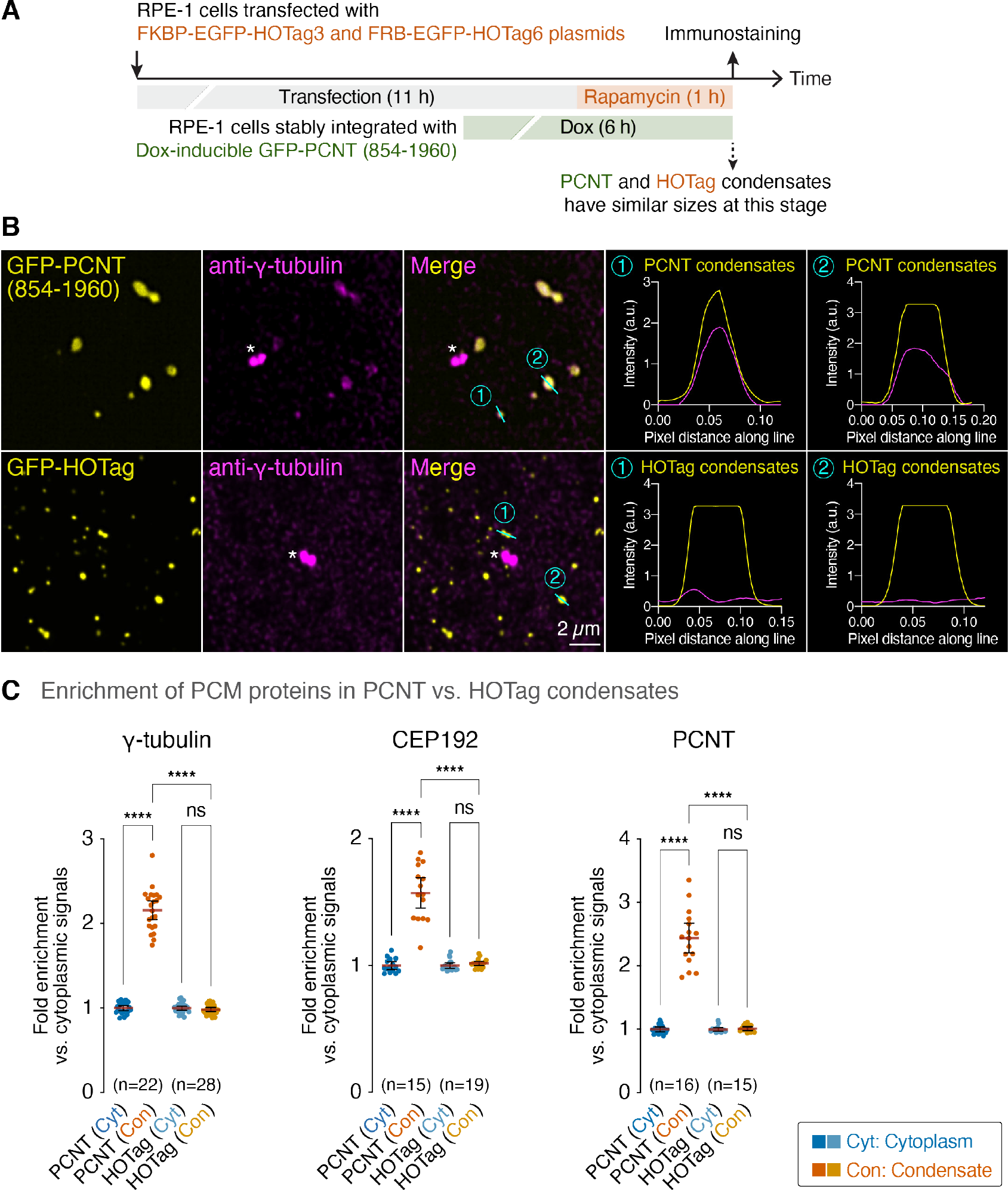
(**A**) Schematic of the recruitment assay to compare the ability of GFP-PCNT (854-1960) and GFP-HOTag condensates to recruit endogenous PCM proteins. Note that the expression of GFP-PCNT (854-1960) was only induced for 6 h (instead of 24 h in Figure 5 so that both PCNT (854-1960) and HOTag condensates would be similar in size. (**B**) Representative images of anti-*y*-tubulin immunofluorescence of GFP-PCNT (854-1960) and GFP-HOTag condensates. Right: The line plots of the selected regions that contain the condensates (cyan lines). Asterisks denote the centrosomes. a.u., arbitrary unit. (**C**) Fold enrichment of fluorescence signals in the PCNT or HOTag condensate relative to those in the cytoplasm was quantified. Data are mean ± 95% CI. n, number of cells analyzed from two, two, and one biological replicates for *y*-tubulin, CEP192, and PCNT, respectively. p-values were determined by one-way ANOVA. ****: p<0.0001; ns, not significant.

**Figure 5–Figure supplement 2.**
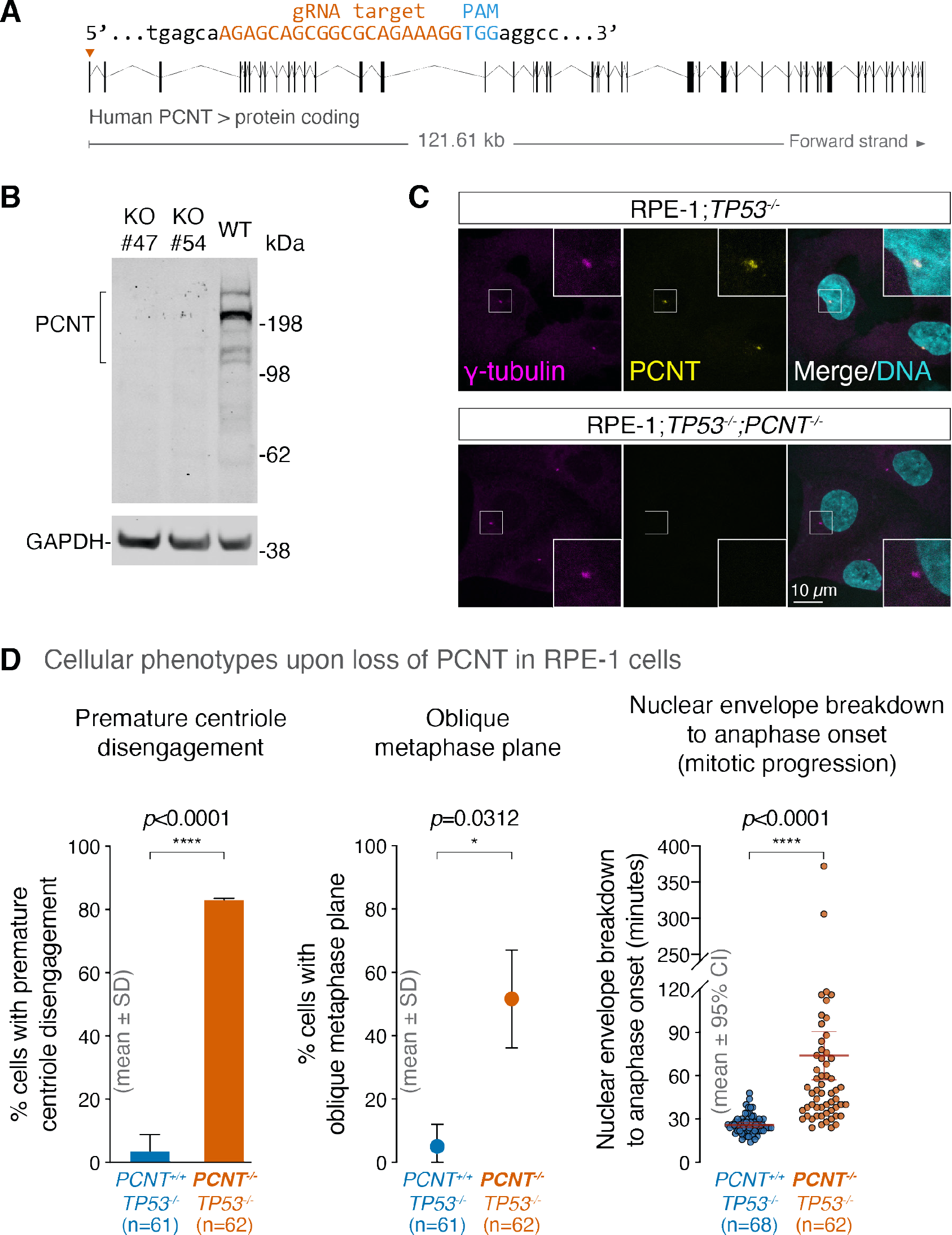
(**A**) Schematic to show the gRNA target to disrupt *PCNT* in RPE1 cells. (**B**) Western blot analysis confirmed the loss of PCNT protein in two *PCNT* knockout cell lines (KO#47 and KO#54). Anti-GAPDH staining served as the loading control. WT: parental *TP53^-/-^* RPE-1 cells, RPE-1_1-1. (**C**) Loss of PCNT signals at the centrosome of a *PCNT* knockout cell line (#47) was also confirmed by anti-PCNT immunostaining. (**D**) The *TP53^-/-^;PCNT^-/-^* RPE-1 cells (e.g., KO#47 shown here) showed several cellular defects, including premature centriole disengagement, oblique metaphase plane, and prolonged mitotic progression from nuclear envelope breakdown to anaphase onset.

**Figure 5–Figure supplement 3.**
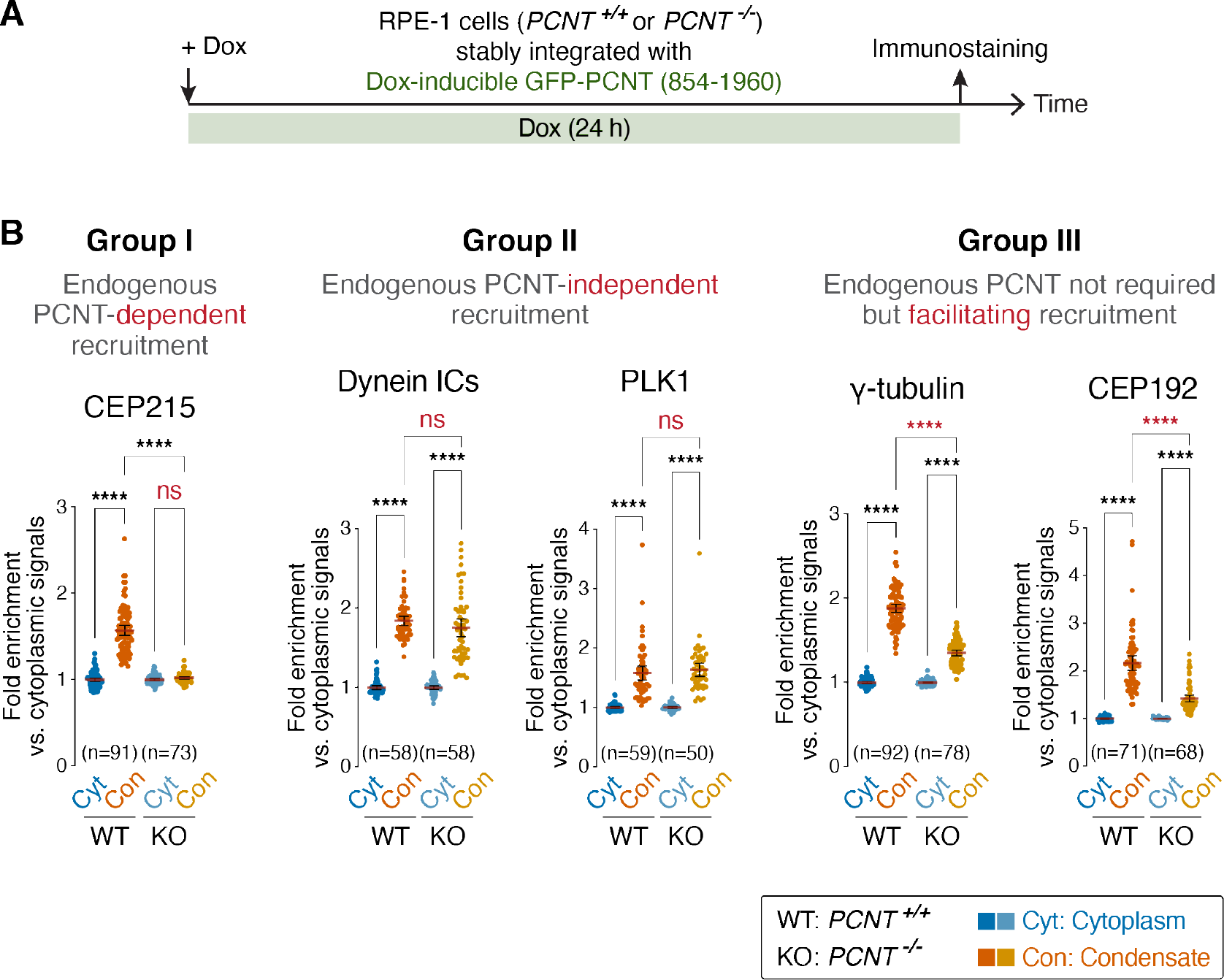
(**A**) Schematic of the recruitment assay to show the timeline of Dox induction and immunostaining. (**B**) Fold enrichment of fluorescence signals in the PCNT (854-1960) condensate relative to those in the cytoplasm in the presence or absence of endogenous PCNT was quantified. Note that endogenous PCNT may (Group I) or may not (Group II) be required for the PCNT (854-1960) condensate to recruit endogenous PCM components and clients. In addition, endogenous PCNT may not be required for but can facilitate the recruitment of certain PCM components (Group III). Data are mean ± 95% CI. n, number of cells analyzed from two biological replicates. p-values were determined by one-way ANOVA. ****: p<0.0001; ns, not significant.

**Figure 6–Figure supplement 1.**
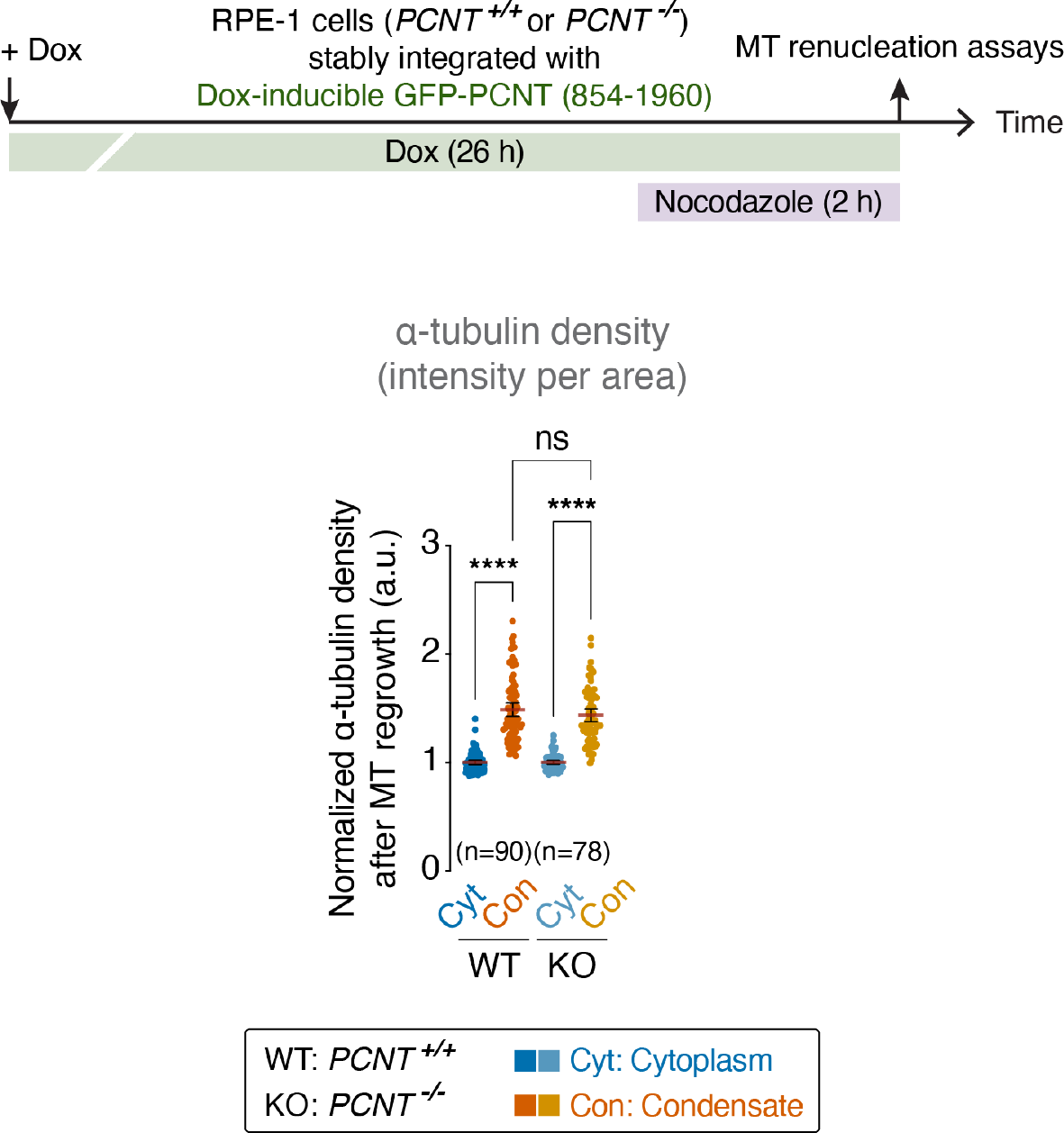
Quantification of *a*-tubulin density (intensity per area) in GFP-PCNT (854-1960) condensates (Con) and in the surrounding cytoplasm (Cyt) during MT renucleation in the control (WT) or *PCNT* knockout (KO) cells. Data are mean ± 95% CI. n, number of condensates analyzed from two biological replicates. The p-value was determined by one-way ANOVA. a.u., arbitrary unit.

**Figure 6–Figure supplement 2.**
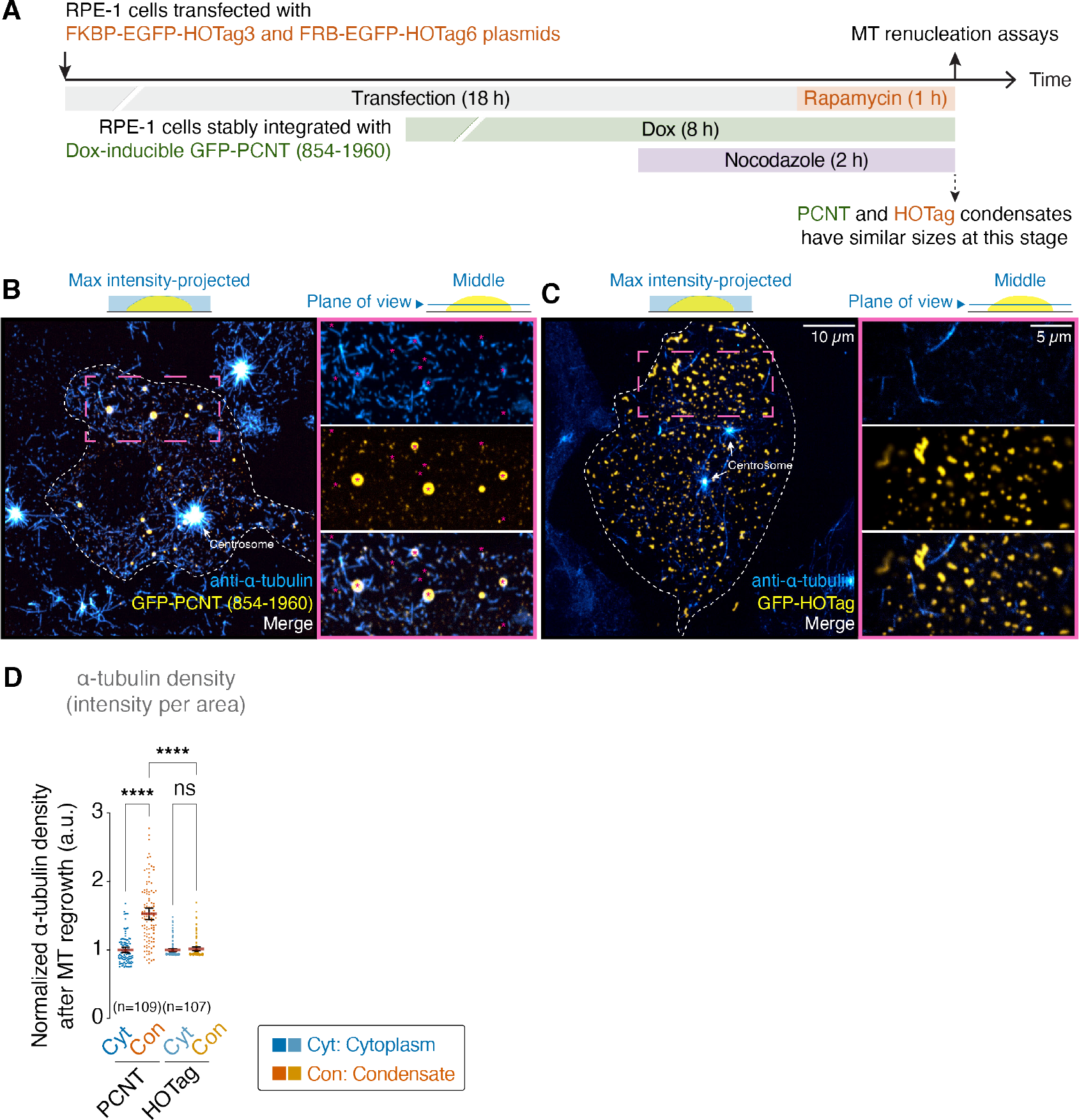
(**A**) Schematic of the MT renucleation assay to compare the ability of GFP-PCNT (854-1960) and GFP-HOTag condensates to nucleate MTs. (**B, C**) Anti-*a*-tubulin immunofluorescence of the cells containing GFP-PCNT (854-1960) (B) or GFP-HOTag (C) condensates after MT renucleation in maximum intensity-projected and single optical section views. Note that MTs were renucleated from some small PCNT condensates; many short MTs also appeared to originate from the very small PCNT condensates (B, some examples depicted by asterisks). However, MT renucleation was not observed in HOTag condensates (C). In contrast, MT renucleation was robust at the centrosome as expected. (**D**) Quantification of *a*-tubulin density (intensity per area) in GFP-PCNT (854-1960) and GFP-HOTag condensates (Con) and in the surrounding cytoplasm (Cyt) after MT renucleation. Data are mean ± 95% CI. n, number of condensates analyzed from three technical replicates. The p-value was determined by one-way ANOVA. ****: p<0.0001; ns, not significant. a.u., arbitrary unit.

**Figure 6–Figure supplement 3.**
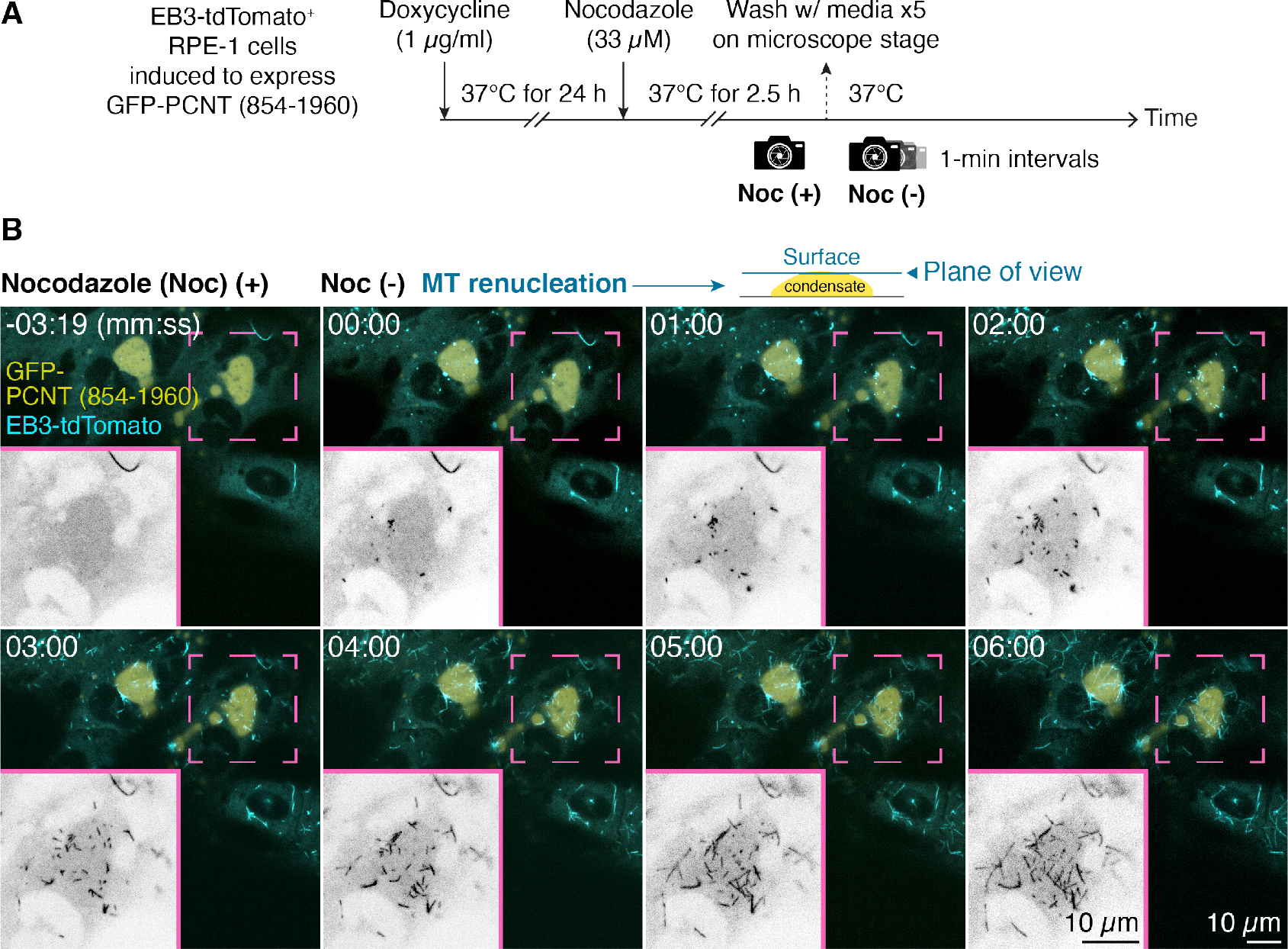
(**A**) Schematic of the MT renucleation assay in live cells. GFP-PCNT (854-1960) condensates in EB3-tdTomato-expressing RPE-1 cells were formed after Dox induction. MTs were then depolymerized by nocodazole (Noc). A pre-wash image was taken [Noc (+)], followed by time-lapse imaging at 1-min intervals after nocodazole was washed out on the microscope stage to follow MT renucleation [Noc (-)]. (**B**) Single optical sections of time-lapse micrographs of EB3-tdTomato-labeled MT plus ends on the surface of a GFP-PCNT (854-1960) condensate during MT renucleation. Time 0 was the time immediately after nocodazole was washed out (also see ***Figure 6– video 1***). Insets show the EB3-tdTomato channel only. Similar results were obtained from three biological replicates.

## References

Alvarez-Rodrigo I, Steinacker TL, Saurya S, Conduit PT, Baumbach J, Novak ZA, Aydogan MG, Wainman A, Raff JW. Evidence that a positive feedback loop drives centrosome maturation in fly embryos. Elife. 2019; 8:DOI: 10.7554/eLife.50130. https://www.ncbi.nlm.nih.gov/pubmed/31498081, doi: 10.7554/eLife.50130.

Andersen JS, Wilkinson CJ, Mayor T, Mortensen P, Nigg EA, Mann M. Proteomic characterization of the human centrosome by protein correlation profiling. Nature. 2003; 426(6966):570–4. http://www.ncbi.nlm.nih.gov/pubmed/14654843, doi: 10.1038/nature02166.

Anitha A, Nakamura K, Yamada K, Iwayama Y, Toyota T, Takei N, Iwata Y, Suzuki K, Sekine Y, Matsuzaki H, Kawai M, Thanseem I, Miyoshi K, Katayama T, Matsuzaki S, Baba K, Honda A, Hattori T, Shimizu S, Kumamoto N, et al. Association studies and gene expression analyses of the DISC1-interacting molecules, pericentrin 2 (PCNT2) and DISC1-binding zinc finger protein (DBZ), with schizophrenia and with bipolar disorder. Am J Med Genet B Neuropsychiatr Genet. 2009; 150B(7):967–76. https://www.ncbi.nlm.nih.gov/pubmed/19191256, doi: 10.1002/ajmg.b.30926.

Anurag M, Singh GP, Dash D. Location of disorder in coiled coil proteins is influenced by its biological role and subcellular localization: a GO-based study on human proteome. Mol Biosyst. 2012; 8(1):346–52. https://www.ncbi.nlm.nih.gov/pubmed/22027861, doi: 10.1039/c1mb05210a.

Asherie N. Protein crystallization and phase diagrams. Methods. 2004; 34(3):266–72. https://www.ncbi.nlm.nih.gov/pubmed/15325646, doi: 10.1016/j.ymeth.2004.03.028.

Atkins JD, Boateng SY, Sorensen T, McGuffin LJ. Disorder Prediction Methods, Their Applicability to Different Protein Targets and Their Usefulness for Guiding Experimental Studies. Int J Mol Sci. 2015; 16(8):19040–54. https://www.ncbi.nlm.nih.gov/pubmed/26287166, doi: 10.3390/ijms160819040.

Banani SF, Lee HO, Hyman AA, Rosen MK. Biomolecular condensates: organizers of cellular biochemistry. Nat Rev Mol Cell Biol. 2017; 18(5):285–298. https://www.ncbi.nlm.nih.gov/pubmed/28225081, doi: 10.1038/nrm.2017.7.

Barr AR, Gergely F. Aurora-A: the maker and breaker of spindle poles. J Cell Sci. 2007; 120(Pt 17):2987–96. https://www.ncbi.nlm.nih.gov/pubmed/17715155, doi: 10.1242/jcs.013136.

Berdnik D, Knoblich JA. Drosophila Aurora-A is required for centrosome maturation and actin-dependent asymmetric protein localization during mitosis. Curr Biol. 2002; 12(8):640–7. https://www.ncbi.nlm.nih.gov/pubmed/11967150, doi: 10.1016/s0960-9822(02)00766-2.

Berry J, Weber SC, Vaidya N, Haataja M, Brangwynne CP. RNA transcription modulates phase transition-driven nuclear body assembly. Proc Natl Acad Sci U S A. 2015; 112(38):E5237–45. https://www.ncbi.nlm.nih.gov/pubmed/26351690, doi: 10.1073/pnas.1509317112.

Bindels DS, Haarbosch L, van Weeren L, Postma M, Wiese KE, Mastop M, Aumonier S, Gotthard G, Royant A, Hink MA, Gadella J T W. mScarlet: a bright monomeric red fluorescent protein for cellular imaging. Nat Methods. 2017; 14(1):53–56. https://www.ncbi.nlm.nih.gov/pubmed/27869816, doi: 10.1038/nmeth.4074.

Boeynaems S, Alberti S, Fawzi NL, Mittag T, Polymenidou M, Rousseau F, Schymkowitz J, Shorter J, Wolozin B, Van Den Bosch L, Tompa P, Fuxreiter M. Protein Phase Separation: A New Phase in Cell Biology. Trends Cell Biol. 2018; 28(6):420–435. https://www.ncbi.nlm.nih.gov/pubmed/29602697, doi: 10.1016/j.tcb.2018.02.004.

Boke E, Ruer M, Wuhr M, Coughlin M, Lemaitre R, Gygi SP, Alberti S, Drechsel D, Hyman AA, Mitchison TJ. Amyloid-like Self-Assembly of a Cellular Compartment. Cell. 2016; 166(3):637–650. https://www.ncbi.nlm.nih.gov/pubmed/27471966, doi: 10.1016/j.cell.2016.06.051.

Chinen T, Yamazaki K, Hashimoto K, Fujii K, Watanabe K, Takeda Y, Yamamoto S, Nozaki Y, Tsuchiya Y, Takao D, Kitagawa D. Centriole and PCM cooperatively recruit CEP192 to spindle poles to promote bipolar spindle assembly. J Cell Biol. 2021; 220(2). https://www.ncbi.nlm.nih.gov/pubmed/33443571, doi: 10.1083/jcb.202006085.

Conduit PT, Brunk K, Dobbelaere J, Dix CI, Lucas EP, Raff JW. Centrioles regulate centrosome size by controlling the rate of Cnn incorporation into the PCM. Curr Biol. 2010; 20(24):2178–86. https://www.ncbi.nlm.nih.gov/pubmed/21145741, doi: 10.1016/j.cub.2010.11.011.

Conduit PT, Feng Z, Richens JH, Baumbach J, Wainman A, Bakshi SD, Dobbelaere J, Johnson S, Lea SM, Raff JW. The centrosome-specific phosphorylation of Cnn by Polo/Plk1 drives Cnn scaffold assembly and centrosome maturation. Dev Cell. 2014; 28(6):659–69. https://www.ncbi.nlm.nih.gov/pubmed/24656740, doi: 10.1016/j.devcel.2014.02.013.

Conduit PT, Richens JH, Wainman A, Holder J, Vicente CC, Pratt MB, Dix CI, Novak ZA, Dobbie IM, Schermelleh L, Raff JW. A molecular mechanism of mitotic centrosome assembly in Drosophila. Elife. 2014; 3:e03399. https://www.ncbi.nlm.nih.gov/pubmed/25149451, doi: 10.7554/eLife.03399.

Conduit PT, Wainman A, Raff JW. Centrosome function and assembly in animal cells. Nat Rev Mol Cell Biol. 2015; 16(10):611–24. https://www.ncbi.nlm.nih.gov/pubmed/26373263, doi: 10.1038/nrm4062.

Delaval B, Doxsey SJ. Pericentrin in cellular function and disease. J Cell Biol. 2010; 188(2):181–90. https://www.ncbi.nlm.nih.gov/pubmed/19951897, doi: 10.1083/jcb.200908114.

Dobbelaere J, Josue F, Suijkerbuijk S, Baum B, Tapon N, Raff J. A genome-wide RNAi screen to dissect centriole duplication and centrosome maturation in Drosophila. PLoS Biol. 2008; 6(9):e224. https://www.ncbi.nlm.nih.gov/pubmed/18798690, doi: 10.1371/journal.pbio.0060224.

Dosztanyi Z, Csizmok V, Tompa P, Simon I. The pairwise energy content estimated from amino acid composition discriminates between folded and intrinsically unstructured proteins. J Mol Biol. 2005; 347(4):827–39. https://www.ncbi.nlm.nih.gov/pubmed/15769473, doi: 10.1016/j.jmb.2005.01.071.

Doxsey S. Re-evaluating centrosome function. Nat Rev Mol Cell Biol. 2001; 2(9):688–98. http://www.ncbi.nlm.nih.gov/pubmed/11533726, doi: 10.1038/35089575.

Dull T, Zufferey R, Kelly M, Mandel RJ, Nguyen M, Trono D, Naldini L. A third-generation lentivirus vector with a conditional packaging system. J Virol. 1998; 72(11):8463–71. https://www.ncbi.nlm.nih.gov/pubmed/9765382.

Edgar RC. MUSCLE: multiple sequence alignment with high accuracy and high throughput. Nucleic Acids Res. 2004; 32(5):1792–7. https://www.ncbi.nlm.nih.gov/pubmed/15034147, doi: 10.1093/nar/gkh340.

Elbaum-Garfinkle S, Kim Y, Szczepaniak K, Chen CC, Eckmann CR, Myong S, Brangwynne CP. The disordered P granule protein LAF-1 drives phase separation into droplets with tunable viscosity and dynamics. Proc Natl Acad Sci U S A. 2015; 112(23):7189–94. https://www.ncbi.nlm.nih.gov/pubmed/26015579, doi: 10.1073/pnas.1504822112.

Fang X, Wang L, Ishikawa R, Li Y, Fiedler M, Liu F, Calder G, Rowan B, Weigel D, Li P, Dean C. Arabidopsis FLL2 promotes liquid-liquid phase separation of polyadenylation complexes. Nature. 2019; 569(7755):265–269. https://www.ncbi.nlm.nih.gov/pubmed/31043738, doi: 10.1038/s41586-019-1165-8.

Felix MA, Antony C, Wright M, Maro B. Centrosome assembly in vitro: role of gamma-tubulin recruitment in Xenopus sperm aster formation. J Cell Biol. 1994; 124(1-2):19–31. https://www.ncbi.nlm.nih.gov/pubmed/8294501, doi: 10.1083/jcb.124.1.19.

Feng Z, Caballe A, Wainman A, Johnson S, Haensele AFM, Cottee MA, Conduit PT, Lea SM, Raff JW. Structural Basis for Mitotic Centrosome Assembly in Flies. Cell. 2017; 169(6):1078–1089 e13. https://www.ncbi.nlm.nih.gov/pubmed/28575671, doi: 10.1016/j.cell.2017.05.030.

Fernandes N, Buchan JR. RPS28B mRNA acts as a scaffold promoting cis-translational interaction of proteins driving P-body assembly. Nucleic Acids Res. 2020; 48(11):6265–6279. https://www.ncbi.nlm.nih.gov/pubmed/32396167, doi: 10.1093/nar/gkaa352.

Firestone AJ, Weinger JS, Maldonado M, Barlan K, Langston LD, O’Donnell M, Gelfand VI, Kapoor TM, Chen JK. Small-molecule inhibitors of the AAA+ ATPase motor cytoplasmic dynein. Nature. 2012; 484(7392):125–9. https://www.ncbi.nlm.nih.gov/pubmed/22425997, doi: 10.1038/nature10936.

Fu J, Glover DM. Structured illumination of the interface between centriole and peri-centriolar material. Open Biol. 2012; 2(8):120104. http://www.ncbi.nlm.nih.gov/pubmed/22977736, doi: 10.1098/rsob.120104.

Galati DF, Sullivan KD, Pham AT, Espinosa JM, Pearson CG. Trisomy 21 Represses Cilia Formation and Function. Dev Cell. 2018; 46(5):641–650 e6. https://www.ncbi.nlm.nih.gov/pubmed/30100262, doi: 10.1016/j.devcel.2018.07.008.

Gibson DG, Young L, Chuang RY, Venter JC, Hutchison r C A, Smith HO. Enzymatic assembly of DNA molecules up to several hundred kilobases. Nat Methods. 2009; 6(5):343–5. https://www.ncbi.nlm.nih.gov/pubmed/19363495, doi: 10.1038/nmeth.1318.

Gillingham AK, Munro S. The PACT domain, a conserved centrosomal targeting motif in the coiled-coil proteins AKAP450 and pericentrin. EMBO Rep. 2000; 1(6):524–9. http://www.ncbi.nlm.nih.gov/pubmed/11263498, doi: 10.1093/embo-reports/kvd105.

Goshima G, Wollman R, Goodwin SS, Zhang N, Scholey JM, Vale RD, Stuurman N. Genes required for mitotic spindle assembly in Drosophila S2 cells. Science. 2007; 316(5823):417–21. https://www.ncbi.nlm.nih.gov/pubmed/17412918, doi: 10.1126/science.1141314.

Grifith E, Walker S, Martin CA, Vagnarelli P, Stiff T, Vernay B, Al Sanna N, Saggar A, Hamel B, Earnshaw WC, Jeggo PA, Jackson AP, O’Driscoll M. Mutations in pericentrin cause Seckel syndrome with defective ATR-dependent DNA damage signaling. Nat Genet. 2008; 40(2):232–6. https://www.ncbi.nlm.nih.gov/pubmed/18157127, doi: 10.1038/ng.2007.80.

Grigoryan G, Keating AE. Structural specificity in coiled-coil interactions. Curr Opin Struct Biol. 2008; 18(4):477– 83. https://www.ncbi.nlm.nih.gov/pubmed/18555680, doi: 10.1016/j.sbi.2008.04.008.

Grigoryan G, Kim YH, Acharya R, Axelrod K, Jain RM, Willis L, Drndic M, Kikkawa JM, DeGrado WF. Computational design of virus-like protein assemblies on carbon nanotube surfaces. Science. 2011; 332(6033):1071–6. https://www.ncbi.nlm.nih.gov/pubmed/21617073, doi: 10.1126/science.1198841.

Gupta GD, Pelletier L. Centrosome Biology: Polymer-Based Centrosome Maturation. Curr Biol. 2017; 27(17):R836–R839. https://www.ncbi.nlm.nih.gov/pubmed/28898644, doi: 10.1016/j.cub.2017.07.036.

Hamill DR, Severson AF, Carter JC, Bowerman B. Centrosome maturation and mitotic spindle assembly in C. elegans require SPD-5, a protein with multiple coiled-coil domains. Dev Cell. 2002; 3(5):673–84. https://www.ncbi.nlm.nih.gov/pubmed/12431374.

Hannak E, Kirkham M, Hyman AA, Oegema K. Aurora-A kinase is required for centrosome maturation in Caenorhabditis elegans. J Cell Biol. 2001; 155(7):1109–16. https://www.ncbi.nlm.nih.gov/pubmed/11748251, doi: 10.1083/jcb.200108051.

Hannak E, Oegema K, Kirkham M, Gonczy P, Habermann B, Hyman AA. The kinetically dominant assembly pathway for centrosomal asters in Caenorhabditis elegans is gamma-tubulin dependent. J Cell Biol. 2002; 157(4):591–602. https://www.ncbi.nlm.nih.gov/pubmed/12011109, doi: 10.1083/jcb.200202047.

Haren L, Stearns T, Luders J. Plk1-dependent recruitment of gamma-tubulin complexes to mitotic centrosomes involves multiple PCM components. PLoS One. 2009; 4(6):e5976. https://www.ncbi.nlm.nih.gov/pubmed/19543530, doi: 10.1371/journal.pone.0005976.

Harris CR, Millman KJ, van der Walt SJ, Gommers R, Virtanen P, Cournapeau D, Wieser E, Taylor J, Berg S, Smith NJ, Kern R, Picus M, Hoyer S, van Kerkwijk MH, Brett M, Haldane A, del Río JF, Wiebe M, Peterson P, Gérard-Marchant P, et al. Array programming with NumPy. Nature. 2020; 585(7825):357–362. https://doi.org/10.1038/s41586-020-2649-2, doi: 10.1038/s41586-020-2649-2.

Hennig S, Kong G, Mannen T, Sadowska A, Kobelke S, Blythe A, Knott GJ, Iyer KS, Ho D, Newcombe EA, Hosoki K, Goshima N, Kawaguchi T, Hatters D, Trinkle-Mulcahy L, Hirose T, Bond CS, Fox AH. Prion-like domains in RNA binding proteins are essential for building subnuclear paraspeckles. J Cell Biol. 2015; 210(4):529–39. https://www.ncbi.nlm.nih.gov/pubmed/26283796, doi: 10.1083/jcb.201504117.

Hoing S, Yeh TY, Baumann M, Martinez NE, Habenberger P, Kremer L, Drexler HCA, Kuchler P, Reinhardt P, Choidas A, Zischinsky ML, Zischinsky G, Nandini S, Ledray AP, Ketcham SA, Reinhardt L, Abo-Rady M, Glatza M, King SJ, Nussbaumer P, et al. Dynarrestin, a Novel Inhibitor of Cytoplasmic Dynein. Cell Chem Biol. 2018; 25(4):357–369 e6. https://www.ncbi.nlm.nih.gov/pubmed/29396292, doi: 10.1016/j.chembiol.2017.12.014.

Holehouse AS, Pappu RV. Functional Implications of Intracellular Phase Transitions. Biochemistry. 2018; 57(17):2415–2423. https://www.ncbi.nlm.nih.gov/pubmed/29323488, doi: 10.1021/acs.biochem.7b01136.

Huang PS, Oberdorfer G, Xu C, Pei XY, Nannenga BL, Rogers JM, DiMaio F, Gonen T, Luisi B, Baker D. High thermodynamic stability of parametrically designed helical bundles. Science. 2014; 346(6208):481–485. https://www.ncbi.nlm.nih.gov/pubmed/25342806, doi: 10.1126/science.1257481.

Hunter JD. Matplotlib: A 2D Graphics Environment. Computing in Science and Engg. 2007 May; 9(3):90–95. https://doi.org/10.1109/MCSE.2007.55, doi: 10.1109/MCSE.2007.55.

Hutchins JR, Toyoda Y, Hegemann B, Poser I, Heriche JK, Sykora MM, Augsburg M, Hudecz O, Buschhorn BA, Bulkescher J, Conrad C, Comartin D, Schleiffer A, Sarov M, Pozniakovsky A, Slabicki MM, Schloissnig S, Stein-macher I, Leuschner M, Ssykor A, et al. Systematic analysis of human protein complexes identifies chromosome segregation proteins. Science. 2010; 328(5978):593–9. https://www.ncbi.nlm.nih.gov/pubmed/20360068, doi: 10.1126/science.1181348.

Hyman AA, Weber CA, Julicher F. Liquid-liquid phase separation in biology. Annu Rev Cell Dev Biol. 2014; 30:39–58. https://www.ncbi.nlm.nih.gov/pubmed/25288112, doi: 10.1146/annurev-cellbio-100913-013325.

Jain A, Vale RD. RNA phase transitions in repeat expansion disorders. Nature. 2017; 546(7657):243–247. https://www.ncbi.nlm.nih.gov/pubmed/28562589, doi: 10.1038/nature22386.

Jao LE, Akef A, Wente SR. A role for Gle1, a regulator of DEAD-box RNA helicases, at centrosomes and basal bodies. Mol Biol Cell. 2017; 28(1):120–127. https://www.ncbi.nlm.nih.gov/pubmed/28035044, doi: 10.1091/mbc.E16-09-0675.

Jeng R, Stearns T. Gamma-tubulin complexes: size does matter. Trends Cell Biol. 1999; 9(9):339–42. https://www.ncbi.nlm.nih.gov/pubmed/10461186.

Jiang F, Taylor DW, Chen JS, Kornfeld JE, Zhou K, Thompson AJ, Nogales E, Doudna JA. Structures of a CRISPR-Cas9 R-loop complex primed for DNA cleavage. Science. 2016; 351(6275):867–71. https://www.ncbi.nlm.nih.gov/pubmed/26841432, doi: 10.1126/science.aad8282.

Jiang F, Zhou K, Ma L, Gressel S, Doudna JA. A Cas9-guide RNA complex preorganized for target DNA recognition. Science. 2015; 348(6242):1477–81. https://www.ncbi.nlm.nih.gov/pubmed/26113724, doi: 10.1126/science.aab1452.

Jiang X, Brust-Mascher I, Jao LE. Three-dimensional Reconstruction and Quantification of Proteins and mRNAs at the Single-cell Level in Cultured Cells. Bio Protoc. 2019; 9(16):e3330. https://www.ncbi.nlm.nih.gov/pubmed/33654837, doi: 10.21769/BioProtoc.3330.

Joshi HC, Palacios MJ, McNamara L, Cleveland DW. Gamma-tubulin is a centrosomal protein required for cell cycle-dependent microtubule nucleation. Nature. 1992; 356(6364):80–3. https://www.ncbi.nlm.nih.gov/pubmed/1538786, doi: 10.1038/356080a0.

Joukov V, Walter JC, De Nicolo A. The Cep192-organized aurora A-Plk1 cascade is essential for centrosome cycle and bipolar spindle assembly. Mol Cell. 2014; 55(4):578–91. https://www.ncbi.nlm.nih.gov/pubmed/25042804, doi: 10.1016/j.molcel.2014.06.016.

Kalderon D, Roberts BL, Richardson WD, Smith AE. A short amino acid sequence able to specify nuclear location. Cell. 1984; 39(3 Pt 2):499–509. https://www.ncbi.nlm.nih.gov/pubmed/6096007, doi: 10.1016/0092-8674(84)90457-4.

Kato M, Han TW, Xie S, Shi K, Du X, Wu LC, Mirzaei H, Goldsmith EJ, Longgood J, Pei J, Grishin NV, Frantz DE, Schneider JW, Chen S, Li L, Sawaya MR, Eisenberg D, Tycko R, McKnight SL. Cell-free formation of RNA granules: low complexity sequence domains form dynamic fibers within hydrogels. Cell. 2012; 149(4):753–67. https://www.ncbi.nlm.nih.gov/pubmed/22579281, doi: 10.1016/j.cell.2012.04.017.

Kemp CA, Kopish KR, Zipperlen P, Ahringer J, O’Connell KF. Centrosome maturation and duplication in C. elegans require the coiled-coil protein SPD-2. Dev Cell. 2004; 6(4):511–23. https://www.ncbi.nlm.nih.gov/pubmed/15068791.

Khodjakov A, Rieder CL. The sudden recruitment of gamma-tubulin to the centrosome at the onset of mitosis and its dynamic exchange throughout the cell cycle, do not require microtubules. J Cell Biol. 1999; 146(3):585–96. https://www.ncbi.nlm.nih.gov/pubmed/10444067.

Kim S, Rhee K. Importance of the CEP215-pericentrin interaction for centrosome maturation during mitosis. PLoS One. 2014; 9(1):e87016. https://www.ncbi.nlm.nih.gov/pubmed/24466316, doi: 10.1371/journal.pone.0087016.

Kim SI, Oceguera-Yanez F, Sakurai C, Nakagawa M, Yamanaka S, Woltjen K. Inducible Transgene Expression in Human iPS Cells Using Versatile All-in-One piggyBac Transposons. Methods Mol Biol. 2016; 1357:111–31. https://www.ncbi.nlm.nih.gov/pubmed/26025620, doi: 10.1007/7651_2015_251.

Kinoshita K, Noetzel TL, Pelletier L, Mechtler K, Drechsel DN, Schwager A, Lee M, Raff JW, Hyman AA. Aurora A phosphorylation of TACC3/maskin is required for centrosome-dependent microtubule assembly in mitosis. J Cell Biol. 2005; 170(7):1047–55. https://www.ncbi.nlm.nih.gov/pubmed/16172205, doi: 10.1083/jcb.200503023.

Knop M, Schiebel E. Spc98p and Spc97p of the yeast gamma-tubulin complex mediate binding to the spindle pole body via their interaction with Spc110p. EMBO J. 1997; 16(23):6985–95. https://www.ncbi.nlm.nih.gov/pubmed/9384578, doi: 10.1093/emboj/16.23.6985.

Kroschwald S, Maharana S, Mateju D, Malinovska L, Nuske E, Poser I, Richter D, Alberti S. Promiscuous interactions and protein disaggregases determine the material state of stress-inducible RNP granules. Elife. 2015; 4:e06807. https://www.ncbi.nlm.nih.gov/pubmed/26238190, doi: 10.7554/eLife.06807.

Kroschwald S, Maharana S, Simon A. Hexanediol: a chemical probe to investigate the material properties of membrane-less compartments. Matters. 2017; doi: 10.19185/matters.201702000010.

Kwan KM, Fujimoto E, Grabher C, Mangum BD, Hardy ME, Campbell DS, Parant JM, Yost HJ, Kanki JP, Chien CB. The Tol2kit: a multisite gateway-based construction kit for Tol2 transposon transgenesis constructs. Dev Dyn. 2007; 236(11):3088–99. https://www.ncbi.nlm.nih.gov/pubmed/17937395, doi: 10.1002/dvdy.21343.

Langdon EM, Qiu Y, Ghanbari Niaki A, McLaughlin GA, Weidmann CA, Gerbich TM, Smith JA, Crutchley JM, Termini CM, Weeks KM, Myong S, Gladfelter AS. mRNA structure determines specificity of a polyQ-driven phase separation. Science. 2018; 360(6391):922–927. https://www.ncbi.nlm.nih.gov/pubmed/29650703, doi: 10.1126/science.aar7432.

Larson AG, Elnatan D, Keenen MM, Trnka MJ, Johnston JB, Burlingame AL, Agard DA, Redding S, Narlikar GJ. Liquid droplet formation by HP1alpha suggests a role for phase separation in heterochromatin. Nature. 2017; 547(7662):236–240. https://www.ncbi.nlm.nih.gov/pubmed/28636604, doi: 10.1038/nature22822.

Lawo S, Hasegan M, Gupta GD, Pelletier L. Subdiffraction imaging of centrosomes reveals higher-order organizational features of pericentriolar material. Nat Cell Biol. 2012; 14(11):1148–58. http://www.ncbi.nlm.nih.gov/pubmed/23086237, doi: 10.1038/ncb2591.

Lee CS, Putnam A, Lu T, He S, Ouyang JPT, Seydoux G. Recruitment of mRNAs to P granules by condensation with intrinsically-disordered proteins. Elife. 2020; 9. https://www.ncbi.nlm.nih.gov/pubmed/31975687, doi: 10.7554/eLife.52896.

Lee K, Rhee K. PLK1 phosphorylation of pericentrin initiates centrosome maturation at the onset of mitosis. J Cell Biol. 2011; 195(7):1093–101. https://www.ncbi.nlm.nih.gov/pubmed/22184200, doi: 10.1083/jcb.201106093.

Li Y, Brown JH, Reshetnikova L, Blazsek A, Farkas L, Nyitray L, Cohen C. Visualization of an unstable coiled coil from the scallop myosin rod. Nature. 2003; 424(6946):341–5. https://www.ncbi.nlm.nih.gov/pubmed/12867988, doi: 10.1038/nature01801.

Lin S, Staahl BT, Alla RK, Doudna JA. Enhanced homology-directed human genome engineering by controlled timing of CRISPR/Cas9 delivery. Elife. 2014; 3:e04766. https://www.ncbi.nlm.nih.gov/pubmed/25497837, doi: 10.7554/eLife.04766.

Lin TC, Neuner A, Schlosser YT, Scharf AN, Weber L, Schiebel E. Cell-cycle dependent phosphorylation of yeast pericentrin regulates gamma-TuSC-mediated microtubule nucleation. Elife. 2014; 3:e02208. https://www.ncbi.nlm.nih.gov/pubmed/24842996, doi: 10.7554/eLife.02208.

Lin Y, Mori E, Kato M, Xiang S, Wu L, Kwon I, McKnight SL. Toxic PR Poly-Dipeptides Encoded by the C9orf72 Repeat Expansion Target LC Domain Polymers. Cell. 2016; 167(3):789–802 e12. https://www.ncbi.nlm.nih.gov/pubmed/27768897, doi: 10.1016/j.cell.2016.10.003.

Lin Y, Protter DS, Rosen MK, Parker R. Formation and Maturation of Phase-Separated Liquid Droplets by RNA-Binding Proteins. Mol Cell. 2015; 60(2):208–19. https://www.ncbi.nlm.nih.gov/pubmed/26412307, doi: 10.1016/j.molcel.2015.08.018.

Lindsay H, Burger A, Biyong B, Felker A, Hess C, Zaugg J, Chiavacci E, Anders C, Jinek M, Mosimann C, Robinson MD. CrispRVariants charts the mutation spectrum of genome engineering experiments. Nat Biotechnol. 2016; 34(7):701–2. https://www.ncbi.nlm.nih.gov/pubmed/27404876, doi: 10.1038/nbt.3628.

Livingstone CD, Barton GJ. Protein sequence alignments: a strategy for the hierarchical analysis of residue conservation. Comput Appl Biosci. 1993; 9(6):745–56. https://www.ncbi.nlm.nih.gov/pubmed/8143162, doi: 10.1093/bioinformatics/9.6.745.

Lu Y, Wu T, Gutman O, Lu H, Zhou Q, Henis YI, Luo K. Phase separation of TAZ compartmentalizes the transcription machinery to promote gene expression. Nat Cell Biol. 2020; https://www.ncbi.nlm.nih.gov/pubmed/32203417, doi: 10.1038/s41556-020-0485-0.

Lupas A, Van Dyke M, Stock J. Predicting coiled coils from protein sequences. Science. 1991; 252(5009):1162–4. https://www.ncbi.nlm.nih.gov/pubmed/2031185, doi: 10.1126/science.252.5009.1162.

Ma W, Zheng G, Xie W, Mayr C. In vivo reconstitution finds multivalent RNA-RNA interactions as drivers of mesh-like condensates. Elife. 2021; 10. https://www.ncbi.nlm.nih.gov/pubmed/33650968, doi: 10.7554/eLife.64252.

Maharana S, Wang J, Papadopoulos DK, Richter D, Pozniakovsky A, Poser I, Bickle M, Rizk S, Guillen-Boixet J, Franzmann TM, Jahnel M, Marrone L, Chang YT, Sterneckert J, Tomancak P, Hyman AA, Alberti S. RNA buffers the phase separation behavior of prion-like RNA binding proteins. Science. 2018; 360(6391):918– 921. https://www.ncbi.nlm.nih.gov/pubmed/29650702, doi: 10.1126/science.aar7366.

Mahen R, Venkitaraman AR. Pattern formation in centrosome assembly. Curr Opin Cell Biol. 2012; 24(1):14–23. https://www.ncbi.nlm.nih.gov/pubmed/22245706, doi: 10.1016/j.ceb.2011.12.012.

Mali P, Yang L, Esvelt KM, Aach J, Guell M, DiCarlo JE, Norville JE, Church GM. RNA-guided human genome engineering via Cas9. Science. 2013; 339(6121):823–6. http://www.ncbi.nlm.nih.gov/pubmed/23287722, doi: 10.1126/science.1232033.

Martinez-Campos M, Basto R, Baker J, Kernan M, Raff JW. The Drosophila pericentrin-like protein is essential for cilia/flagella function, but appears to be dispensable for mitosis. J Cell Biol. 2004; 165(5):673–83. http://www.ncbi.nlm.nih.gov/pubmed/15184400, doi: 10.1083/jcb.200402130.

Meng L, Park JE, Kim TS, Lee EH, Park SY, Zhou M, Bang JK, Lee KS. Bimodal Interaction of Mammalian Polo-Like Kinase 1 and a Centrosomal Scaffold, Cep192, in the Regulation of Bipolar Spindle Formation. Mol Cell Biol. 2015; 35(15):2626–40. https://www.ncbi.nlm.nih.gov/pubmed/26012549, doi: 10.1128/MCB.00068-15.

Mennella V, Agard DA, Huang B, Pelletier L. Amorphous no more: subdiffraction view of the pericentriolar material architecture. Trends Cell Biol. 2014; 24(3):188–97. https://www.ncbi.nlm.nih.gov/pubmed/24268653, doi: 10.1016/j.tcb.2013.10.001.

Mennella V, Keszthelyi B, McDonald KL, Chhun B, Kan F, Rogers GC, Huang B, Agard DA. Subdiffraction-resolution fluorescence microscopy reveals a domain of the centrosome critical for pericentriolar material organization. Nat Cell Biol. 2012; 14(11):1159–68. http://www.ncbi.nlm.nih.gov/pubmed/23086239, doi: 10.1038/ncb2597.

Merriam EB, Millette M, Lumbard DC, Saengsawang W, Fothergill T, Hu X, Ferhat L, Dent EW. Synaptic regulation of microtubule dynamics in dendritic spines by calcium, F-actin, and drebrin. J Neurosci. 2013; 33(42):16471– 82. https://www.ncbi.nlm.nih.gov/pubmed/24133252, doi: 10.1523/JNEUROSCI.0661-13.2013.

Mikule K, Delaval B, Kaldis P, Jurcyzk A, Hergert P, Doxsey S. Loss of centrosome integrity induces p38-p53-p21-dependent G1-S arrest. Nat Cell Biol. 2007; 9(2):160–70. https://www.ncbi.nlm.nih.gov/pubmed/17330329, doi: 10.1038/ncb1529.

Molliex A, Temirov J, Lee J, Coughlin M, Kanagaraj AP, Kim HJ, Mittag T, Taylor JP. Phase separation by low complexity domains promotes stress granule assembly and drives pathological fibrillization. Cell. 2015; 163(1):123–33. https://www.ncbi.nlm.nih.gov/pubmed/26406374, doi: 10.1016/j.cell.2015.09.015.

Moritz M, Braunfeld MB, Fung JC, Sedat JW, Alberts BM, Agard DA. Three-dimensional structural characterization of centrosomes from early Drosophila embryos. J Cell Biol. 1995; 130(5):1149–59. https://www.ncbi.nlm.nih.gov/pubmed/7657699, doi: 10.1083/jcb.130.5.1149.

Moritz M, Braunfeld MB, Sedat JW, Alberts B, Agard DA. Microtubule nucleation by gamma-tubulin-containing rings in the centrosome. Nature. 1995; 378(6557):638–40. https://www.ncbi.nlm.nih.gov/pubmed/8524401, doi: 10.1038/378638a0.

Moritz M, Zheng Y, Alberts BM, Oegema K. Recruitment of the gamma-tubulin ring complex to Drosophila salt-stripped centrosome scaffolds. J Cell Biol. 1998; 142(3):775–86. https://www.ncbi.nlm.nih.gov/pubmed/9700165, doi: 10.1083/jcb.142.3.775.

Neumann B, Walter T, Heriche JK, Bulkescher J, Erfle H, Conrad C, Rogers P, Poser I, Held M, Liebel U, Cetin C, Sieckmann F, Pau G, Kabbe R, Wunsche A, Satagopam V, Schmitz MH, Chapuis C, Gerlich DW, Schneider R, et al. Phenotypic profiling of the human genome by time-lapse microscopy reveals cell division genes. Nature. 2010; 464(7289):721–7. https://www.ncbi.nlm.nih.gov/pubmed/20360735, doi: 10.1038/nature08869.

Nott TJ, Petsalaki E, Farber P, Jervis D, Fussner E, Plochowietz A, Craggs TD, Bazett-Jones DP, Pawson T, Forman-Kay JD, Baldwin AJ. Phase transition of a disordered nuage protein generates environmentally responsive membraneless organelles. Mol Cell. 2015; 57(5):936–947. https://www.ncbi.nlm.nih.gov/pubmed/25747659, doi: 10.1016/j.molcel.2015.01.013.

Numata S, Iga J, Nakataki M, Tayoshi S, Tanahashi T, Itakura M, Ueno S, Ohmori T. Positive association of the pericentrin (PCNT) gene with major depressive disorder in the Japanese population. J Psychiatry Neurosci. 2009; 34(3):195–8. https://www.ncbi.nlm.nih.gov/pubmed/19448849.

Oakley BR, Oakley CE, Yoon Y, Jung MK. Gamma-tubulin is a component of the spindle pole body that is essential for microtubule function in Aspergillus nidulans. Cell. 1990; 61(7):1289–301. https://www.ncbi.nlm.nih.gov/pubmed/2194669, doi: 10.1016/0092-8674(90)90693-9.

Oegema K, Wiese C, Martin OC, Milligan RA, Iwamatsu A, Mitchison TJ, Zheng Y. Characterization of two related Drosophila gamma-tubulin complexes that differ in their ability to nucleate microtubules. J Cell Biol. 1999; 144(4):721–33. https://www.ncbi.nlm.nih.gov/pubmed/10037793, doi: 10.1083/jcb.144.4.721.

Palazzo RE, Vogel JM, Schnackenberg BJ, Hull DR, Wu X. Centrosome maturation. Curr Top Dev Biol. 2000; 49:449–70. https://www.ncbi.nlm.nih.gov/pubmed/11005031.

Patel A, Lee HO, Jawerth L, Maharana S, Jahnel M, Hein MY, Stoynov S, Mahamid J, Saha S, Franzmann TM, Pozniakovski A, Poser I, Maghelli N, Royer LA, Weigert M, Myers EW, Grill S, Drechsel D, Hyman AA, Alberti S. A Liquid-to-Solid Phase Transition of the ALS Protein FUS Accelerated by Disease Mutation. Cell. 2015; 162(5):1066–77. https://www.ncbi.nlm.nih.gov/pubmed/26317470, doi: 10.1016/j.cell.2015.07.047.

Patel SS, Belmont BJ, Sante JM, Rexach MF. Natively unfolded nucleoporins gate protein diffusion across the nuclear pore complex. Cell. 2007; 129(1):83–96. https://www.ncbi.nlm.nih.gov/pubmed/17418788, doi: 10.1016/j.cell.2007.01.044.

Peng K, Radivojac P, Vucetic S, Dunker AK, Obradovic Z. Length-dependent prediction of protein intrinsic disorder. BMC Bioinformatics. 2006; 7:208. https://www.ncbi.nlm.nih.gov/pubmed/16618368, doi: 10.1186/1471-2105-7-208.

Peng K, Vucetic S, Radivojac P, Brown CJ, Dunker AK, Obradovic Z. Optimizing long intrinsic disorder predictors with protein evolutionary information. J Bioinform Comput Biol. 2005; 3(1):35–60. https://www.ncbi.nlm.nih.gov/pubmed/15751111, doi: 10.1142/s0219720005000886.

Piehl M, Tulu US, Wadsworth P, Cassimeris L. Centrosome maturation: measurement of microtubule nucleation throughout the cell cycle by using GFP-tagged EB1. Proc Natl Acad Sci U S A. 2004; 101(6):1584–8. https://www.ncbi.nlm.nih.gov/pubmed/14747658, doi: 10.1073/pnas.0308205100.

Raff JW. Phase Separation and the Centrosome: A Fait Accompli? Trends Cell Biol. 2019; https://www.ncbi.nlm.nih.gov/pubmed/31076235, doi: 10.1016/j.tcb.2019.04.001.

Rauch A, Thiel CT, Schindler D, Wick U, Crow YJ, Ekici AB, van Essen AJ, Goecke TO, Al-Gazali L, Chrzanowska KH, Zweier C, Brunner HG, Becker K, Curry CJ, Dallapiccola B, Devriendt K, Dorfler A, Kinning E, Megarbane A, Meinecke P, et al. Mutations in the pericentrin (PCNT) gene cause primordial dwarfism. Science. 2008; 319(5864):816–9. http://www.ncbi.nlm.nih.gov/pubmed/18174396, doi: 10.1126/science.1151174.

Reback J, McKinney W, jbrockmendel, den Bossche JV, Augspurger T, Cloud P, gfyoung, Sinhrks, Klein A, Roeschke M, Tratner J, She C, Ayd W, Hawkins S, Petersen T, Schendel J, Hayden A, Garcia M, Jancauskas V, MomIsBestFriend, et al., pandas-dev/pandas: Pandas 1.0.0. Zenodo; 2020. https://doi.org/10.5281/zenodo.3630805, doi: 10.5281/zenodo.3630805.

Riback JA, Katanski CD, Kear-Scott JL, Pilipenko EV, Rojek AE, Sosnick TR, Drummond DA. Stress-Triggered Phase Separation Is an Adaptive, Evolutionarily Tuned Response. Cell. 2017; 168(6):1028–1040 e19. https://www.ncbi.nlm.nih.gov/pubmed/28283059, doi: 10.1016/j.cell.2017.02.027.

Ribbeck K, Gorlich D. The permeability barrier of nuclear pore complexes appears to operate via hydrophobic exclusion. EMBO J. 2002; 21(11):2664–71. https://www.ncbi.nlm.nih.gov/pubmed/12032079, doi: 10.1093/em-boj/21.11.2664.

Rieder CL, Borisy GG. The centrosome cycle in PtK2 cells: Asymmetric distribution and structural changes in the pericentriolar material. Biology of the Cell. 1982; 44:117–132.

Rog O, Kohler S, Dernburg AF. The synaptonemal complex has liquid crystalline properties and spatially regulates meiotic recombination factors. Elife. 2017; 6. https://www.ncbi.nlm.nih.gov/pubmed/28045371, doi: 10.7554/eLife.21455.

Romero CM, Páez MS, Miranda JA, Hernández DJ, Oviedo LE. Effect of temperature on the surface tension of diluted aqueous solutions of 1,2-hexanediol, 1,5-hexanediol, 1,6-hexanediol and 2,5-hexanediol. Fluid Phase Equilibria. 2007; 258(1):67–72. http://www.sciencedirect.com/science/article/pii/S0378381207002968, doi: https://doi.org/10.1016/j.fluid.2007.05.029.

Romero P, Obradovic Z, Dunker AK. Folding minimal sequences: the lower bound for sequence complexity of globular proteins. FEBS Lett. 1999; 462(3):363–7. https://www.ncbi.nlm.nih.gov/pubmed/10622726, doi: 10.1016/s0014-5793(99)01557-4.

Rose A, Meier I. Scaffolds, levers, rods and springs: diverse cellular functions of long coiled-coil proteins. Cell Mol Life Sci. 2004; 61(16):1996–2009. https://www.ncbi.nlm.nih.gov/pubmed/15316650, doi: 10.1007/s00018-004-4039-6.

Salisbury JL. Centrosomes: coiled-coils organize the cell center. Curr Biol. 2003; 13(3):R88–90. https://www.ncbi.nlm.nih.gov/pubmed/12573233, doi: 10.1016/s0960-9822(03)00033-2.

Sanders A, Chang K, Zhu X, Thoppil RJ, Holmes WR, Kaverina I. Nonrandom gamma-TuNA-dependent spatial pattern of microtubule nucleation at the Golgi. Mol Biol Cell. 2017; 28(23):3181–3192. https://www.ncbi.nlm.nih.gov/pubmed/28931596, doi: 10.1091/mbc.E17-06-0425.

Schindelin J, Arganda-Carreras I, Frise E, Kaynig V, Longair M, Pietzsch T, Preibisch S, Rueden C, Saalfeld S, Schmid B, Tinevez JY, White DJ, Hartenstein V, Eliceiri K, Tomancak P, Cardona A. Fiji: an open-source plat-form for biological-image analysis. Nat Methods. 2012; 9(7):676–82. https://www.ncbi.nlm.nih.gov/pubmed/22743772, doi: 10.1038/nmeth.2019.

Schmidt HB, Gorlich D. Nup98 FG domains from diverse species spontaneously phase-separate into particles with nuclear pore-like permselectivity. Elife. 2015; 4. https://www.ncbi.nlm.nih.gov/pubmed/25562883, doi: 10.7554/eLife.04251.

Schnackenberg BJ, Khodjakov A, Rieder CL, Palazzo RE. The disassembly and reassembly of functional centrosomes in vitro. Proc Natl Acad Sci U S A. 1998; 95(16):9295–300. https://www.ncbi.nlm.nih.gov/pubmed/9689074, doi: 10.1073/pnas.95.16.9295.

Schnackenberg BJ, Palazzo RE. Identification and function of the centrosome centromatrix. Biology of the Cell. 1999; 91(6):429–438. https://onlinelibrary.wiley.com/doi/abs/10.1111/j.1768-322X.1999.tb01098.x, doi: https://doi.org/10.1111/j.1768-322X.1999.tb01098.x.

Schwartz JC, Wang X, Podell ER, Cech TR. RNA seeds higher-order assembly of FUS protein. Cell Rep. 2013; 5(4):918–25. https://www.ncbi.nlm.nih.gov/pubmed/24268778, doi: 10.1016/j.celrep.2013.11.017.

Sepulveda G, Antkowiak M, Brust-Mascher I, Mahe K, Ou T, Castro NM, Christensen LN, Cheung L, Jiang X, Yoon D, Huang B, Jao LE. Co-translational protein targeting facilitates centrosomal recruitment of PCNT during centrosome maturation in vertebrates. Elife. 2018; 7. https://www.ncbi.nlm.nih.gov/pubmed/29708497, doi: 10.7554/eLife.34959.

Shcherbakova DM, Baloban M, Emelyanov AV, Brenowitz M, Guo P, Verkhusha VV. Bright monomeric near-infrared fluorescent proteins as tags and biosensors for multiscale imaging. Nat Commun. 2016; 7:12405. https://www.ncbi.nlm.nih.gov/pubmed/27539380, doi: 10.1038/ncomms12405.

Shin Y, Brangwynne CP. Liquid phase condensation in cell physiology and disease. Science. 2017; 357(6357). https://www.ncbi.nlm.nih.gov/pubmed/28935776, doi: 10.1126/science.aaf4382.

Shulga N, Goldfarb DS. Binding dynamics of structural nucleoporins govern nuclear pore complex permeability and may mediate channel gating. Mol Cell Biol. 2003; 23(2):534–42. https://www.ncbi.nlm.nih.gov/pubmed/12509452, doi: 10.1128/mcb.23.2.534-542.2003.

Smith J, Calidas D, Schmidt H, Lu T, Rasoloson D, Seydoux G. Spatial patterning of P granules by RNA-induced phase separation of the intrinsically-disordered protein MEG-3. Elife. 2016; 5. https://www.ncbi.nlm.nih.gov/pubmed/27914198, doi: 10.7554/eLife.21337.

So C, Seres KB, Steyer AM, Monnich E, Clift D, Pejkovska A, Mobius W, Schuh M. A liquid-like spindle domain promotes acentrosomal spindle assembly in mammalian oocytes. Science. 2019; 364(6447). https://www.ncbi.nlm.nih.gov/pubmed/31249032, doi: 10.1126/science.aat9557.

Sonnen KF, Schermelleh L, Leonhardt H, Nigg EA. 3D-structured illumination microscopy provides novel in-sight into architecture of human centrosomes. Biol Open. 2012; 1(10):965–76. http://www.ncbi.nlm.nih.gov/pubmed/23213374, doi: 10.1242/bio.20122337.

Sonnichsen B, Koski LB, Walsh A, Marschall P, Neumann B, Brehm M, Alleaume AM, Artelt J, Bettencourt P, Cassin E, Hewitson M, Holz C, Khan M, Lazik S, Martin C, Nitzsche B, Ruer M, Stamford J, Winzi M, Heinkel R, et al. Full-genome RNAi profiling of early embryogenesis in Caenorhabditis elegans. Nature. 2005; 434(7032):462–9. https://www.ncbi.nlm.nih.gov/pubmed/15791247, doi: 10.1038/nature03353.

Stearns T, Evans L, Kirschner M. Gamma-tubulin is a highly conserved component of the centrosome. Cell. 1991; 65(5):825–36. https://www.ncbi.nlm.nih.gov/pubmed/1840506, doi: 10.1016/0092-8674(91)90390-k.

Stearns T, Kirschner M. In vitro reconstitution of centrosome assembly and function: the central role of gamma-tubulin. Cell. 1994; 76(4):623–37. https://www.ncbi.nlm.nih.gov/pubmed/8124706, doi: 10.1016/0092-8674(94)90503-7.

Strom AR, Emelyanov AV, Mir M, Fyodorov DV, Darzacq X, Karpen GH. Phase separation drives heterochromatin domain formation. Nature. 2017; 547(7662):241–245. https://www.ncbi.nlm.nih.gov/pubmed/28636597, doi: 10.1038/nature22989.

Sundberg HA, Davis TN. A mutational analysis identifies three functional regions of the spindle pole component Spc110p in Saccharomyces cerevisiae. Mol Biol Cell. 1997; 8(12):2575–90. https://www.ncbi.nlm.nih.gov/pubmed/9398677, doi: 10.1091/mbc.8.12.2575.

Szappanos B, Suveges D, Nyitray L, Perczel A, Gaspari Z. Folded-unfolded cross-predictions and protein evolution: the case study of coiled-coils. FEBS Lett. 2010; 584(8):1623–7. https://www.ncbi.nlm.nih.gov/pubmed/20303956, doi: 10.1016/j.febslet.2010.03.026.

Takahashi M, Yamagiwa A, Nishimura T, Mukai H, Ono Y. Centrosomal proteins CG-NAP and kendrin provide microtubule nucleation sites by anchoring gamma-tubulin ring complex. Mol Biol Cell. 2002; 13(9):3235–45. https://www.ncbi.nlm.nih.gov/pubmed/12221128, doi: 10.1091/mbc.e02-02-0112.

Thomson AR, Wood CW, Burton AJ, Bartlett GJ, Sessions RB, Brady RL, Woolfson DN. Computational design of water-soluble alpha-helical barrels. Science. 2014; 346(6208):485–8. https://www.ncbi.nlm.nih.gov/pubmed/25342807, doi: 10.1126/science.1257452.

Tynan SH, Purohit A, Doxsey SJ, Vallee RB. Light intermediate chain 1 defines a functional subfraction of cyto-plasmic dynein which binds to pericentrin. J Biol Chem. 2000; 275(42):32763–8. https://www.ncbi.nlm.nih.gov/pubmed/10893222, doi: 10.1074/jbc.M001536200.

Updike DL, Hachey SJ, Kreher J, Strome S. P granules extend the nuclear pore complex environment in the C. elegans germ line. J Cell Biol. 2011; 192(6):939–48. https://www.ncbi.nlm.nih.gov/pubmed/21402789, doi: 10.1083/jcb.201010104.

Uversky VN, Gillespie JR, Fink AL. Why are “natively unfolded” proteins unstructured under physiologic conditions? Proteins. 2000; 41(3):415–27. https://www.ncbi.nlm.nih.gov/pubmed/11025552, doi: 10.1002/1097-0134(20001115)41:3<415::aid-prot130>3.0.co;2-7.

Vega IE, Umstead A, Kanaan NM. EFhd2 Affects Tau Liquid-Liquid Phase Separation. Front Neurosci. 2019; 13:845. https://www.ncbi.nlm.nih.gov/pubmed/31456657, doi: 10.3389/fnins.2019.00845.

Vorobjev IA, Chentsov Yu S. Centrioles in the cell cycle. I. Epithelial cells. J Cell Biol. 1982; 93(3):938–49. https://www.ncbi.nlm.nih.gov/pubmed/7119006, doi: 10.1083/jcb.93.3.938.

Wang J, Choi JM, Holehouse AS, Lee HO, Zhang X, Jahnel M, Maharana S, Lemaitre R, Pozniakovsky A, Drechsel D, Poser I, Pappu RV, Alberti S, Hyman AA. A Molecular Grammar Governing the Driving Forces for Phase Separation of Prion-like RNA Binding Proteins. Cell. 2018; 174(3):688–699 e16. https://www.ncbi.nlm.nih.gov/pubmed/29961577, doi: 10.1016/j.cell.2018.06.006.

Wang WJ, Soni RK, Uryu K, Tsou MF. The conversion of centrioles to centrosomes: essential coupling of duplication with segregation. J Cell Biol. 2011; 193(4):727–39. http://www.ncbi.nlm.nih.gov/pubmed/21576395, doi: 10.1083/jcb.201101109.

Watanabe S, Meitinger F, Shiau AK, Oegema K, Desai A. Centriole-independent mitotic spindle assembly relies on the PCNT-CDK5RAP2 pericentriolar matrix. J Cell Biol. 2020; 219(12). https://www.ncbi.nlm.nih.gov/pubmed/33170211, doi: 10.1083/jcb.202006010.

Waterhouse AM, Procter JB, Martin DM, Clamp M, Barton GJ. Jalview Version 2–a multiple sequence alignment editor and analysis workbench. Bioinformatics. 2009; 25(9):1189–91. https://www.ncbi.nlm.nih.gov/pubmed/19151095, doi: 10.1093/bioinformatics/btp033.

Wegmann S, Eftekharzadeh B, Tepper K, Zoltowska KM, Bennett RE, Dujardin S, Laskowski PR, MacKenzie D, Kamath T, Commins C, Vanderburg C, Roe AD, Fan Z, Molliex AM, Hernandez-Vega A, Muller D, Hyman AA, Mandelkow E, Taylor JP, Hyman BT. Tau protein liquid-liquid phase separation can initiate tau aggregation. EMBO J. 2018; 37(7). https://www.ncbi.nlm.nih.gov/pubmed/29472250, doi: 10.15252/embj.201798049.

Wippich F, Bodenmiller B, Trajkovska MG, Wanka S, Aebersold R, Pelkmans L. Dual specificity kinase DYRK3 couples stress granule condensation/dissolution to mTORC1 signaling. Cell. 2013; 152(4):791–805. https://www.ncbi.nlm.nih.gov/pubmed/23415227, doi: 10.1016/j.cell.2013.01.033.

Woodruff JB, Ferreira Gomes B, Widlund PO, Mahamid J, Honigmann A, Hyman AA. The Centrosome Is a Selective Condensate that Nucleates Microtubules by Concentrating Tubulin. Cell. 2017; 169(6):1066–1077 e10. https://www.ncbi.nlm.nih.gov/pubmed/28575670, doi: 10.1016/j.cell.2017.05.028.

Woodruff JB, Wueseke O, Hyman AA. Pericentriolar material structure and dynamics. Philos Trans R Soc Lond B Biol Sci. 2014; 369(1650). https://www.ncbi.nlm.nih.gov/pubmed/25047613, doi: 10.1098/rstb.2013.0459.

Woodruff JB, Wueseke O, Viscardi V, Mahamid J, Ochoa SD, Bunkenborg J, Widlund PO, Pozniakovsky A, Zanin E, Bahmanyar S, Zinke A, Hong SH, Decker M, Baumeister W, Andersen JS, Oegema K, Hyman AA. Regulated assembly of a supramolecular centrosome scaffold in vitro. Science. 2015; 348(6236):808–12. https://www.ncbi.nlm.nih.gov/pubmed/25977552, doi: 10.1126/science.aaa3923.

Wootton JC. Non-globular domains in protein sequences: automated segmentation using complexity measures. Comput Chem. 1994; 18(3):269–85. https://www.ncbi.nlm.nih.gov/pubmed/7952898, doi: 10.1016/0097-8485(94)85023-2.

Wueseke O, Zwicker D, Schwager A, Wong YL, Oegema K, Julicher F, Hyman AA, Woodruff JB. Polo-like kinase phosphorylation determines Caenorhabditis elegans centrosome size and density by biasing SPD-5 toward an assembly-competent conformation. Biol Open. 2016; 5(10):1431–1440. https://www.ncbi.nlm.nih.gov/pubmed/27591191, doi: 10.1242/bio.020990.

Zeng M, Shang Y, Araki Y, Guo T, Huganir RL, Zhang M. Phase Transition in Postsynaptic Densities Underlies Formation of Synaptic Complexes and Synaptic Plasticity. Cell. 2016; 166(5):1163–1175 e12. https://www.ncbi.nlm.nih.gov/pubmed/27565345, doi: 10.1016/j.cell.2016.07.008.

Zhang H, Elbaum-Garfinkle S, Langdon EM, Taylor N, Occhipinti P, Bridges AA, Brangwynne CP, Gladfelter AS. RNA Controls PolyQ Protein Phase Transitions. Mol Cell. 2015; 60(2):220–30. https://www.ncbi.nlm.nih.gov/pubmed/26474065, doi: 10.1016/j.molcel.2015.09.017.

Zhang JP, Li XL, Li GH, Chen W, Arakaki C, Botimer GD, Baylink D, Zhang L, Wen W, Fu YW, Xu J, Chun N, Yuan W, Cheng T, Zhang XB. Efficient precise knockin with a double cut HDR donor after CRISPR/Cas9-mediated double-stranded DNA cleavage. Genome Biol. 2017; 18(1):35. https://www.ncbi.nlm.nih.gov/pubmed/28219395, doi: 10.1186/s13059-017-1164-8.

Zhang Q, Huang H, Zhang L, Wu R, Chung CI, Zhang SQ, Torra J, Schepis A, Coughlin SR, Kornberg TB, Shu X. Visualizing Dynamics of Cell Signaling In Vivo with a Phase Separation-Based Kinase Reporter. Mol Cell. 2018; 69(2):347. https://www.ncbi.nlm.nih.gov/pubmed/29351851, doi: 10.1016/j.molcel.2018.01.008.

Zheng Y, Jung MK, Oakley BR. Gamma-tubulin is present in Drosophila melanogaster and Homo sapiens and is associated with the centrosome. Cell. 1991; 65(5):817–23. https://www.ncbi.nlm.nih.gov/pubmed/1904010, doi: 10.1016/0092-8674(91)90389-g.

Zheng Y, Wong ML, Alberts B, Mitchison T. Nucleation of microtubule assembly by a gamma-tubulin-containing ring complex. Nature. 1995; 378(6557):578–83. https://www.ncbi.nlm.nih.gov/pubmed/8524390, doi: 10.1038/378578a0.

Zimmerman WC, Sillibourne J, Rosa J, Doxsey SJ. Mitosis-specific anchoring of gamma tubulin complexes by pericentrin controls spindle organization and mitotic entry. Mol Biol Cell. 2004; 15(8):3642–57. https://www.ncbi.nlm.nih.gov/pubmed/15146056, doi: 10.1091/mbc.E03-11-0796.

Zwicker D, Decker M, Jaensch S, Hyman AA, Julicher F. Centrosomes are autocatalytic droplets of pericentriolar material organized by centrioles. Proc Natl Acad Sci U S A. 2014; 111(26):E2636–45. https://www.ncbi.nlm.nih.gov/pubmed/24979791, doi: 10.1073/pnas.1404855111.

