## Supplementary material for "Condensation of pericentrin proteins in human cells illuminates phase separation in centrosome assembly": Figure 4-source code 1

**Figure 4 source code 1:** Python code for analyzing condensate movement in Figure 4

*# Import libraries*

**import** **numpy** **as** **np**

**import** **pandas** **as** **pd**

**import** **os**

**import** **matplotlib.pyplot** **as** **plt**

os.chdir('/Users/jaolab/Desktop')

*# Import Data*

paths = [

'condensate_Results.csv',

'condensate_count.csv',

'centrosome_Results.csv',

'centrosome_count.csv',

'h2a_Results.csv',

'h2a_count.csv'

]

condensate_df = pd.read_csv(paths[0])

condensate_slice_count = pd.read_csv(paths[1], usecols=['Slice', 'Count'])

centrosome_df = pd.read_csv(paths[2])

centrosome_slice_count = pd.read_csv(paths[3], usecols=['Slice', 'Count'])

h2a_df = pd.read_csv(paths[4])

h2a_slice_count = pd.read_csv(paths[5], usecols=['Slice', 'Count'])

*# Explore Data*

condensate_df.columns = ['Condensate_index', 'Condensate_area', 'Condensate_XM', 'Condensate_YM']

centrosome_df.columns = ['Centrosome_index', 'Centrosome_area', 'Centrosome_XM', 'Centrosome_YM']

h2a_df.columns = ['H2A_index', 'H2A_area', 'H2A_XM', 'H2A_YM']

**print**(condensate_df.head())

**print**(condensate_slice_count.head())

*# Add slice number to dataframe*

*# Convert slice to string with a trailing space -- to work around double digits and tripe digits*

condensate_slice_str = list(map(**lambda** x: str(x) + ' ', condensate_slice_count['Slice']))

condensate_slice_index = []

**for** i, j **in** list(zip(condensate_slice_str, condensate_slice_count['Count'])): *# Repeat slice number by count number*

condensate_slice_index.extend((i*j).split())

condensate_slice_index_df = pd.DataFrame(condensate_slice_index, columns=['Slice'])

condensate_slice_df = pd.concat([condensate_df, condensate_slice_index_df], axis=1)

condensate_slice_df

*# Add slice to centrosome dataframe*

centrosome_slice_str = list(map(**lambda** x: str(x) + ' ', centrosome_slice_count['Slice']))

centrosome_slice_index = []

**for** i, j **in** list(zip(centrosome_slice_str, centrosome_slice_count['Count'])): *# Repeat slice number by count number*

centrosome_slice_index.extend((i*j).split())

centrosome_slice_index_df = pd.DataFrame(centrosome_slice_index, columns=['Slice'])

centrosome_slice_df = pd.concat([centrosome_df, centrosome_slice_index_df], axis=1)

centrosome_slice_df

*# Add slice to H2A dataframe*

h2a_slice_str = list(map(**lambda** x: str(x) + ' ', h2a_slice_count['Slice']))

h2a_slice_index = []

**for** i, j **in** list(zip(h2a_slice_str, h2a_slice_count['Count'])): *# Repeat slice number by count number*

h2a_slice_index.extend((i*j).split())

h2a_slice_index_df = pd.DataFrame(h2a_slice_index, columns=['Slice'])

h2a_slice_df = pd.concat([h2a_df, h2a_slice_index_df], axis=1)

h2a_slice_df.head()

*# Within each slice: Group condensate and centrosome to nearest H2A*

*# Calculate distance of all condensates to all H2A within the same slice*

slice_num = []

condensate_index_counter =[]

euclidean_dist = []

condensate_XM = []

condensate_YM = []

H2A_XM = []

H2A_YM = []

orig_H2A_index = []

**for** s **in** range(1, len(condensate_slice_count)+1): *# within each slice*

condensate_ind_counter = 0

**for** y_XM, y_YM **in** list(zip(condensate_slice_df.loc[condensate_slice_df['Slice']==str(s),'Condensate_XM'], condensate_slice_df.loc[condensate_slice_df['Slice']==str(s),'Condensate_YM'])): *# for each condensate*

condensate_ind_counter += 1

**for** h_XM, h_YM, ori_h2a_index **in** list(zip(h2a_slice_df.loc[h2a_slice_df['Slice']==str(s),'H2A_XM'], h2a_slice_df.loc[h2a_slice_df['Slice']==str(s),'H2A_YM'], h2a_slice_df.loc[h2a_slice_df['Slice']==str(s),'H2A_index'])): *# for each H2A*

*# calculate distance*

slice_num.append(s)

euclidean_dist.append(np.sqrt((y_XM - h_XM)**2 + (y_YM - h_YM)**2))

orig_H2A_index.append(ori_h2a_index)

condensate_index_counter.append(condensate_ind_counter)

condensate_XM.append(y_XM)

condensate_YM.append(y_YM)

H2A_XM.append(h_XM)

H2A_YM.append(h_YM)

condensate_h2a_per_slice = pd.DataFrame({'condensate_index_counter':condensate_index_counter,

'condensate_XM':condensate_XM, 'condensate_YM':condensate_YM,

'orig_H2A_index':orig_H2A_index,

'H2A_XM':H2A_XM, 'H2A_YM':H2A_YM,

'euclidean_dist':euclidean_dist, 'slice_num':slice_num})

condensate_h2a_per_slice

*# Calculate distance of all centrosomes to all H2A within the same slice*

slice_num = []

centrosome_index_counter =[]

euclidean_dist = []

centrosome_XM = []

centrosome_YM = []

H2A_XM = []

H2A_YM = []

orig_H2A_index = []

**for** s **in** range(1, len(centrosome_slice_count)+1): *# within each slice*

centrosome_ind_counter = 0

**for** y_XM, y_YM **in** list(zip(centrosome_slice_df.loc[centrosome_slice_df['Slice']==str(s),'Centrosome_XM'], centrosome_slice_df.loc[centrosome_slice_df['Slice']==str(s),'Centrosome_YM'])): *# for each centrosome*

centrosome_ind_counter += 1

**for** h_XM, h_YM, ori_h2a_index **in** list(zip(h2a_slice_df.loc[h2a_slice_df['Slice']==str(s),'H2A_XM'], h2a_slice_df.loc[h2a_slice_df['Slice']==str(s),'H2A_YM'], h2a_slice_df.loc[h2a_slice_df['Slice']==str(s),'H2A_index'])): *# for each H2A*

*# calculate distance*

slice_num.append(s)

euclidean_dist.append(np.sqrt((y_XM - h_XM)**2 + (y_YM - h_YM)**2))

orig_H2A_index.append(ori_h2a_index)

centrosome_index_counter.append(centrosome_ind_counter)

centrosome_XM.append(y_XM)

centrosome_YM.append(y_YM)

H2A_XM.append(h_XM)

H2A_YM.append(h_YM)

centrosome_h2a_per_slice = pd.DataFrame({'centrosome_index_counter':centrosome_index_counter,

'centrosome_XM':centrosome_XM, 'centrosome_YM':centrosome_YM,

'orig_H2A_index':orig_H2A_index,

'H2A_XM':H2A_XM, 'H2A_YM':H2A_YM,

'euclidean_dist':euclidean_dist, 'slice_num':slice_num})

centrosome_h2a_per_slice

*# Find the nearest H2A for every condensate and centrosome*

*""" Condensate """*

*# Get its index*

condensate_df_index = []

**for** slice_num **in** pd.unique(condensate_h2a_per_slice['slice_num']):

**for** condensate_index_counter **in** pd.unique(condensate_h2a_per_slice.loc[condensate_h2a_per_slice['slice_num']==slice_num, 'condensate_index_counter']):

*# Calculate minimum distance from a condensate to the condensates within a slice*

min_dist = np.min(condensate_h2a_per_slice.loc[(condensate_h2a_per_slice.slice_num==slice_num) & (condensate_h2a_per_slice.condensate_index_counter==condensate_index_counter) , 'euclidean_dist'])

*# find the dataframe index for this minimum distance*

condensate_df_index.append(condensate_h2a_per_slice.loc[(condensate_h2a_per_slice.slice_num==slice_num)&

(condensate_h2a_per_slice.condensate_index_counter==condensate_index_counter)&

(condensate_h2a_per_slice.euclidean_dist== min_dist), :].index)

*# Subset dataframe condensate_h2a_per_slice with condensate_df_index*

condensate_nearest_h2a = condensate_h2a_per_slice.iloc[np.array(condensate_df_index).flatten()]

*# Add original condensate index*

condensate_nearest_h2a.insert(0, 'orig_condensate_index', np.arange(1, len(condensate_slice_df)+1), False)

condensate_nearest_h2a

*# Insert condensate area*

condensate_nearest_h2a = condensate_nearest_h2a.reset_index(drop=True) *# reset index to properly insert values*

condensate_nearest_h2a.insert(4, 'condensate_area', condensate_slice_df.Condensate_area, True)

condensate_nearest_h2a

*""" Centrosome """*

centrosome_df_index = []

**for** slice_num **in** pd.unique(centrosome_h2a_per_slice['slice_num']):

**for** centrosome_index_counter **in** pd.unique(centrosome_h2a_per_slice.loc[centrosome_h2a_per_slice['slice_num']==slice_num, 'centrosome_index_counter']):

*# Calculate minimum distance from a centrosome to the centrosomes within a slice*

min_dist = np.min(centrosome_h2a_per_slice.loc[(centrosome_h2a_per_slice.slice_num==slice_num) & (centrosome_h2a_per_slice.centrosome_index_counter==centrosome_index_counter) , 'euclidean_dist'])

*# find the dataframe index for this minimum distance*

centrosome_df_index.append(centrosome_h2a_per_slice.loc[(centrosome_h2a_per_slice.slice_num==slice_num)&

(centrosome_h2a_per_slice.centrosome_index_counter==centrosome_index_counter)&

(centrosome_h2a_per_slice.euclidean_dist== min_dist), :].index)

*# Subset dataframe centrosome_h2a_per_slice with centrosome_df_index*

centrosome_nearest_h2a = centrosome_h2a_per_slice.iloc[np.array(centrosome_df_index).flatten()]

*# Add original centrosome index*

centrosome_nearest_h2a.insert(0, 'orig_centrosome_index', np.arange(1, len(centrosome_slice_df)+1), False)

centrosome_nearest_h2a

*# Remove cell/H2A with more than 2 centrosomes*

*# Get orig_H2A_index with no more than 2 centrosomes*

eligible_orig_H2A_index = centrosome_nearest_h2a.orig_H2A_index.value_counts().index[centrosome_nearest_h2a.orig_H2A_index.value_counts()<=2].tolist()

*# Subset condensate_nearest_h2a and centrosome_nearest_h2a with eligible_orig_H2A_index*

eligible_condensate_nearest_h2a = condensate_nearest_h2a[condensate_nearest_h2a.orig_H2A_index.isin(eligible_orig_H2A_index)]

eligible_centrosome_nearest_h2a = centrosome_nearest_h2a[centrosome_nearest_h2a.orig_H2A_index.isin(eligible_orig_H2A_index)]

eligible_centrosome_nearest_h2a

*# Calculate distance of condensate to centrosome within the same cell / with the same H2A index*

distance_condensate_to_centrosome = []

orig_h2a = []

H2A_X = []

H2A_Y = []

orig_condensate_index_list = []

orig_centrosome_index_list = []

condensate_XM_list = []

condensate_YM_list = []

centrosome_XM_list = []

centrosome_YM_list = []

condensate_area_list = []

slice_num = []

**for** elig_h2a **in** pd.unique(eligible_centrosome_nearest_h2a.orig_H2A_index): *# for every unique eligible H2A*

**for** condensate_XM, condensate_YM, orig_condensate_index, cond_area **in** list(zip(condensate_nearest_h2a.loc[condensate_nearest_h2a.orig_H2A_index == elig_h2a,'condensate_XM'],

condensate_nearest_h2a.loc[condensate_nearest_h2a.orig_H2A_index == elig_h2a,'condensate_YM'],

condensate_nearest_h2a.loc[condensate_nearest_h2a.orig_H2A_index == elig_h2a,'orig_condensate_index'],

condensate_nearest_h2a.loc[condensate_nearest_h2a.orig_H2A_index == elig_h2a,'condensate_area'])):

**for** centrosome_XM, centrosome_YM, orig_centrosome_index, s_num, h2a_x, h2a_y **in** list(zip(centrosome_nearest_h2a.loc[centrosome_nearest_h2a.orig_H2A_index == elig_h2a,'centrosome_XM'],

centrosome_nearest_h2a.loc[centrosome_nearest_h2a.orig_H2A_index == elig_h2a,'centrosome_YM'],

centrosome_nearest_h2a.loc[centrosome_nearest_h2a.orig_H2A_index == elig_h2a,'orig_centrosome_index'],

centrosome_nearest_h2a.loc[centrosome_nearest_h2a.orig_H2A_index == elig_h2a,'slice_num'],

centrosome_nearest_h2a.loc[centrosome_nearest_h2a.orig_H2A_index == elig_h2a,'H2A_XM'],

centrosome_nearest_h2a.loc[centrosome_nearest_h2a.orig_H2A_index == elig_h2a,'H2A_YM'])):

*# Calculate every condensate to every centrosome's distance with the same H2A*

orig_h2a.append(elig_h2a)

H2A_X.append(h2a_x)

H2A_Y.append(h2a_y)

condensate_area_list.append(cond_area)

slice_num.append(s_num)

orig_condensate_index_list.append(orig_condensate_index)

orig_centrosome_index_list.append(orig_centrosome_index)

distance_condensate_to_centrosome.append(np.sqrt((condensate_XM-centrosome_XM)**2 + (condensate_YM-centrosome_YM)**2))

condensate_XM_list.append(condensate_XM)

condensate_YM_list.append(condensate_YM)

centrosome_XM_list.append(centrosome_XM)

centrosome_YM_list.append(centrosome_YM)

condensate_to_centrosome_by_h2a = pd.DataFrame({'slice_num':slice_num, 'orig_condensate_index':orig_condensate_index_list,

'orig_centrosome_index':orig_centrosome_index_list,

'orig_h2a_index':orig_h2a, 'H2A_X':H2A_X, 'H2A_Y':H2A_Y,

'condensate_XM':condensate_XM_list,

'condensate_YM':condensate_YM_list, 'condensate_area':condensate_area_list,

'centrosome_XM':centrosome_XM_list, 'centrosome_YM':centrosome_YM_list,

'distance_condensate_to_centrosome':distance_condensate_to_centrosome})

condensate_to_centrosome_by_h2a

*# Use H2A for quantification*

*# Given starting_h2a_index, derive h2a_X, h2a_Y and starting_slice*

starting_h2a_index = 1053

end_slice = 97

h2a_slice_df.Slice = h2a_slice_df.Slice.astype('int32') *# Convert str to int for Slice column*

starting_XM = np.unique(h2a_slice_df.H2A_XM[h2a_slice_df.H2A_index==starting_h2a_index]).item()

starting_YM = np.unique(h2a_slice_df.H2A_YM[h2a_slice_df.H2A_index==starting_h2a_index]).item()

starting_slice = np.unique(h2a_slice_df.Slice[h2a_slice_df.H2A_index==starting_h2a_index]).item()

*# Find this h2a's index and its corresponding x, y from the next slice*

*# by choosing the closest coordination to the current coordination (handpicked_XM1, handpicked_YM1)*

*# Cache the new index and its corresponding x and y*

*# Repeat the procedure until the end_slice*

current_slice = starting_slice

candidate_X = starting_XM

candidate_Y = starting_YM

all_candidate_index = [starting_h2a_index]

all_candidate_Xs = [starting_XM]

all_candidate_Ys = [starting_YM]

all_slices = [starting_slice]

**while** current_slice < end_slice:

*# Get current H2A's coordinates*

current_h2a_X = candidate_X

current_h2a_Y = candidate_Y

next_slice = current_slice + 1

all_dist = []

all_index = []

all_x = []

all_y = []

*# Calculate distance of current H2A to all the H2As from the next slice*

**for** next_h2a_index, next_h2a_X, next_h2a_Y **in** list(zip(h2a_slice_df.H2A_index[h2a_slice_df.Slice==next_slice],

h2a_slice_df.H2A_XM[h2a_slice_df.Slice==next_slice],

h2a_slice_df.H2A_YM[h2a_slice_df.Slice==next_slice])):

all_dist.append(np.sqrt((current_h2a_X-next_h2a_X)**2+(current_h2a_Y-next_h2a_Y)**2))

all_index.append(next_h2a_index)

all_x.append(next_h2a_X)

all_y.append(next_h2a_Y)

*# Select candidate by the minimum dist and get its index & coordinates*

candidate_X = all_x[np.argmin(all_dist)]

candidate_Y = all_y[np.argmin(all_dist)]

candidate_index = all_index[np.argmin(all_dist)]

*# Cache the index & coordinates*

all_candidate_index.append(candidate_index)

all_candidate_Xs.append(candidate_X)

all_candidate_Ys.append(candidate_Y)

current_slice += 1

all_slices.append(current_slice)

cell_track_table = pd.DataFrame({'Slice_num':all_slices, 'H2A_index':all_candidate_index, 'H2A_X':all_candidate_Xs, 'H2A_Y':all_candidate_Ys})

*# We scrutinized through every slice to ensure our classification is correct, and modified incorrect data.*

*# ================ Below is SITUATIONAL =====================*

*# Remove/skip slices in which a cell is identified with more than 2 centrosomes.*

pd.set_option('display.max_rows', 10)

Handpicked_table = condensate_to_centrosome_by_h2a[(condensate_to_centrosome_by_h2a.orig_h2a_index.isin(cell_track_table.H2A_index))&(~condensate_to_centrosome_by_h2a.slice_num.isin([76,81,91]))] *# remove outliers*

*#Handpicked_table = condensate_to_centrosome_by_h2a[(condensate_to_centrosome_by_h2a.orig_h2a_index.isin(cell_track_table.H2A_index))]*

Handpicked_table

*# ================ Below is also SITUATIONAL =====================*

*# Exclude misclassified condensate index*

misclassified_condensate_index = [6917]

Handpicked_table = Handpicked_table[~Handpicked_table.orig_condensate_index.isin(misclassified_condensate_index)]

*# ================ Above is SITUATIONAL =====================*

*# Fill skipped slice with average of previous non-skipped slice and the next non-skipped slice*

*# We are only interested in slice_num, distance_condensate_to_centrosome, condensate_area*

handpicked_table_less = Handpicked_table[['slice_num', 'condensate_area', 'distance_condensate_to_centrosome']]

*# First, we find the skipped slices*

slices_stream = pd.Series(np.arange(starting_slice, end_slice+1))

skipped_slices = slices_stream[~slices_stream.isin(np.unique(Handpicked_table.slice_num))]

*# Filling average values of previous and next slides for each skipped slice*

**for** ss **in** skipped_slices:

**if** ss == starting_slice:

**print**('Select another starting_h2a_index and starting slice, current starting slice is skipped!')

**break**

*# Search previous non-skipped slice*

**for** searching_slice_backward **in** range(ss, starting_slice-1, -1):

**if** searching_slice_backward **in** skipped_slices.values:

**continue**

**else**:

prev_nonskipped_slide = handpicked_table_less[handpicked_table_less.slice_num==searching_slice_backward]

**break**

*# Search next non-skipped slice*

**for** searching_slice_forward **in** range(ss, end_slice+1):

**if** searching_slice_forward **in** skipped_slices.values:

**continue**

**else**:

next_nonskipped_slide = handpicked_table_less[handpicked_table_less.slice_num==searching_slice_forward]

**break**

*# Taking average of previous and next non-skipped slice to fill in skipped slices*

average_slice = pd.DataFrame(pd.concat([prev_nonskipped_slide,next_nonskipped_slide]).mean().values.reshape(1,3), columns=['slice_num', 'condensate_area', 'distance_condensate_to_centrosome'])

average_slice.slice_num = ss

handpicked_table_less = pd.concat([handpicked_table_less, average_slice])

handpicked_table_less = handpicked_table_less.sort_values('slice_num')

handpicked_table_less

*# GRAPHS*

*# Average Distance between Condensate and Centrosome by slice*

average_table = handpicked_table_less.groupby(['slice_num']).mean()

slices = pd.unique(handpicked_table_less.slice_num)

distance = average_table.distance_condensate_to_centrosome

plt.plot(slices,distance)

*# Average Condensate Area by slice*

area = average_table.condensate_area

plt.plot(slices, area)

*# Condensate count by slice*

count_gpby_slice = Handpicked_table.groupby('slice_num')['orig_condensate_index'].nunique()

count_gpby_non_skipped_slice = count_gpby_slice[~count_gpby_slice.index.isin(skipped_slices)]

skipped_slice_counts = []

*# Fill in skipped slices with average counts of previous and next nonskipped slices*

**for** s **in** skipped_slices:

**for** s_back **in** range(s, starting_slice-1, -1):

**if** s_back **in** skipped_slices.values:

**continue**

**else**:

prev_count = int(count_gpby_non_skipped_slice.values[count_gpby_non_skipped_slice.index==s_back])

**break**

**for** s_fore **in** range(s, end_slice+1):

**if** s_fore **in** skipped_slices.values:

**continue**

**else**:

next_count = int(count_gpby_non_skipped_slice.values[count_gpby_non_skipped_slice.index==s_fore])

**break**

*# Take average*

average_count = np.mean([prev_count, next_count])

skipped_slice_counts.append(average_count)

skipped_slice_count_df = pd.DataFrame({'slice_num':skipped_slices, 'counts':skipped_slice_counts})

count_gpby_slice_adjusted = count_gpby_non_skipped_slice.reset_index()[['slice_num','orig_condensate_index']]

count_gpby_slice_adjusted = count_gpby_slice_adjusted.rename(columns={'orig_condensate_index':'counts'})

count_gpby_slice_final = pd.concat([count_gpby_slice_adjusted, skipped_slice_count_df])

count_gpby_slice_final = count_gpby_slice_final.sort_values('slice_num')

count_gpby_slice_final

plt.plot(count_gpby_slice_final.slice_num, count_gpby_slice_final.counts)

*# Export*

Handpicked_table.to_csv('cell_track_table.csv') *# No skipped slices*

pd.DataFrame({'slices':slices, 'distance': distance}).to_csv('average_dist_vs_slice.csv') *# Skipped slices are filled*

pd.DataFrame({'slices':slices, 'area': area}).to_csv('average_area_vs_slice.csv') *# Skipped slices are filled*

count_gpby_slice_final.to_csv('condensate_counts_vs_slice.csv') *# Skipped slices are filled*
